# Functional dissection of *Drosophila Myc cis*-regulatory modules (Myc-CRMs) reveals developmentally active DNA-protein interactions

**DOI:** 10.64898/2026.07.02.736081

**Authors:** Jasmine Kharazmi, Thomas Brody, Cameron Moshfegh

## Abstract

Precise regulation of *Drosophila Myc* is essential for growth and homeostasis, yet regulation of its transcriptional control remains incompletely understood. We investigated the *Myc cis*-regulatory landscape using *in vivo* reporter assays, EMSA, and LC-MS/MS–based identification of DNA-associated proteins. By truncating *Myc cis*-regulatory modules (CRMs), we delineated the activity of conserved non-coding elements across adult female tissues and larval stages. Specific DNA-protein interactions were confirmed by EMSA using nuclear embryonic extracts. We developed a Solid Surface Magnetic Enrichment protocol (SSMEP) to pull down DNA-protein complexes formed on *Myc cis*-elements. Affinity purification followed by LC-MS/MS enabled the identification of candidate transcriptional regulators associated with Myc-CRMs. This integrative approach provides new insights into promoter structure and trans-regulatory architecture of *Myc* and their roles in developmental gene expression programs. Our study identifies a distal enhancer required for larval and pupal patterning, with activation dependent on specific spacing relative to the TATA-box core promoter, a strong enhancer cluster within 5′-UTR *cis*-elements active in ovaries and embryos, and a DPE-core promoter requiring nearby enhancer action. The DPE-linked enhancer can function with both Inr–DPE and TATA-box promoters.The identified Myc-CRMs interact with conserved signaling pathways to tightly control *Myc* during development.

**Graphical Abstract:** 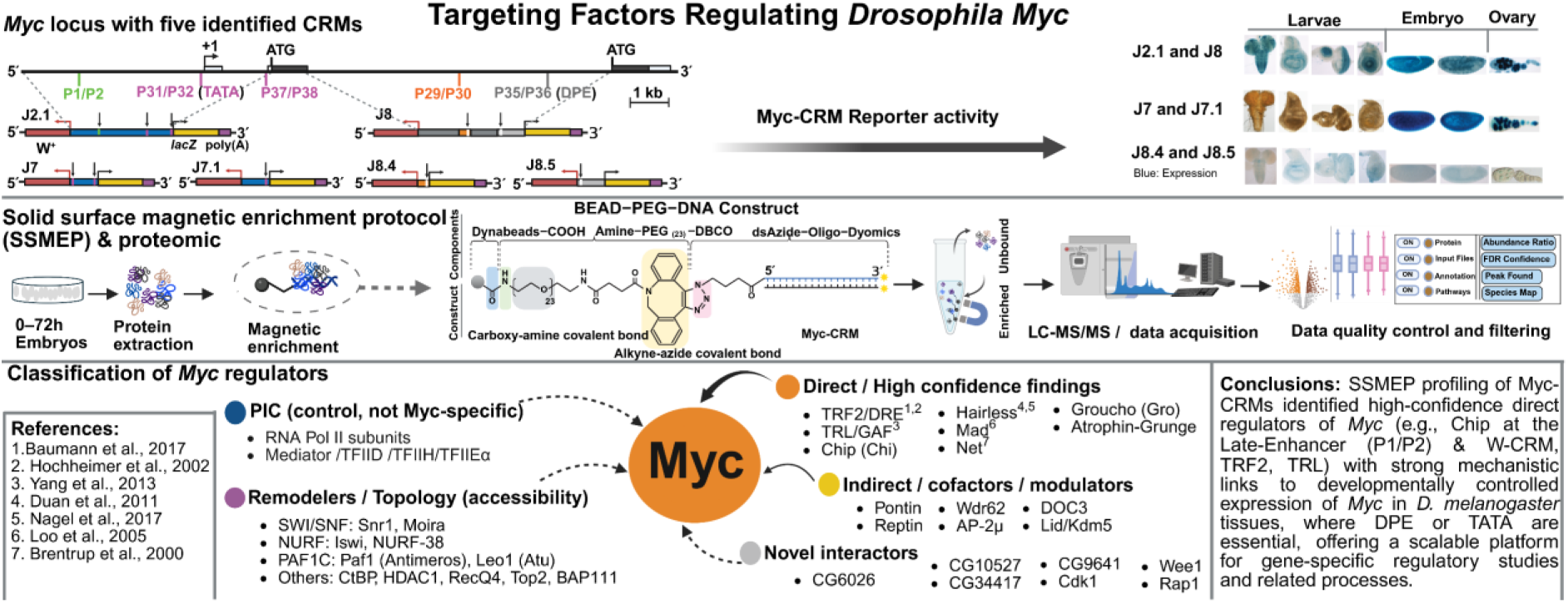

**Highlights:**

- *Myc* Oocyte Element: an eRNA-producing enhancer in ovaries and early embryos
- DPE promoter synergizes with nearby enhancer to drive *Drosophila Myc* transcription
- Late enhancer licenses TATA promoter for larval tissue-specific *Myc* transcription
- Promoter-enhancer dynamics differentially drive *Myc* during development
- SSMEP protocol helps purify and enrich low-abundance DNA-protein complexes

## 1. Introduction

Developmental patterning is tightly controlled by interactions between transcription factors and *cis*-regulatory elements such as enhancers and promoters, which together encode the logic of spatial and temporal gene expression (Davidson and Erwin, 2006), coordinating cell growth, proliferation, apoptosis, and differentiation throughout development. Such tightly orchestrated regulation is especially critical during early embryogenesis, when precise gene activity specifies cell fates and organ primordia to ensure normal body size and organ function (Davidson and Erwin, 2006). Among the transcriptional regulators governing these processes, *MYC* functions as an evolutionarily conserved and potent proto-oncogene whose activity requires tight regulation at both transcriptional and post-transcriptional levels during development and adulthood (Gallant et al., 1996). MYC/MAX heterodimers bind to E-box sequences to activate target genes that drive S-phase cyclin expression, DNA replication, and cell growth and proliferation (Carroll et al., 2018).

Conversely, when MYC binds to other bHLHzip factors such as Miz1, it activates genes involved in cell cycle arrest (Fioresi et al., 2022), thereby countering its proliferative effects. MAD/MAX and MNT/MAX heterodimers also compete with MYC/MAX for the same E-box sites, repressing MYC/MAX-driven transcriptional activation (Grandori et al., 2000). Beyond these core regulatory roles, *Drosophila Myc*, like its human ortholog *MYC*, contributes to regenerative growth, proliferation, and stem cell maintenance. For example, Myc cooperates with Wingless signaling to promote tissue regeneration after injury or apoptosis in imaginal discs (Smith-Bolton et al., 2009). In terminally differentiated organs, somatic stem cells (SSCs) support tissue renewal after damage, and MYC family proteins are similarly essential for regeneration following injury or cancer therapy. They regulate processes such as midgut and neuroblast maintenance, neural stem cell proliferation, and epidermal regeneration (Samuels et al., 2020).

Because deregulated *MYC* expression is linked to numerous diseases, from neurodegeneration to cancer and diabetes, precise *MYC* control is essential for maintaining physiological homeostasis throughout development and adult life. The evolutionary conservation and functional similarity between *Drosophila Myc* and human *c-MYC* (Gallant et al., 1996) make the fruit fly, with its short life cycle and genetic tractability, an ideal model for studying Myc biology. *Myc* expression is highly responsive to developmental and mitogenic signaling pathways including Dpp, Wingless, Notch, Hippo, and Ecdysone (Johnston et al., 1999).

These signaling inputs can positively or negatively regulate Myc levels, maintaining the balance between cell growth, proliferation, and differentiation. Despite extensive research, the mechanisms transmitting these divers signals to control *Myc* transcription remain incompletely understood.

Here, we identified multiple *cis*-regulatory modules as conserved sequence blocks within the *Myc* gene, along with the two major transcription start sites, a TATA core promoter at the proximal 5′-end and a downstream promoter element (DPE) within the second intron (Kharazmi et al., 2012). We further analyzed truncated *Myc-lacZ* transgenes encompassing these promoters and conserved regulatory regions. Our results confirm that dynamic expression of *Myc* during oogenesis and early embryogenesis arises from transcription initiation at multiple sites, including a newly identified start site at the 3′-end of intron 1 within the proximal 5′-UTR, which contains a strong developmental enhancer within the Myc-CRM P37/P38 (ChrX:3,374,908..3,374,960). We also found that high expression levels in larval discs and brain require additional late or differentiation enhancers beyond those sufficient for embryonic and ovarian expression. DNA-protein interaction analyses identified key regulatory factors binding *Myc cis*-elements, primarily cell cycle regulators and components of signaling pathway including Wingless, Notch, Dpp, Hedgehog, MAPK, JAK/STAT, insulin-like receptor (InR), and Hippo. Together, these findings define a multilayered regulatory architecture integrating developmental and metabolic signaling inputs at the *Myc* locus and provide a framework and dataset to facilitate future mechanistic dissection of *Myc* transcriptional control during development.

## 2. Materials and Methods

### 2.1. Generation of lacZ reporter fly strains

Previously described constructs (pJ2.1, pJ7, pJ8, pJ8.4, pJ8.5) and intermediate vectors were used as templates. The J7 promoter region (ChrX:3,373,200..3,375,005) includes 100 bp upstream of the TSS, noncoding exon 1, intron 1, and noncoding sequences of exon 2. Variants J7.1–J7.5 contained truncated or composite promoter/enhancer fragments, as summarized in Table S1a. All transgenes were sequenced using proximal and distal primers (Table S1b). Detailed information on molecular cloning and reporter plasmid construction is provided in Methods S1.

### 2.2. Drosophila stocks and X-Gal staining

Transgenic reporter lines were generated using random P-element–mediated transformation or ɸC31 integrase-mediated site-specific integration (Kharazmi et al., 2012; Kharazmi and Moshfegh, 2013). For constructs pJ7.1– pJ7.5, transgenesis was performed in y[1] w[1118] embryos (Rainbow Transgenic Flies Inc., CA), yielding lines J7.1–J7.5. Multiple independent insertion lines were established for each construct to control for position effects associated with random integration, and a subset of lines was analyzed for *lacZ* expression. Reporter expression patterns were reproducible across most of independent insertion lines examined for each construct. For selected fly strains (e.g., J8.4 and J8.5), expression patterns were also consistent with those previously reported (Kharazmi et al., 2012), further supporting the robustness of the observed reporter activity. *dpp-lacZ* and *w[1118]* served as positive and negative controls (Table S1c). Flies were maintained on standard medium at 25°C. X-Gal staining was performed on homozygous F2 tissues using modified protocols (Methods S2).

### 2.3. Protein extraction from embryos

Wild-type *Canton-S* embryos were collected 0–12 h AEL, for up to 72 h, stored at 4°C, dechorionated, and suspended in cold buffer I. Homogenization, centrifugation, and fractionation yielded soluble cytoplasmic (SCF) and nuclear (SNF) extracts, which were frozen at −80°C. Protein concentrations were determined by BCA assay, and quality confirmed by SDS-PAGE and Coomassie staining. Detailed information on the extraction protocols are provided in Methods S3.

### 2.4. EMSA oligonucleotides design and assays

EMSA oligonucleotides were designed from conserved Myc-CRMs. A dTCF/Pan W-CRM served as positive control. A 45 bp sequence from the noncoding region of the mouse Mdm2 gene (C57BL/6J; NC_000076.6:117,686,685–117,712,961, gi|372099100|) was randomized using a sequence shuffling tool to generate a scrambled oligonucleotide (Scm) serving as a negative control (Table S1d).

Oligos were labelled with IR-Dye Dyomics 781 and Azide-NHS ester (Microsynth AG, Switzerland) and annealed as described (Tables S1d, S1e, S1g). EMSA reactions contained 200 µg SCF or SNF protein, binding buffer, and labelled oligonucleotides, incubated for 30 min in the dark. Native TBE-PAGE (5%) was used for separation, and gels were imaged on an Odyssey Classic Infrared Imager.

For proteomic analysis, the same CRM-derived and control oligonucleotides were used as affinity baits to identify associated nuclear proteins.

Detailed oligo end-labelling, cycling protocols for annealing, and EMSA assays are provided in Methods S4 and S5.

### 2.5. Magnetic Bead–PEG_(23)_–DNA construction and protein enrichment

The DBCO–PEG_(23)_–coated magnetic beads (prepared via EDC-mediated amide coupling and stored at 10 μg/μL in storage buffer) were used for copper-free click chemistry attachment of azide-modified dsDNA oligonucleotide: 140 μL beads were reacted with 7 μL dsDNA (0.45 pmol/μL) in 300 μL storage buffer for 4.5 h at room temperature (dark, 7 rpm). The resulting Bead–PEG_(23)_–DNA construct was washed 4x (10 min each) in storage buffer, resuspended in 500 μL storage buffer, and stored at 4°C (<2 days).

For protein enrichment, Bead–PEG_(23)_–DNA (140 μL, 10 μg/μL) was magnetically separated, supernatant removed, rinsed once with storage buffer, and resuspended at 10 μg/μL. Binding reactions (100 μL total) contained: 140 μg beads, 10 μL 10x binding buffer (Odyssey® Infrared EMSA Kit), 10 μL 25 mM DTT/2.5% Tween-20, 5 μL 1 μg/μL poly(dI·dC), 5 μL 1% NP-40, 5 μL 100 mM MgCl₂, 200 μg protein (SCF for cytoplasmic or SNF for nuclear samples), and water. Mixtures were incubated for 30 min in the dark with occasional gentle mixing. Bead–PEG_(23)_–DNA/protein complexes (e.g., IR-Pos1/Pos2:SNF; Table S1f) were washed four times with storage buffer, residual liquid removed, and stored at –80°C until further analysis. Full coupling and binding protocols are in Methods S6 and S7.

### 2.6. Sample processing for LC-MS/MS analysis

Beads were denatured in 8 M urea, reduced with DTT, alkylated with iodoacetamide, and digested with trypsin. Peptides were desalted, dissolved in 0.1% formic acid, and separated by reverse-phase chromatography (2 cm trap and 15 cm analytical C18 column) using a 5–35% acetonitrile gradient at 250 nL/min. Mass spectrometry was performed on a Thermo Scientific Orbitrap Eclipse Tribrid mass spectrometer coupled to an EASY-nLC 1200 system. Detailed digestion and LC-MS/MS conditions are provided in Methods S8.

### 2.7. Proteomic data processing and analysis

Spectrum matches were searched against the *Drosophila melanogaster* reference proteome, supplemented with a contaminant database. Standard search settings were applied (including trypsin specificity where applicable), and identifications were filtered to a false discovery rate (FDR) <1% at peptide and protein levels. Protein quantification was based on unique peptides, and abundance values were normalized across samples within Proteome Discoverer prior to downstream analysis.

Filtered and normalized data were imported into ComplexBrowser (Michalak et al., 2019; https://computproteomics.bmb.sdu.dk/app_direct/ComplexBrowser/) for quality control and protein complex analysis. QC metrics included generation of normalized log2 abundance boxplots, coefficient of variation (CV) histograms, and pairwise scatter plots with Pearson correlation coefficients to assess reproducibility between samples. Analyses were performed across all sample pairs, including negative control (Scm), experimental conditions (Myc-CRMs), and positive control samples. ComplexBrowser analyses were performed using curated protein complex databases (CORUM and Complex Portal), enabling evaluation of complex-level abundance changes across experimental conditions.

Differential protein abundance analysis was performed using VolcaNoseR (Goedhart and Luijsterburg, 2020; https://huygens.science.uva.nl/VolcaNoseR2/; plotting log2 abundance ratios versus −log10(q-value)). Volcano plots were generated from log2 abundance ratios and corresponding q-values. Significance thresholds were set at q < 0.01 and an absolute abundance ratio > 2.

*Drosophila* relevant protein complexes were identified by integrating ComplexBrowser results with manual curation using UniProt, FlyBase, STRING, and The Interactive Fly, supplemented with literature review. This combined workflow, ComplexBrowser outputs and curated annotations, was used to identify and interpret relevant protein complexes from the MS datasets showing consistent changes across conditions. Detailed software parameters, quality control metrics, and implementation-specific settings are provided in Methods S9.

## 3. Results

### 3.1. Reporter assays: functional dissection of Myc cis-regulatory modules

To establish the functional relevance of *Myc cis*-regulatory modules (Myc-CRMs), we previously analyzed the reporter activity of J2.1 and J7 transgenes. J2.1, containing 7.2 kb upstream sequences, drives *Myc* expression in larval and adult female tissues, whereas the smallest J7 (1.928 kb; ChrX:3,373,008..3,375,005) can express *lacZ* in embryos and ovaries (Fig. 1A–B; Table S1a).

**Fig. 1.**
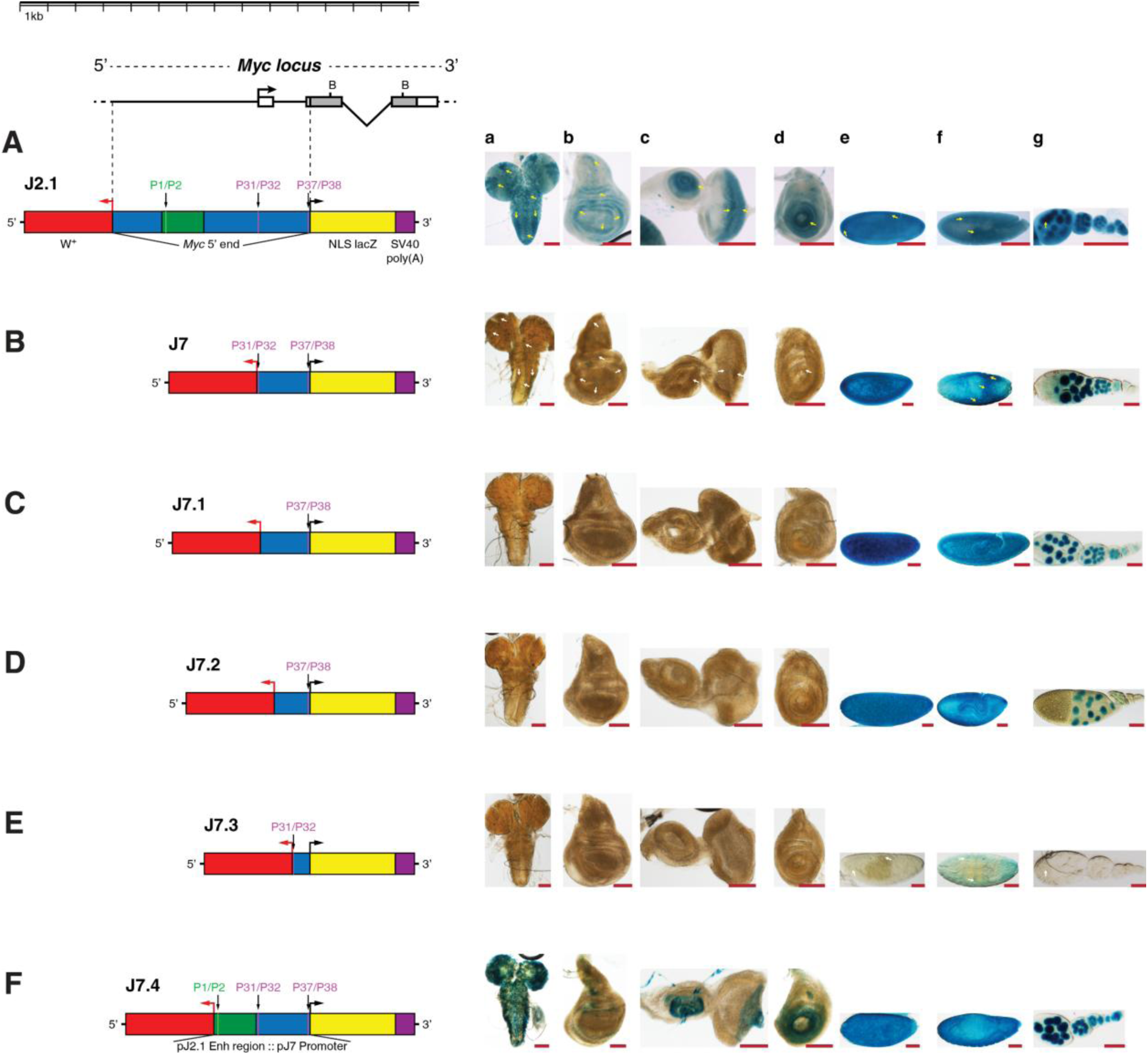
Partitioning *Myc* 5′ regulatory sequences identifies a developmental Late-Enhancer and an Oocyte Element. (A) J2.1 contains 7.2 kb of *Myc* 5′-end noncoding sequences (blue/green), including the upstream Late-Enhancer (P1/P2), the TATA-CorePromoter (P31/P32), and the proximal Oocyte Element (P37/P38). (B) J7 (1.923 kb) is a truncation lacking P1/P2. (C–E) J7-derived truncations: J7.1–J7.3 retain or remove P37/P38. (F) J7.4 fuses the P1/P2 enhancer with J7 sequences. Panels (a–g) show *lacZ* reporter activity in larval brain and imaginal discs, embryos, and adult ovaries. Construct sequences and coordinates of Myc-CRMs are provided in Tables S1a, S1d, and S1g. β-Galactosidase staining was performed overnight at 25°C on 3rd instar larval tissues and adult female ovaries/embryos. Scale bar: 100 µm. **Abbreviations:** a, brain; b, wing disc; c, eye-antennal disc; d, leg disc; e–f, embryos; g, ovary.

Truncation analysis of J7-derived constructs in this study revealed that the 53 bp P37/P38 element is necessary and sufficient for expression in embryos and ovarian tissues, independent of the TATA promoter (Fig. 1C–E; J7.1–J7.3). Based on its strong activity in these tissues, we here designate the 53 bp P37/P38 Myc-CRM (ChrX:3,374,908..3,374,960; Table S1d and S1g) the *Myc* Oocyte Element.

Fusion of the J2.1 enhancer region containing the P1/P2 element to the J7 promoter (J7.4, Fig. 1F) restored reporter activity in larval brain, imaginal discs, embryos, and ovaries—with weaker expression in eye and wing discs compared to J2.1—indicating that the P1/P2 enhancer drives *Myc* expression in larval tissues and the proper distance between the enhancer and promoter is required for strong transcriptional activation. These results in this study highlight that the P1/P2 enhancer functions as a late developmental enhancer, active during the onset of differentiation in larval limb, eye, wing, and brain patterning. Details on the expression patterns of reporter constructs J7.1–J7.4 are described in Results S3.1.

Previous analysis of the 8 kb full-length intron 2 sequences within the *Myc* locus identified a dense upstream regulatory region enriched for multiple *cis*-regulatory modules (CRMs), including the P29/P30 enhancer cluster and a downstream Inr–DPE core promoter that function together to mimic endogenous *Myc* expression in larval imaginal discs, the brain, and adult female tissues (Fig. 2A). The two truncations J8.4 and J8.5 showed that neither of the regions alone is sufficient to express *lacZ* in the tested tissues (Fig. 2B–C).

**Fig. 2.**
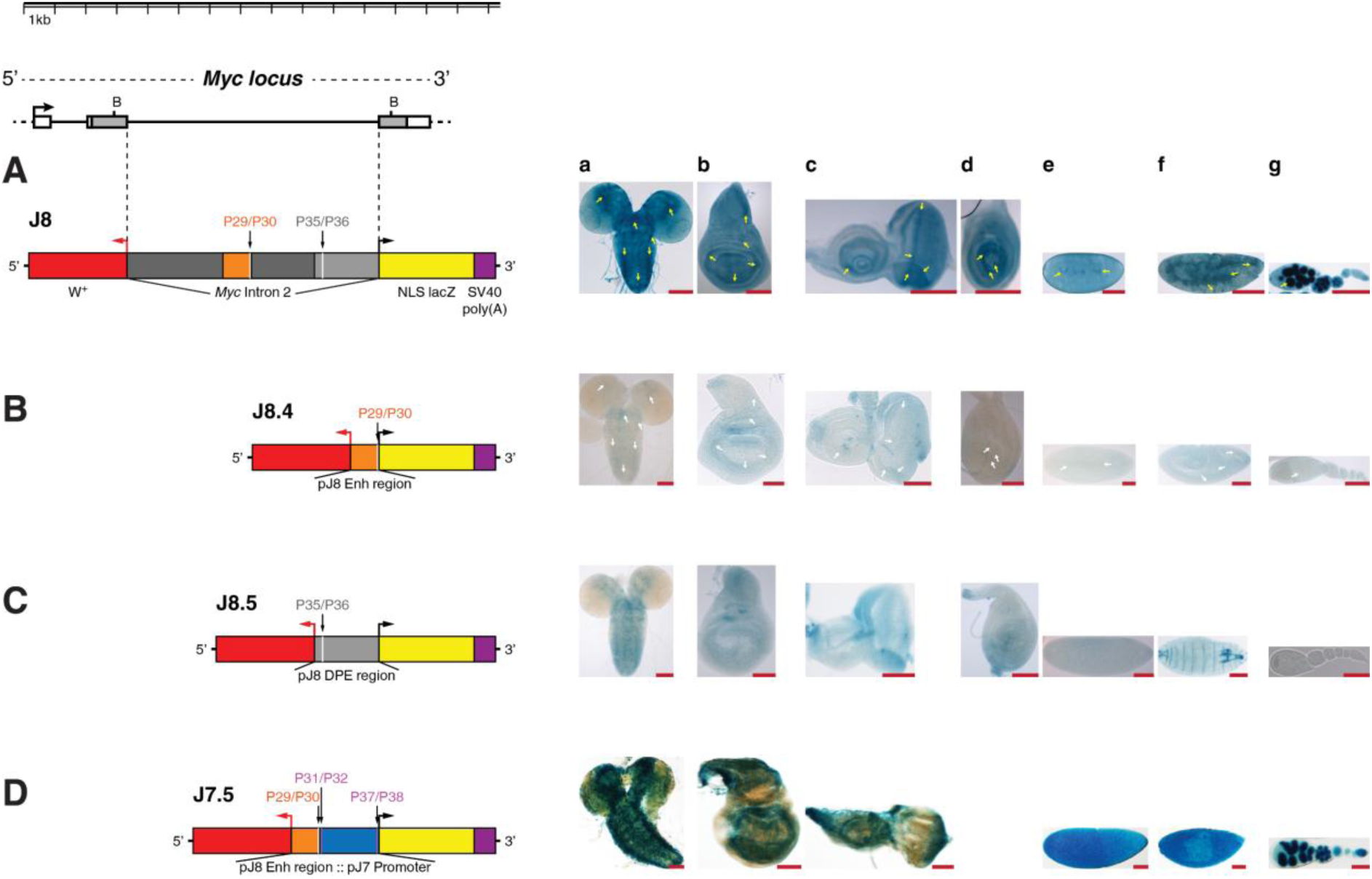
The DPE-CorePromoter requires the DPE-Enhancer for transcriptional activation. Reporters J8, J8.4, and J8.5 were adapted from Kharazmi et al. (2012). (A) J8 comprises the full 8 kb intron 2 sequences (dark gray, orange, white, light gray) and contains both the DPE-CorePromoter (P35/P36) and DPE-Enhancer (P29/P30). (B–C) Truncated constructs J8.4 and J8.5 carry only enhancer or promoter sequences, respectively. (D) J7.5 fuses the DPE-Enhancer upstream of the J7 promoter. J8 drives *lacZ* expression in larval brain and discs (a-d), embryos (e-f) and ovary (g) reflecting endogenous *Myc* mRNA (yellow arrowheads), whereas J8.4 and J8.5 fail to reproduce this pattern (white arrowheads). The composite J7.5 construct restores expression in larval and adult female tissues. Construct sequences and coordinates of Myc-CRMs are provided in Tables S1a, S1d, and S1g. β-Gal staining was performed overnight at 25°C on 3^rd^ instar larval and adult female tissues. Scale bar: 100 µm. **Abbreviations**: a, brain; b, wing disc; c, eye-antennal disc; d, leg disc; e-f, embryos; g, ovary.

In this study, we further isolated one conserved CRM from this region (P29/P30), selected based on sequence features suggestive of transcription factor binding, to test whether it is sufficient to recapitulate J8-like *Myc* expression and whether its activity depends on an Inr–DPE core promoter. Fusion of P29/P30 to the J7 TATA-containing core promoter in a composite construct J7.5 restored reporter activity (Fig. 2D), demonstrating that this CRM is sufficient to drive *Myc* expression and can function with either Inr–DPE or TATA promoters. Details on the creation of J7.5 construct (Methods S1) and the expression patterns of J7.5, J8, J8.4, and J8.5 are described in Results S3.1.

### 3.2 Myc-CRMs as regulatory DNA baits with scrambled oligonucleotide and W-CRM controls

Building on prior activities in reporter assays in embryonic, ovarian, and larval tissues and in silico analyses, we selected five oligonucleotides from the noncoding regions of the *Myc* locus, corresponding to functional Myc-CRMs as targets to isolate DNA–protein complexes. To identify proteins associated with these regulatory DNA elements, we performed affinity purification–mass spectrometry using CRM-derived oligonucleotides as baits and embryonic nuclear extracts as the protein source. A scrambled oligonucleotide from the mouse noncoding region of *Mdm2* gene served as a negative control, and a well-characterized dTCF/Pangolin binding site (W-CRM) was used as a positive control (Fig. S3; Methods, Section 2.4; Table S1d and S1f). The W-CRM contains an HMG-box with a basic tail and a C-clamp motif, analogous to mammalian LEF1 site, and recruits nuclear dTCF/Pangolin upon Wingless signaling.

Oligonucleotide annealing and EMSA assays confirmed probe integrity and binding capacity (Fig. 3, A–L). DNA– protein complexes from Myc-CRMs and controls were then subjected to LC-MS/MS to identify associated regulatory factors. See Results S3.2 for extensive quality control assays of oligonucleotides and protein extracts.

**Fig. 3.**
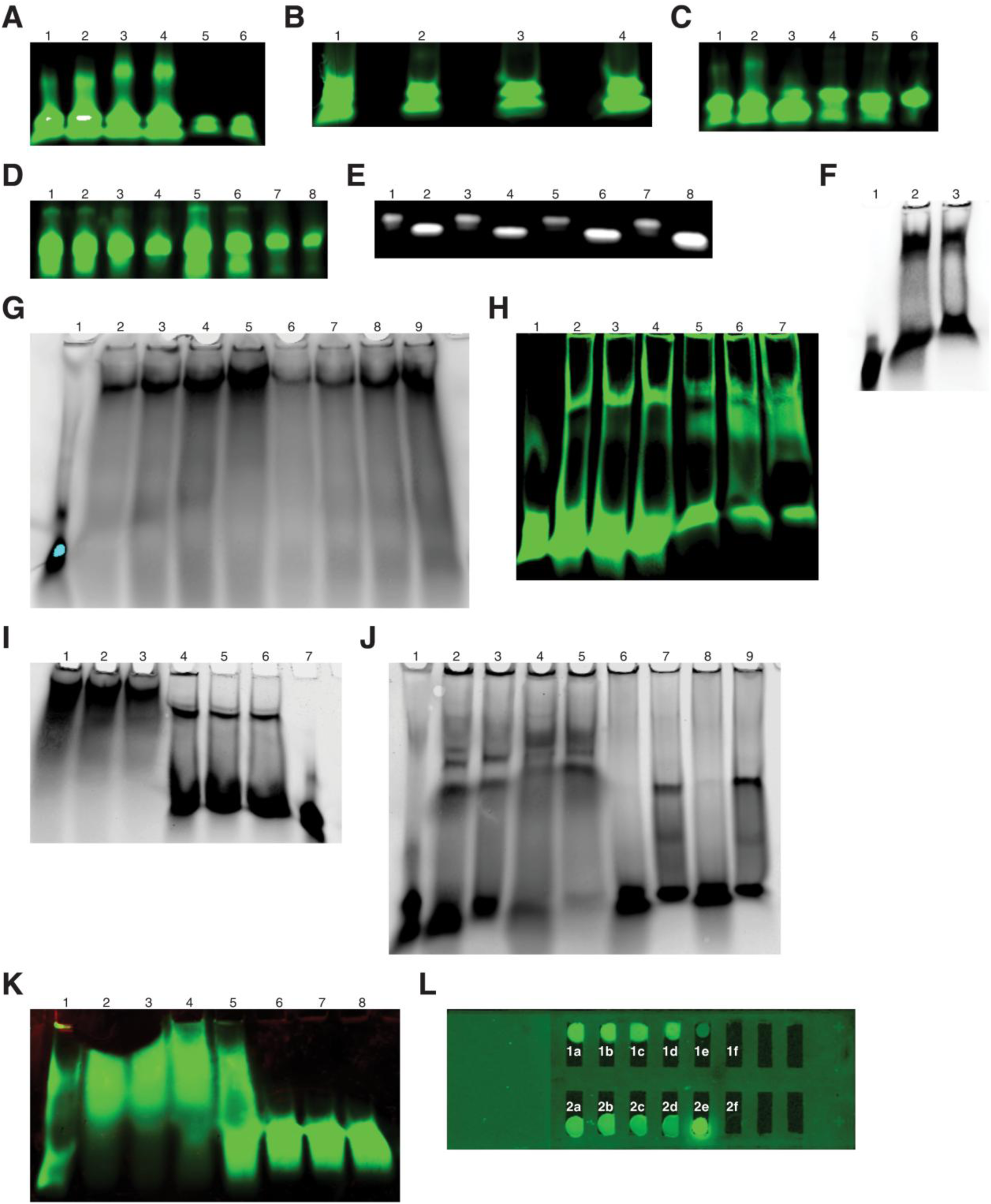
Annealing of oligonucleotides and EMSA assays with *Drosophila* embryonic extracts. **(A)** Annealing of IR-labeled positive control oligonucleotides. Lanes 1-4:azide-labeled; 5-6: unlabeled. Amounts loaded: 1, 3, 5 = 0.9 pmol; 2, 4, 6 = 1.35 pmol. **(B)** Annealing of IR- and azide-labeled Myc-CRM oligos (1.35 pmol/lane): 1 = P31/P32; 2 = P32/P31; 3 = P37/P38; 4 = P38/P37. **(C)** Annealing of IR/azide-labeled oligos (1.35 pmol/lane): 1 = Pos1/Pos2; 2 = pos2/Pos1; 3 = Scm1/Scm2; 4 = Scm2/Scm1; 5 = P1/P2; 6 = P2/P1. **(D)** Annealing and pegylation check for Myc-CRMs (0.45 pmol lanes 1,3,5,7; 0.225 pmol lanes 2,4,6,8). Oligos: 1-2 = P31/P32; 3-4 = P32/P31; 5-6 = P35/P36; 7-8 = P36/P35. **(E)** Migration comparison of unpegylated (lanes 2,4,6,8) vs. PEG_(23)_–labeled positive control DNA (0.2 pmol in lanes 1,5; 0.4 pmol in lanes 2,4,6; 0.3 pmol in lanes 3,7; 0.6 pmol in lane 8). **(F)** EMSA comparing SNF binding to pegylated vs. non-pegylated positive control DNA (0.4 pmol/lanes: 1 = free DNA; 2-3 = DNA + 100 µg SNF). **(G-I)** EMSA titrations showing SCF and SNF binding to: (**G**) positive control (2.25 pmol/lane input) 1 = free DNA; 2–5 = 100, 200, 300, 400 µg SCF; 6–9 = SNF; **(H)** Myc-CRM P31/P32 (1.35 pmol) 1 = free DNA; 2-4 = 70,100,150 µg SCF; 5-7 = SNF); **(I)** Myc-CRM P37/P38 (0.45 pmol/lane) 1-3 = 500,400,300 µg SCF; 4-6 = SNF; 7 = free DNA. **(J)** EMSA specificity: SCF and SNF binding to Myc-CRM P1/P2 vs. Scm negative control (1.35 pmol/lane); Scm1/Scm2 (1 = free DNA; 2,4 = 70,150 µg SCF; 6,8 = SNF); P1/P2 (3,5 = 70,150 µg SCF; 7,9 = SNF). **(K)** Competition EMSA: IR-labeled P1/P2 probe (1.35 pmol) with increasing unlabeled P3/P4 competitor (1x–60x molar excess) and 130 µg SNF (**1** = free DNA; **2** = P1/P2 + SNF; **3** = 1:1 molar excess; **4** = 1:5; **5** = 1:15; **6** = 1:30; **7** = 1:45; **8** = 1:60).**(L)** Visualization of IR-labeled Bead–PEG₍₂₃₎–DNA constructs (3 µL, from 0.3 pmol/300 µL) before protein binding. Panels **1a–1d**: labeled Bead–PEG₍₂₃₎–P37/P38 and P38/P37; **1e**: raw beads; **1f**: protein binding buffer only: Panels **2a-2d**: supernatants from Bead–PEG₍₂₃₎ preps; **2e**: free P37/P38 (0.3 pmol); **2f**: beads only supernatant. All reactions were run on 5% non-denaturing TBE PAGE gel (Bio-Rad): 40 V for 2 h (Panel E) or 100 V for 55 min (others).

**Fig. 4.**
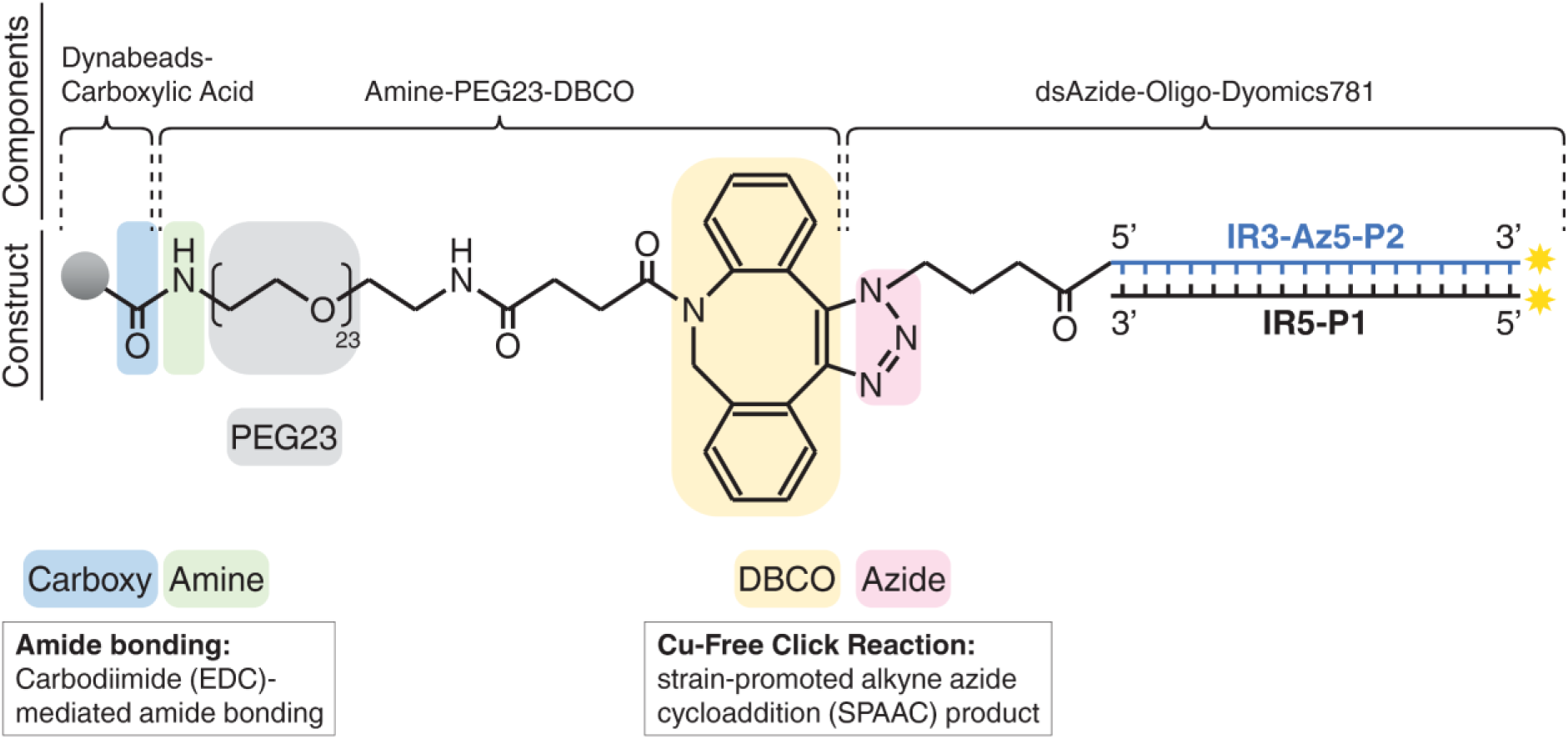
Construct for Solid Surface Magnetic Enrichment Protocol (SSMEP). Amine–PEG_(23)_–DBCO was covalently linked to carbodiimide-activated Dynabeads® MyOne™ Carboxylic Acid beads via amide bond formation between Dibenzocyclooctyne (DBCO) and N-hydorxysuccinimide (NHS) ester Azide. The resulting chimeric structure underwent copper-free click chemistry (Jewett and Bertozzi, 2010; Eeftens et al., 2015) between the DBCO– alkyne and the azide-modified oligonucleotide, producing the Bead–PEG₍₂₃₎–Oligonucleotide construct. This construct was used for biomagnetic separation and affinity purification of *Myc* regulators from *Drosophila* embryonic nuclear extracts (see Methods, section 2.6).

### 3.3. Solid Surface Magnetic Enrichment Protocol (SSMEP) for enrichment of proteins and identification by mass spectrometry

To enable purification of DNA-bound protein complexes for mass spectrometry, we developed a solid-surface magnetic enrichment protocol (SSMEP). Standard approaches often fail to reduce background noise and achieve sufficient sensitivity and selectivity for the effective purification of low-abundance proteins from whole cell lysates, limiting the quality of downstream mass spectrometry (MS) analyses (Goodman et al., 2018). Methods that immobilize and enrich nucleic acid-protein or protein-protein complexes on a solid surface can substantially decrease background noise of chimeric spectra, thereby lowering the false discovery rate (FDR) and enabling higher-confidence identification of low-abundance target proteins (Leitner et al., 2010). Building on these principles, we developed and applied a solid-surface magnetic enrichment protocol (SSMEP) that immobilizes protein complexes bound to DNA on carboxylated magnetic Bead–PEG_(23)_–DNA constructs (Fig. 5; Methods 2.6; Methods S6) and recovers them by biomagnetic separation from crude embryonic nuclear extracts of *Drosophila.* The soluble cytoplasmic fraction (SCF) showed non-specific binding to the negative control (Scm; Fig. 3J, lanes 2 and 4). Therefore, only the transcription factor–enriched soluble nuclear fraction (SNF) was used for protein identification. This approach identified *Myc*-associated transcription factors, coregulators, and protein complexes implicated in *Myc* regulatory networks at multiple levels and can be adapted to other protein–RNA/microRNA or antigen–antibody interaction studies. Proteins identified by mass spectrometry are discussed in subsequent sections, while a detailed workflow for the enrichment of *Myc* regulators and assay conditions is provided in Results S3.3.

**Fig. 5.**
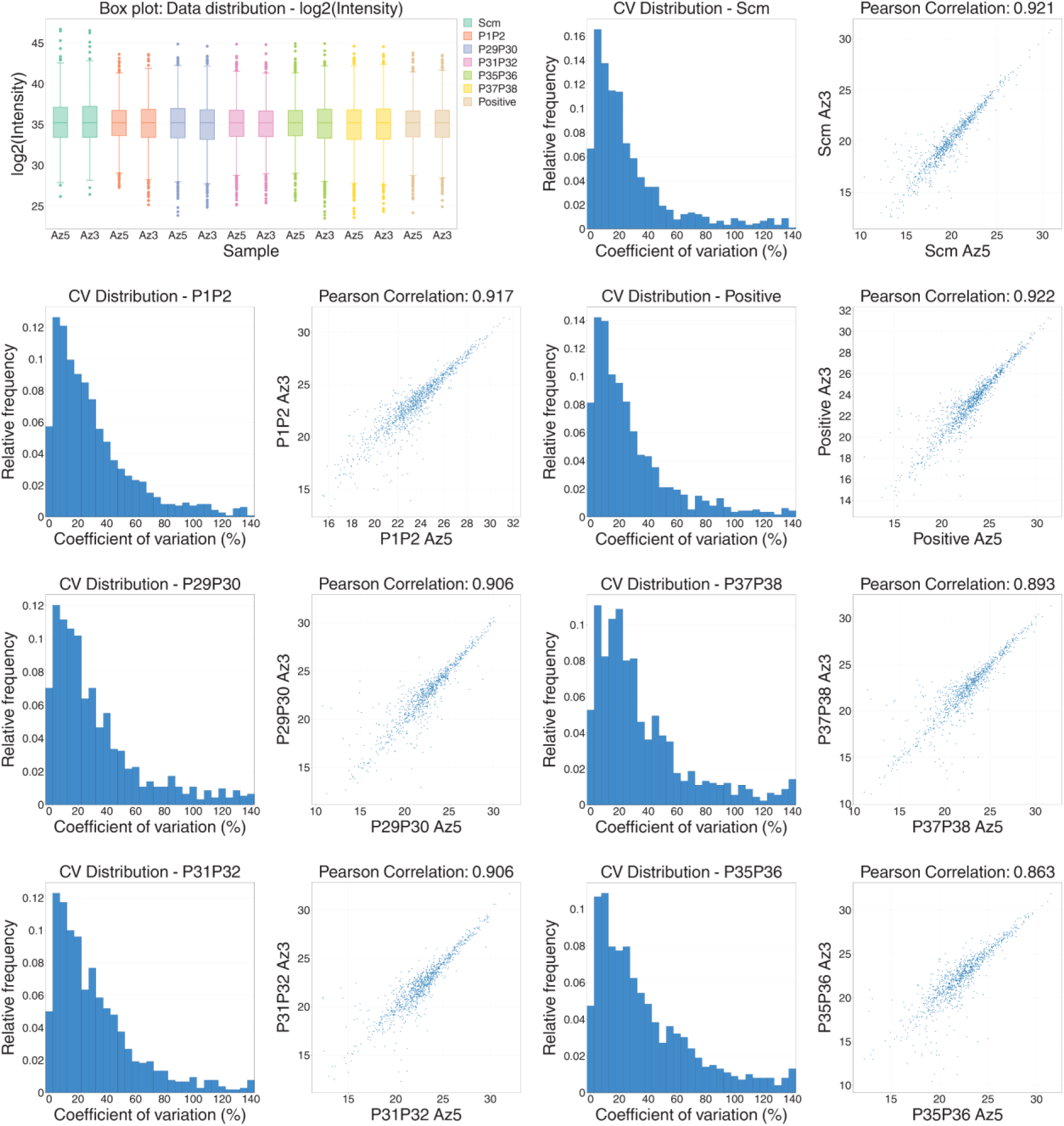
Quality control analysis of label-free proteomic data using ComplexBrowser. ComplexBrowser was used to visualize data quality and the overall structure of replicate and sample relationships. Box plots show log2-transformed intensities across duplicate samples for the negative control (Scm), Myc-CRMs, and positive control. Histograms display coefficient of variation (CV) distributions for replicates within each condition (Scm, Positive, P1P2, P29P30, P35P36, P37P38). Scatter plots compare replicates’ log2 intensities, with Pearson correlation coefficients indicated. Az5 and Az3 denote 5′- or 3′-end azide modifications of the oligos. Azide (N-hydroxysuccinimide ester) moieties were removed during protein purification prior to LC-MS/MS analysis.

### 3.4. Mass spectrometry data quality, reproducibility, and enrichment analysis

LC-MS/MS datasets from raw magnetic beads, embryonic nuclear extract (SNF), and DNA-bound protein complexes were processed, with raw bead datasets used exclusively to assess potential contaminants. Downstream analysis of MS output data from control and Myc-CRM samples was performed using Proteome Discoverer™ 2.5 and ComplexBrowser (Results S3.4; Supplementary Data S2: Fig. S7 and Table S2; Methods, Sections 2.6–2.7). Sample replicates, abundance ratios relative to the negative control (Scm), and applied filters are provided in Supplementary Data S2–S7: Figs. S7–S12 and Tables S2–S7; Supplementary Data S9–S11: Tables S9a–S9h, S10a–S10g, and S11a–Table S11f.

Proteome Discoverer–generated datasets (after curation, filter application, and contaminant removal) from two replicates per sample group were analyzed using ComplexBrowser to assess data quality and reproducibility (Fig. 5). Each sample included two replicates (5 Myc-CRM targets, plus positive and negative controls), resulting in 14 datasets. ComplexBrowser requires ungrouped replicates as input data; therefore, each replicate was submitted separately without prior averaging. Analyses were then performed on replicate pairs for each sample (Supplementary Data S5: Fig. S10 and Table S5). Low coefficients of variation and strong correlations confirmed dataset reproducibility (Fig. 5). Volcano plots comparing Myc-CRM targets to controls highlight statistically enriched proteins relevant to *Myc* regulation (q < 0.01; absolute abundance ratio > 2.0; Figs. 6–8). The complete lists of proteins (UniProt accessions) underlying the volcano plots are provided in Supplementary Tables S13– S18.

**Fig. 6.**
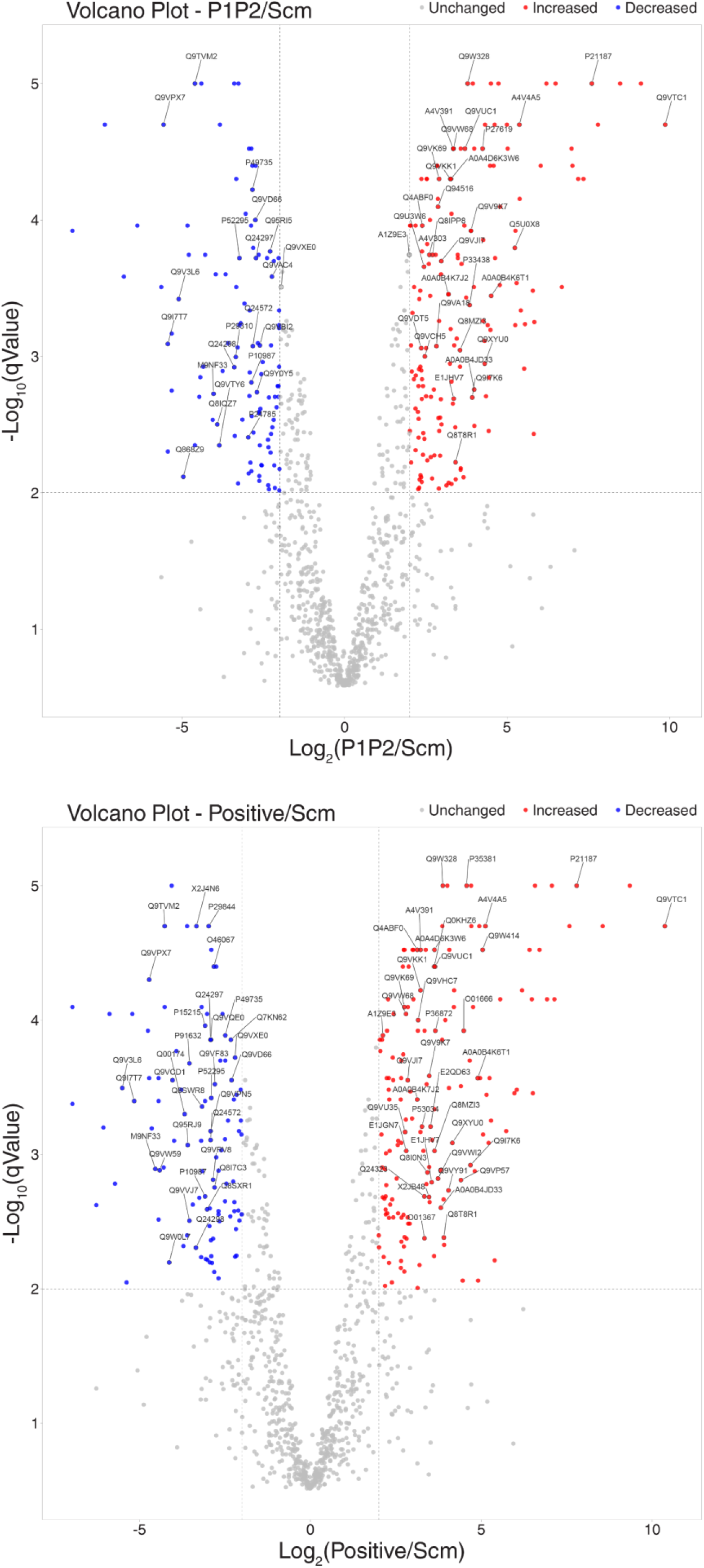
Volcano plots comparing Myc-CRM P1/P2 and the positive control to the negative control. Volcano plots show log2(abundance ratio) vs. –log10(q-value) for P1P2 or positive samples (heat-map pairs, see Figs. S3–S4) compared to the Scm negative control. Significance thresholds are marked at q < 0.01 (–log10(q) > 2) and absolute abundance ratio > 2.0.

**Fig. 7.**
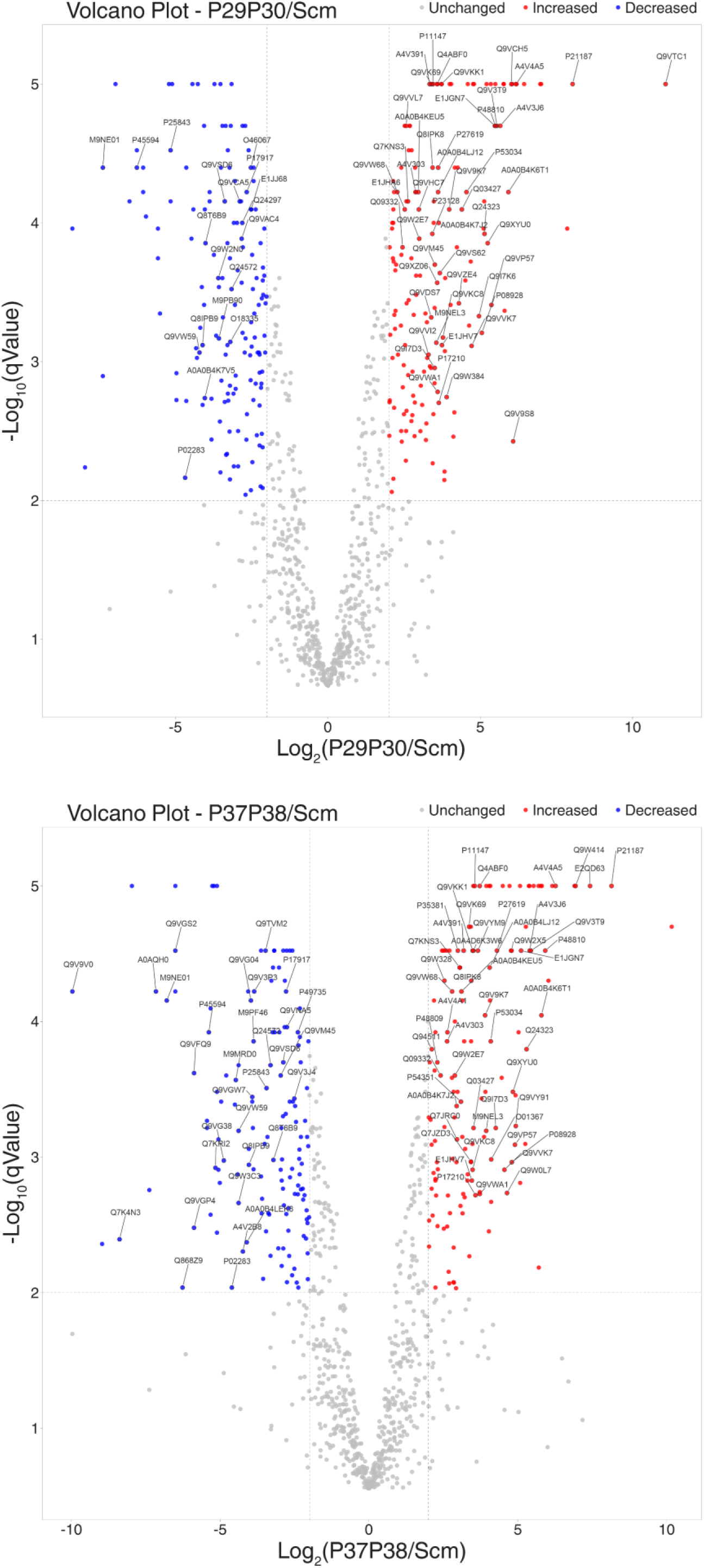
Volcano plots comparing Myc-CRM P29/P30 and P37/P38 to negative control. Volcano plots show log2(abundance ratio) vs. –log10(q-value) for P29/P30 or P37/P38 samples (heat-map pairs) compared to the Scm negative control. Significance thresholds are set at q < 0.01 (–log10(q) > 2) and absolute abundance ratio > 2.0.

**Fig. 8.**
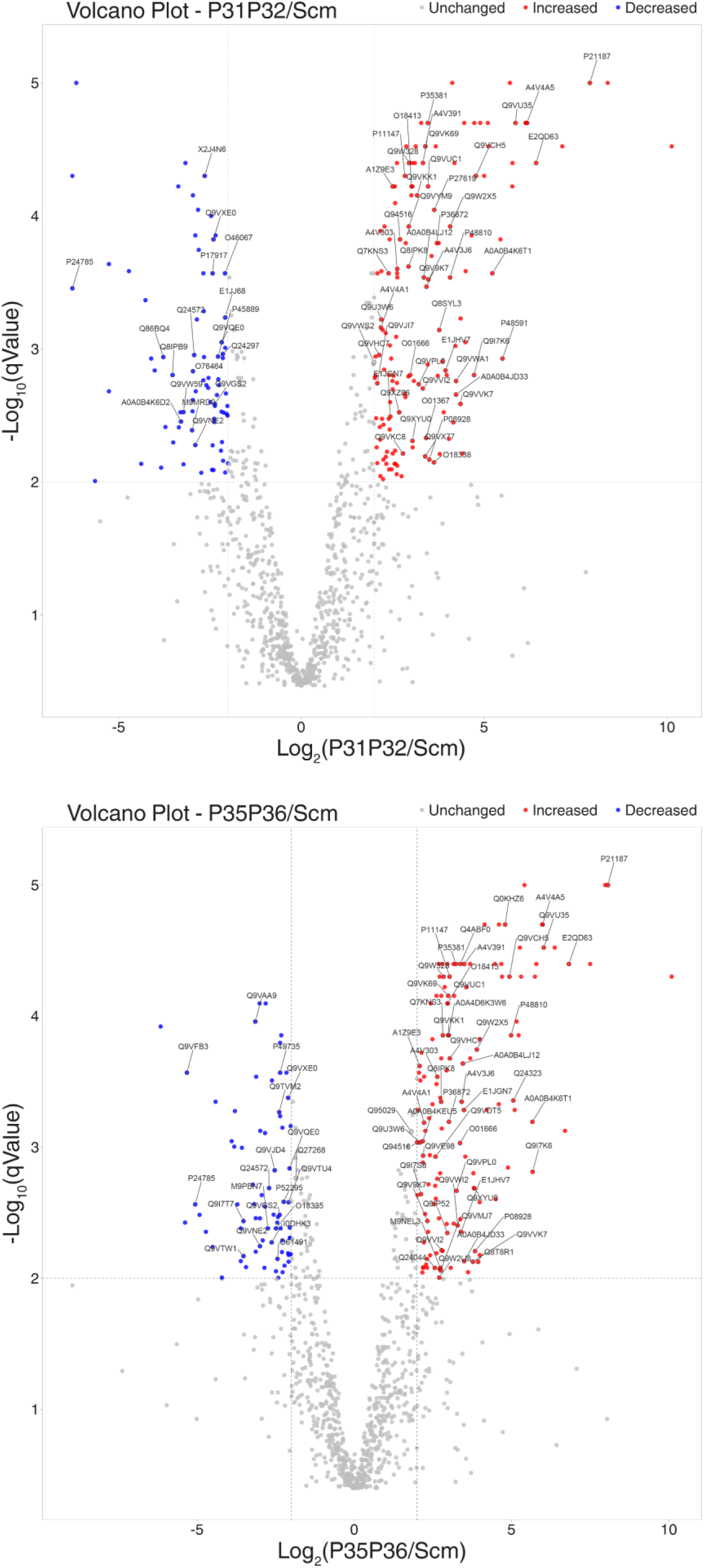
Volcano plots comparing Myc-CRM P31/P32 and P35/P36 to the negative control. Volcano plots show log2(abundance ratio) vs. –log10(q-value) for P31/P32 or P35/P36 samples (heat-map pairs) compared to the Scm negative control. Significance thresholds are set at q < 0.01 (–log10(q) > 2) and absolute abundance ratio > 2.0.

### 3.5. Heat-map analysis of proteins enriched at Myc targets and controls

The proteome dataset of grouped replicate (2,264 proteins; Supplementary Data S6: Fig. S11 and Table S6) was filtered in Proteome Discoverer 2.5 for high-confidence hits (protein FDR ‘high’, >10-fold enrichment over scrambled control (Scm), low Scm peak confidence, *D. melanogaster*–specific annotation), yielding 1,001 CRM-associated candidates (Supplementary Data S7: Fig. S12 and Table S7). These were visualized as heat-maps (Figs. S3–S4) to assess clustering of functionally related protein groups for downstream analyses. Further details are in Results S3.5.

### 3.6. Functional classification of proteins associated with the positive control W-CRM and Myc-CRMs

High-confidence proteins associated with the positive control W-CRM and each Myc-CRM were categorized by biological activity and molecular function (Fig. 9; Supplementary Data S9: Tables S9a–S9h). Identification of known regulators—including components of Wnt/Wingless (Wg), Hippo, Dpp, Hedgehog, and Notch pathways— validated our enrichment approach, particularly for the W-CRM positive control. Remaining classifications, proportions (e.g., cell signaling 25–30%, posttranscriptional regulators 30–39%), unique/shared factors, and sample details (**A–F**) are provided in Fig. 9 legend, Tables S9a–S9h, and Results S3.6.

**Fig. 9.**
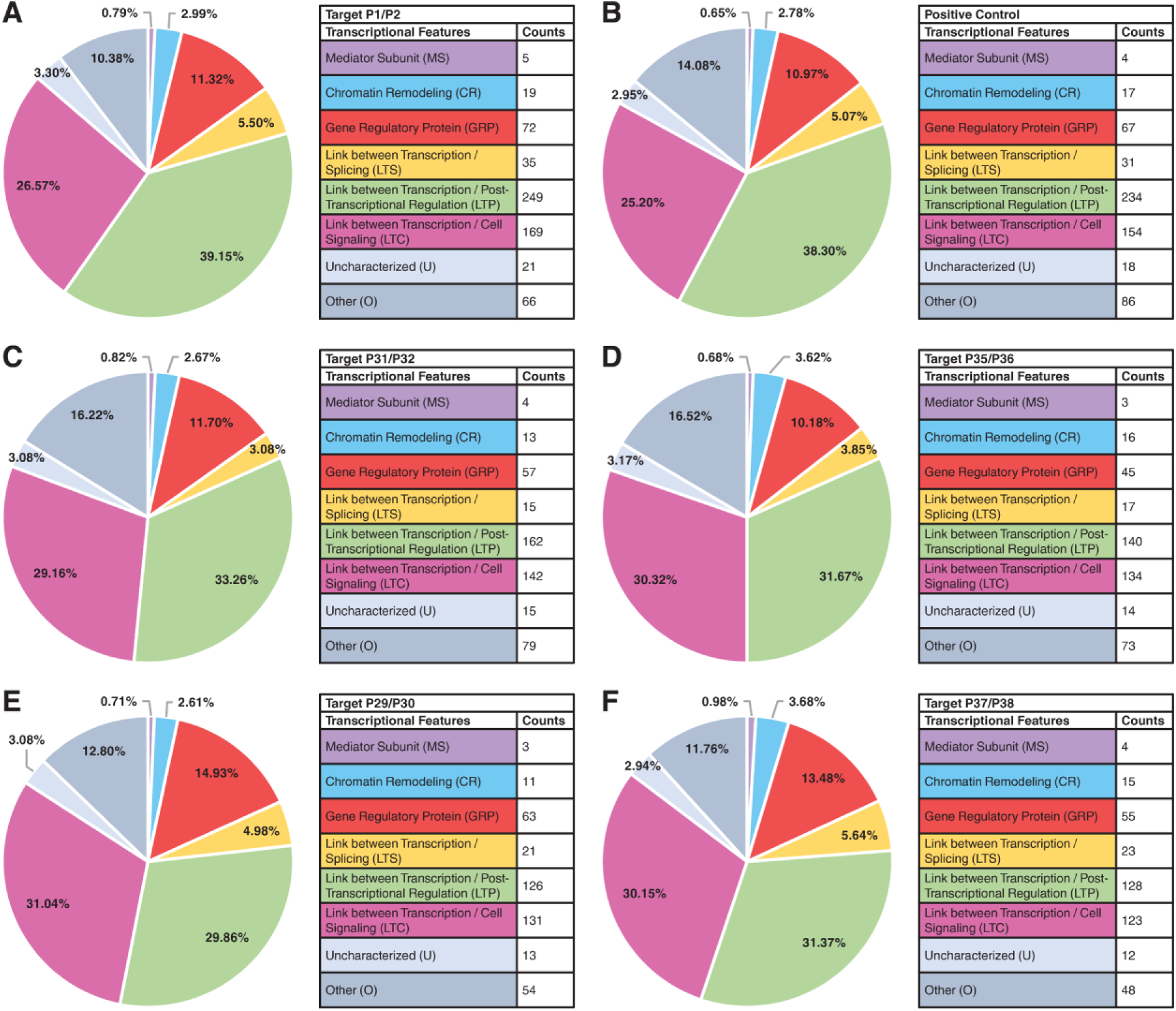
Functional classification of factors associated with Myc-CRMs and positive control. Pairs of samples identified as functionally similar by heat-map analysis (Figs. S3–S4) of high-confidence proteins (Table S7; Fig. S12) are positioned adjacent to each other. Among the 1,001 identified factors, approximate proportions were: mediator complex 0.65–0.98%; chromatin remodeling 2.61–3.68%; gene-specific proteins 10.18–14.93%; splicing machinery 3.08–5.64%; posttranscriptional regulators 29.86–39.15%; cell signaling 25.20–30.32%; uncharacterized 2.94–3.30%; and other categories 10.38–16.52%. Samples: (**A**) Late-Enhancer (P1/P2); (**B**) positive control (W-CRM); (**C**) TATA-CorePromoter (P31/P32); (**D**) DPE-CorePromoter (P35/P36); (**E**) DPE-Enhancer (P29/P30); (**F**) Oocyte Element (P37/P38). See Tables S9a–S9h for detailed pie chart data.

### 3.7. Validation of SSMEP using W-CRM: recovery of transcriptional and chromatin regulatory complexes

The TCF/Pangolin DNA recognition motif containing an HMG-helper site is known to respond to Wingless signaling in a tissue-dependent manner (Archbold et al., 2014), making the W-CRM (Fig. S5) a suitable positive control for the proteomic workflow applied to *Myc* regulatory elements.

In addition to the components of basal transcription machinery (Pol II complex, TFIID, Mediator) and chromatin remodeling factors, including NURD (Simjang), NURF (ISWI, E(bx)), SWI/SNF (Brahma) (Snrl, Moira), and PaflC (Atu/Leo1, Antimeros/Paf1) the positive control W-CRM validates SSMEP sensitivity for known Wnt/Wg-associated factors and reveals functional clustering with the *Myc* Late-Enhancer (P1/P2) (Figs. 9-10; Figs. S3-4).

**Fig. 10.**
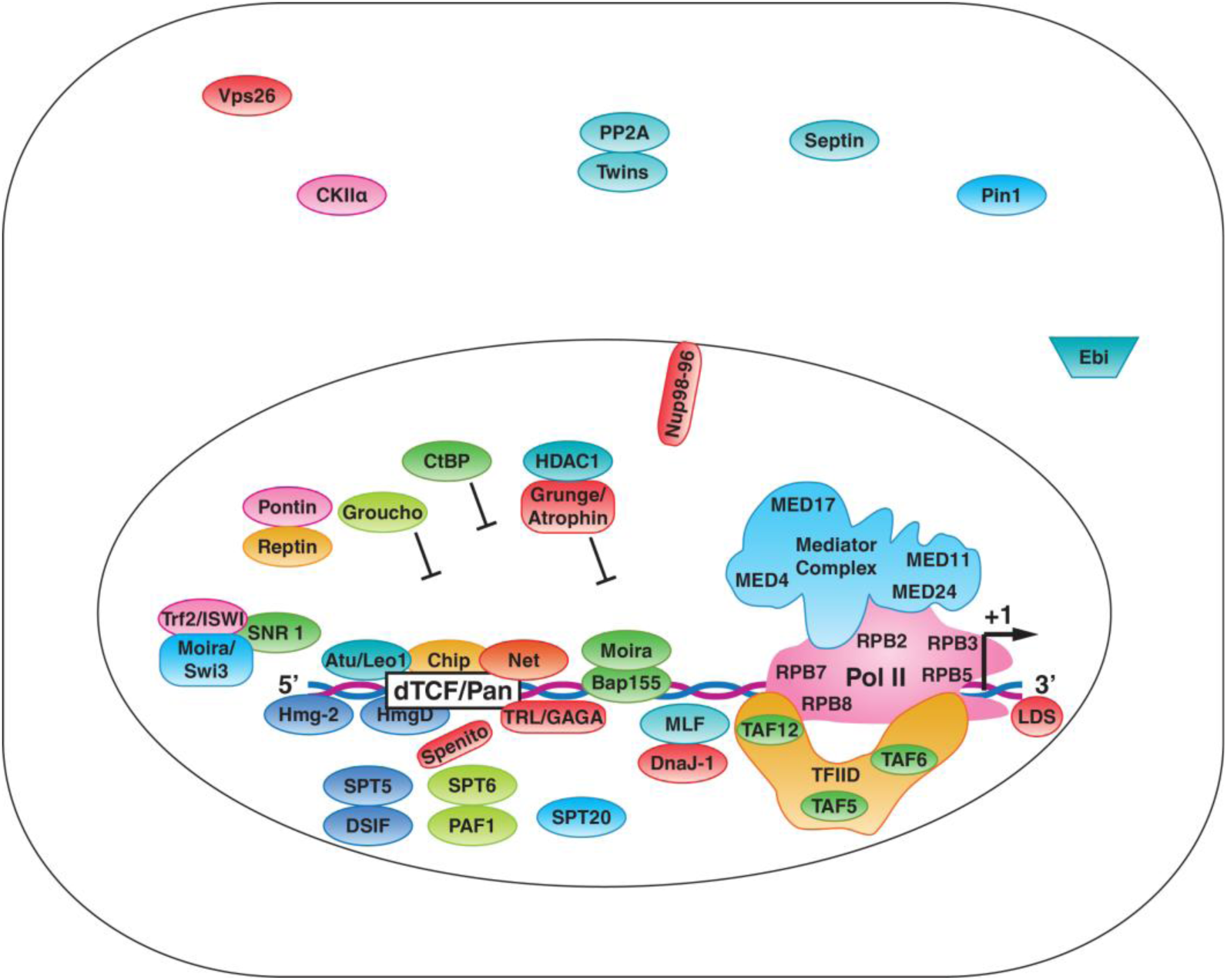
Validation of the SSMEP protocol through the recovery of known Wnt/Wg regulators from embryonic nuclear extracts using W-CRM. Identified proteins include known Wnt/Wg-associated regulators and transcriptional machinery components, such as HMG proteins (HmgD, Hmg-2), transcription factors (Chip, Atu, Net, Trl), and general repressors (CtBP, Groucho, Atrophin). Chip (Chi) – highly enriched specifically at Pos W-CRM and *Myc* Late-Enhancer P1/P2 (absent from SNF input); both regulatory elements co-cluster in heat map. This architectural cofactor of the Wnt enhanceosome; has been reported to contribute to enhancer–promoter communication and chromatin organization. Additional factors include Mediator complex, RNA pol II subunits, and TFIID components (Supplementary Data S9: Table S9h). MED24 was uniquely associated with the heat-map pair linking the positive control and the *Myc* upstream Late-Enhancer cluster (P1/P2) and is implicated in larval salivary gland histolysis, pupal development (Wang et al., 2010), and ecdysone signaling during metamorphosis (Ihry and Bashirullah, 2014).

Among enriched factors, the Pol II-specific transcription co-factor Chip and MED24, both responsive to signaling inputs and enhancer activity, were specifically enriched at both W-CRM and P1/P2 and were absent from nuclear input. W-CRM and Myc-CRM (P1/P2) cluster together in the heat map as functionally related groups; both elements associate with active promoters and signaling-responsive transcriptional regulation (Table S6; Figs. S3 and S4).

In addition, HmgD, Hmg-2, and CtBP were enriched at both elements. HmgD and Hmg-2 bend DNA in association with the SWI/SNF remodeling complex to promote accessibility at active promoters of target genes (Dragan et al., 2003; GO Reference Genome Project, 2011), whereas the nuclear protein CtBP acts as a bifunctional regulator of the Wingless pathway in mediating repression and, in part, activation of Wnt/Wg targets (Bhambhani et al., 2011). Together, the recovery of these Wnt/Wg-associated factors supports the use of W-CRM as a positive control for SSMEP affinity pulldown. Additional validation analyses are provided in Supplementary Results S3.10.

Several additional Wnt/Wingless-associated factors were identified, including Groucho (*gro*), Atrophin-Grunge (*Gug/Atro*), and Net. These factors interact with TCF/Lef-dependent transcriptional regulation and integrate differentiation and signaling cues within Wnt-responsive regulatory complexes during development (Fiedler et al., 2015).

Further modulators of Wnt/Wg signaling included Casein kinase IIalpha (CKIIα), Armadillo (Arm), and Legless (*lgs*); Armadillo and Legless did not meet enrichment thresholds (Table S6).

Finally, ECM and transport proteins enriched include multifunctional signaling regulator Perlecan (*trol*), AP-2μ (AP-2mu, CG7057), Vps26, and Pmm2, linking W-CRM to morphogen distribution and receptor trafficking (Park et al., 2003; Port et al., 2008).

Together, detection of these factors supports recovery of the components of the Wingless pathway, co-clustering with the *Myc* Late-Enhancer (P1/P2), and validates the SSMEP protocol. Detailed regulatory factors enriched at W-CRM are in Results S3.7 and full protein lists are provided in Supplementary Data S3 and S6. The full list of factors associated with the W-CRM are provided in Supplementary Data S10, Table S10g.

### 3.8. Subdivisions of the factors associated with Myc cis-Regulatory modules (Myc-CRMs)

Deciphering the transcriptional control of *MYC* is critical, as it regulates ∼15% of transcribed genes (Dhanasekaran et al., 2022). Using our novel protein purification strategy coupled with LC-MS/MS discovery proteomic, we identified candidate *Myc* regulators among Myc-CRMs–associated factors. Analyses included heat-map (Figs. S3–S4), Pie Charts (Fig. 9; Supplementary Data S9: Tables S9a–S9h), protein databanks, and protein complex annotation (Table S8). Proteins from all categories regulating *Myc* at the transcriptional, posttranscriptional, and protein modifications level—associated with each Myc-CRM were consolidated into various subclassifications of factors associated with *Myc* regulatory networks at multiple levels (Fig. 11; Supplementary Data: Tables S10a– S10g). Candidate interactors identified with only a single unique peptide and low PSM (1-2) support are indicated by an asterisk (*) in the corresponding supplementary tables and should be interpreted with appropriate caution pending further experimental validation.

**Fig. 11.**
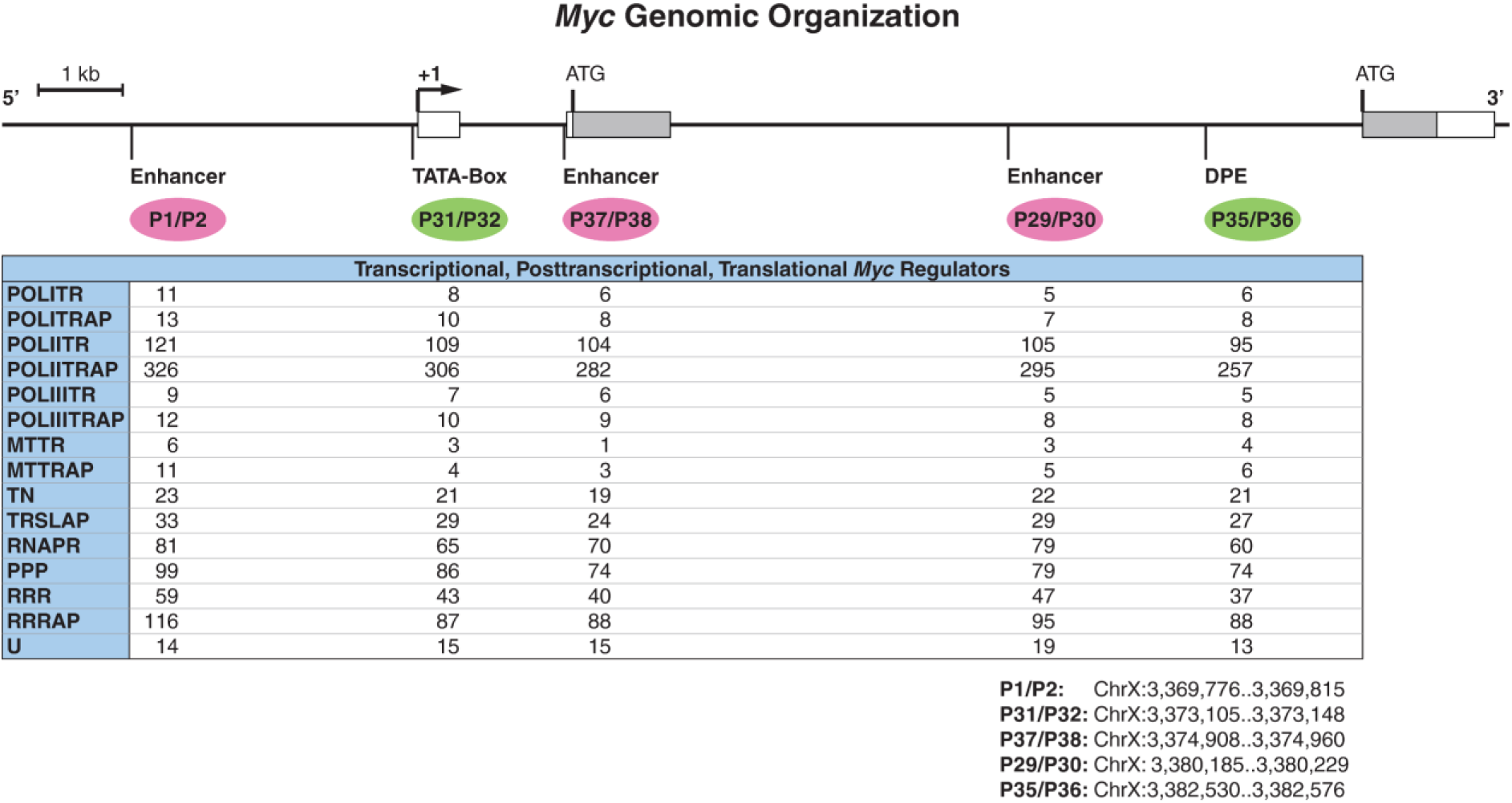
Subdivisions of regulatory factors associated with Myc-CRMs as potential *Myc* regulators based on their mode of regulatory involvement. Enhancers (upstream P1/P2, 5′-proximal P37/P38, and intron 2 P29/P30) are indicated in magenta; promoter elements (TATA-box-containing P31/P32 and DPE P35/P36) are shown in green. The Myc-CRMs are organized according to their positions within the *Myc* genomic locus. **Abbreviations**: POLITR, RNA Pol I Transcription; POLITRAP, RNA Pol I Transcription-Associated Processes; POLIITR, RNA Pol II Transcription; POLIITRAP, RNA Pol II Transcription-Associated Processes; POLIIITR, RNA Pol III Transcription; POLIIITRAP, RNA Pol III Transcription-Associated Processes; MTTR, Mitochondrial Transcription; MTTRAP, Mitochondrial Transcription-Associated Processes; TN, Translation; TRSLAP, Translation-Associated Processes; RNAPR, RNA Processing; PPP, Posttranscriptional/Posttranslational Processes; RRR, Replication-Repair-Recombination; RRRAP, Replication-Repair-Recombination-Associated Processes; U, Uncharacterized. A full list of potential *Myc* regulators, including factors involved in transcription, post-transcriptional/post-translational processes, and protein modification, is provided in Table S3; detailed transcriptional category analyses are shown in Tables S10a–S10g.

We evaluated the association of these proteins with Myc-CRMs, their putative regulatory roles, and their connections to developmental signals linked to *Myc* expression modulation and cell cycle progression.

We characterized early and late enhancer elements and their trans-acting factors controlling *Myc* expression during growth and patterning. We also screened for novel transcription initiation sites beyond the known P1 (TATA) and P2 (DPE) promoters (Kharazmi et al., 2012). The proximal enhancer P37/P38 is active in early embryos and during oogenesis (Fig. 1B). Associated factors include PIC components, TRF2, and co-transcriptional RNA processing factors, suggesting a potential initiation site producing eRNAs during early cell cycle phases (Fig. 13; Table S12). The distal enhancer P1/P2 is required for *Myc* patterning in larval tissues (Fig. 1A,B). Factors enriched at this enhancer include the differentiation-inducing T-box transcription factor DOC3, Chip (involved in wing development and associated with long-range enhancer–promoter interactions), the E-box repressor Net, and MED24 (required for pupal development), indicating that P1/P2 acts as a late differentiation enhancer.

The DPE promoter (P35/P36) requires the adjacent enhancer P29/P30 for developmental patterning of *Myc* (Fig. 2A). The recruitment of TfIIEalpha (TFIIEα), TAF6, and SPT20 by the DPE enhancer (factors which are absent from the DPE promoter) contributes to stable PIC formation and productive elongation (Fig. 13).

Posttranscriptional regulators were also considered (Fig. S6). These factors influence mRNA stability, translation, ribosome biogenesis, and protein turnover, thereby providing feedback regulation that fine-tune *Myc* expression in response to proliferation or differentiation signals.

#### 3.8.1. Trans-acting factors associated with Myc cis-Regulatory modules (Myc-CRMs)

For clarity, subdivisions (Fig. 11; Supplementary Data, Tables S10a–S10g) were consolidated into five main groups: Component of transcription machinery (**Blue**), Chromatin remodelers / histone modifiers / DNA topology **(Purple)**, Direct regulators **(Orange)**, Indirect regulators **(Yellow)**, and Putative / unconfirmed / novel (Gray):

*1. Component of transcription machinery (Blue)*: This category represents the general RNA Pol II transcriptional apparatus required for transcription of the *Myc* gene. Its enrichment supports the validity of the SSMEP method by demonstrating the capture of core factors that assemble at active *cis*-regulatory modules (CRMs) regulating *Myc*. These components participate in transcription initiation at promoter elements such as TATA and DPE, as well as enhancer-dependent transcription. These include Pol II subunits (Polr2B/RpII140/RPB2, Polr2C/RPB3, Polr2E/RPB5, Polr2G/RPB7, Polr2H/RPB8, and RPII215/RPB1); TFIID components (TAF5, TAF6, TAF12, and Cabeza); TFIIH subunits (CycH and Ssl1); TFIIEα (TfIIEalpha), Mediator subunits (MED4, MED11, MED13 (Skuld), MED17, MED 24, MED25), and Nclb/dPWP (TFIIH/Pol I elongation factor implicated in recruitment of Cdk7 to the nucleolus).
*2. Chromatin remodelers / histone modifiers / DNA topology (Purple):* Proteins in this group are well-established companions of chromatin context, that modulate chromatin structure, relieve torsional stress, slide nucleosomes, and contribute to DNA accessibility. Main candidates consist of NURF (Iswi), SWI/SNF (Snr1, Moira/BAP155), PAF1C (Paf1, Leo1), CtBP, HDAC1, HmgD, Hmg-2, BAP55, BAP111, MRG15, SAYP/e(y)3, Ash2, CP190, mod(mdg4, SPT6, SPT20, Etl1, ROW, Bin1/Sin3A, Caf1-55, His2Av, BAF, BEAF-32, RecQ4, and Top2.
*3. Direct regulators (Orange)*: DNA-binding / high-confidence *Myc* regulators group explicitly ties candidates to SSMEP results; this group is intentionally small and focused on factors with strong mechanistic plausibility for direct regulatory activity at *Myc* CRMs. The factors in this group include Chip (*Chi*); TRF2/DREF; TRL/GAF; Net (CG11450); Hairless; Mad; Groucho; Atrophin-Grunge (*Gug/Atro*); DOC3; Lid/Kdm5; and Scribbler (*sbb, mtv, brakeless*).
*4. Indirect regulators (Yellow)*: These factors likely confer *Myc* transcription functional support, with potential direct or indirect DNA association or regulatory involvement. Prominent members are Pontin, Reptin, AP-2μ, Wdr62, Mael, Piwi, Putzig, Su(fu), Cropped/AP-4, MLF, Psi, Hfp, Ebi, and SMRTER.
*5. Putative / unconfirmed / novel (Gray)*: Low-confidence, possible false positives, putative unknown regulators constitute the main fraction of this group (CG6026, CG10527, CG34417, CG9641, CG10543, CG15506), kinases (Cdk1, Cdk5, Wee1, TLK), small GTPases (Rap1), SkpA, Rolled (*rl*), kinase Misshapen (msn), replication/repair (Cutlet (CTF18-RFC)), and posttranscriptional/posttranslational [Modulo, pAbp, Sumo (smt3), CNBP, Cpsf160, (Su(sable), Cpsf5, CPSF6, Cdt2 (l(2)dtl), Rumi, Larp4B, Pumilio/Pop2, How/Struthio, Abstrakt, EJC core components (eIF4A3/CG483, Ddx1, Mago/Y14(Tsunagi)) and Barentsz (*btz*), Spenito/Nito, Prp31, Su(f)], Ogt/sxc, Pin1/dod, Spaghetti/Spag, and Rumi.

These results show that SSMEP captures both direct DNA binders and proteins in complexes such as kinases, phosphatases, splicing factors, ‘minute’ phenotype ribosomal proteins, etc. Direct binders /high-confidence *Myc* regulators, Chip (specific to P1/P2 & W-CRM; Wnt enhanceosome scaffold), TRF2/DREF (TATA-promoter selector), TRL/GAF (chromatin accessibility), and Net (E-box repressor) represent the key findings validating our approach.

For detailed information on factors in all categories, see Supplementary Tables S3, S6, S7, and the simplified Table S12.

#### 3.8.2. RNA Pol II–associated and chromatin remodeling factors at Myc cis-regulatory modules

While core RNA polymerase II (Pol II) transcription machinery is detected across all *Myc cis*-regulatory modules (Myc-CRMs) (see “blue” category), the transition to productive transcription differs depending on promoter architecture and enhancer coupling.

At the mechanistic level, the TATA core promoter (P31/P32) assembles a stable and compositionally complete PIC, including Pol II, TFIID, TFIIH, and Mediator subunits. However, reporter assays show that the TATA element alone is insufficient to drive transcription. Full expression across tissues requires integration of distal regulatory inputs, including the upstream Late-Enhancer, intervening sequences, and the proximal Oocyte Element (P37/P38), indicating a dependence on combinatorial long-range regulatory inputs (Fig. 1A).

In contrast, the DPE-containing promoter requires coupling to its cognate proximal enhancer (P29/P30) to achieve transcriptional activity, consistent with an enhancer-dependent mode of promoter function.

Separately, the Oocyte Element (P37/P38) represents a TATA-less developmental regulatory module active in early embryos and ovaries, where it supports formation of a PIC-like transcriptional assembly state and drives *Myc* expression in early embryos and ovaries (Fig. 1C; Fig. 13).

#### 3.8.3. The Myc TATA promoter supports intrinsically competent transcription

The *Myc* TATA-CorePromoter (P31/P32) assembles a complete and transcriptionally competent PIC, including Pol II, TFIID, TFIIH, and Mediator subunits. The co-occurrence of TAF components (TAF5, TAF6, TAF12) and Mediator (MED4, MED11, MED13, MED17, MED25) supports stable PIC formation.

Elongation-associated factors, including SPT6, SPT20, and TFIIH (CycH, Ssl1), are enriched at the TATA promoter, indicating efficient transition from initiation to productive elongation within this promoter context.

Mediator subunit distribution supports enhancer–promoter communication, with MED24 enriched at the upstream Late-Enhancer (P1/P2) and MED25 at the core promoter, suggesting coordination of regulatory inputs across promoter and enhancer regions. The presence of both TAF–TBP and TRF2-associated complexes indicates flexibility in promoter recognition and context-dependent regulation.

These findings are consistent with reporter assays (Fig. 1A), where the largest truncation containing P1/P2, intervening sequences, and the TATA promoter retains expression in all tissues, indicating that the TATA promoter contributes to sustained transcriptional output within a full regulatory context.

Chromatin remodelers (SAYP, SWI/SNF, MRG15, Ash2) and histone-modifying complexes (HDAC1, COMPASS) are detected in association with the transcriptional machinery, supporting chromatin accessibility and transcriptional progression. In addition, detection of RNA processing and termination factors (CPSF components, PABP) indicates tight coupling between transcription and mRNA maturation.

Collectively, these findings indicate that the *Myc* TATA promoter supports intrinsically competent transcription, in which PIC assembly and productive elongation are efficiently coupled, but whose full transcriptional output is dependent on integration of distal and proximal regulatory inputs.

A detailed description of the factors associated with the TATA promoter is provided in Supplementary Results S3.11.

#### 3.8.4. DPE-dependent transcription requires enhancer-mediated activation

SMEP LC–MS/MS analysis indicates that the *Drosophila Myc* DPE promoter (DPE-CorePromoter, P35/P36) does not support assembly of a fully competent PIC in isolation. Reporter assays show that separation of the DPE promoter from its upstream DPE-Enhancer (P29/P30) abolishes *Myc-lacZ* expression, demonstrating that DPE-dependent transcription is strictly enhancer-dependent (Fig. 2).

Consistent with this, the DPE-CorePromoter assembles core PIC components, including Pol II, TFIID (TAF5, TAF12), and TFIIH, indicating that transcription initiation can occur. However, elongation-associated factors are differentially distributed, with SPT6 enriched at the DPE-Enhancer (P29/P30) and PAF1 (PAF1C/Antimeros) associated with the promoter, rather than stably co-assembling at the core promoter in a manner that supports productive elongation, indicating that elongation competence is not fully established at the isolated DPE-CorePromoter.

The DPE-Enhancer (P29/P30) compensates for this limitation by promoting a transcriptionally competent state. When coupled to the DPE promoter, it restores enrichment of initiation and elongation-associated machinery, including TFIIEα, SPT6, SPT20, TFIIH components, PAF1C, and TRF2/DREF, thereby enabling productive transcription. These findings indicate that the enhancer provides a critical step required for elongation competence, likely through facilitating recruitment or stabilization of factors associated with pause release and transcriptional processivity.

Additional factors associated with the DPE enhancer—including chromatin regulators (SAYP, His2Av), signaling mediators, and transcriptional regulators such as TRF2/DREF—support integration of developmental and signaling inputs. The presence of both activating and repressive components (e.g., Hairless, Groucho, CtBP) further suggests that enhancer-dependent DPE transcription is tightly regulated and context-specific.

Thus, while the DPE promoter ca assemble a PIC, it requires enhancer-mediated activation to achieve productive elongation and transcriptional output, distinguishing it mechanistically from the TATA promoter.

A detailed description of the factors associated with the DPE promoter is provided in Supplementary Results S3.12.

#### 3.8.5. The Oocyte Element functions as a transcribing enhancer driving Myc expression in early development

The Oocyte Element (P37/P38) drives strong *Myc-lacZ* expression in ovaries and early embryos despite lacking a canonical core promoter, indicating that it functions as a transcribing enhancer associated with direct engagement of the transcriptional machinery.

In line with this, Pol II subunits, TFIID components, and Mediator subunits are enriched at P37/P38, consistent with assembly of transcriptionally competent complexes. The presence of Mediator components (MED11, MED13, MED17, MED25) and TAFs suggests that enhancer-bound factors facilitate recruitment and stabilization of Pol II, supporting transcription initiation and likely enhancer RNA (eRNA) production.

Notably, elongation-associated factors, including SPT6 and TFIIH components, are also enriched, indicating that transcription at the Oocyte Element proceeds into productive elongation. This distinguishes it from the isolated DPE promoter and aligns it more closely with enhancer-coupled active transcription units.

A broad set of chromatin remodelers, signaling mediators, and developmental regulators further supports a role in integrating inputs required for early oogenesis and embryogenesis. In addition, broad enrichment of RNA processing, ribosome biogenesis, and translational regulators suggests tight coupling between transcriptional activation and downstream gene expression programs required for rapid developmental transitions.

Together, these data indicate that the Oocyte Element acts as a developmental transcribing enhancer, capable of recruiting Pol II, supporting productive elongation, and driving robust *Myc* expression during early development in a manner consistent with enhancer-driven transcriptional activation.

A detailed description of the factors associated with the *Myc* Oocyte Element is provided in Supplementary Results S3.13.

#### 3.8.6. Psi and Hfp contribute to promoter-specific regulation

Psi and Half pint (Hfp), orthologs of the mammalian proteins FBP and FIR (Guo et al., 2016), respectively, were detected at Myc-CRMs and showed promoter-specific regulatory associations. Psi was enriched across all CRMs, whereas Hfp was recovered at high abundance at the Late Enhancer (P1/P2) and at low abundance at the TATA-CorePromoter (P31/P32), DPE-Enhancer (P29/P30), DPE-CorePromoter (P35/P36), and Oocyte Element (P37/P38) (Table S6). Their distribution, together with that of TFIIH and Mediator subunits, suggests context-dependent modulation of transcription linked to promoter architecture.

The enrichment of Skuld/MED13 at the TATA-CorePromoter is consistent with regulatory inputs that may support both activation and repression. In contrast, DPE-associated regions preferentially involve Mediator tail components (e.g., MED25), consistent with activator-associated transcriptional regulation.

These patterns align with conserved mechanisms in which FBP/Psi promotes transcriptional activation and elongation, whereas FIR/Hfp can restrain transcription via TFIIH modulation. Their co-occurrence at *Myc* regulatory elements suggests a dynamic regulatory balance between activation and repression influenced by promoter context and enhancer engagement.

Together, these findings identify Psi and Hfp as *Myc*-associated regulatory factors within promoter- and enhancer-specific transcriptional complexes (Guo et al., 2016), supporting the ability of SSMEP to capture components of *Myc* regulatory chromatin beyond core DNA-binding transcription factors and associated coregulators.

#### 3.8.7. DNA-binding high-confidence regulators identified at the Myc-CRMs **(Orange group)**

*Chip (Chi):* The transcriptional co-factor Chip emerged as one of the direct regulators identified in our LC-MS/MS dataset. Chip peptides were highly and specifically enriched only in the *Myc* Late-Enhancer (P1/P2) and the positive-control W-CRM pull-downs and were absent from the soluble nuclear fraction (SNF) input (Table S6 and S12). This enrichment pattern, together with hierarchical heat map clustering showing co-clustering of P1/P2 and Pos W-CRM samples (Figs. S3 and S4), indicates a highly specific association with these regulatory regions. Chip (the *Drosophila* ortholog of LDB1) is a LIM-domain-binding cofactor implicated in enhancer-associated regulatory complexes and long-range enhancer–promoter organization (Fiedler et al., 2015). It has been reported to participate in multi-protein enhancer assemblies, including Wnt-associated regulatory systems (van Meyel et al., 1999; van Meyel et al., 2000).

The co-enrichment of Chip at the *Myc* P1/P2 late-Enhancer and W-CRM, together with their shared heat-map clustering, therefore, suggests that Chip associates with enhancer regulatory complexes in a context-dependent manner, consistent with its known role in developmental gene regulation.

*TRF2/DREF:* TRF2/DREF was enriched at the TATA-CorePromoter (P31/P32) and detected across all three enhancer clusters Late-Enhancer (P1/P2), DPE-Enhancer (P29/P30), and Oocyte Element (P37/P38) but absent from nuclear extract input, indicating specific association with multiple *Myc* regulatory modules. This pattern is consistent with its role as a core promoter selectivity factor involved in developmental transcriptional regulation (Hochheimer et al., 2002). TRF2/DREF distribution suggests context-dependent involvement of TRF2-associated complexes across promoter and enhancer-linked regulatory regions and may point to its role in.stem cell maintenance, differentiation, and RNA processing (Neves and Eisenman, 2019).

*TRL/GAF:* was enriched at the Myc-CRMs P1/P2, P31/P32, and P37/P38. As a pioneer transcription factor, TRL/GAF is required for early embryonic development, zygotic gene activation, and chromatin accessibility. It contributes to establishing nucleosome-depleted regions and recruiting remodeling complexes, including SWI/SNF (BAP) (Gaskill et al., 2021). These activities establish a permissive regulatory architecture consistent with promoters that support proximal pausing. Because *Myc* is a well-characterized pause-regulated gene that undergoes rapid transcriptional activation via the release of paused Pol II (Rahl et al., 2010), this accessible landscape provides a mechanistic rationale for its rapid induction without requiring direct occupancy profiling.

*Net* (CG11450): was enriched with higher abundance ratio at the E-box containing Late-Enhancer (P1/P2) (Table S1d, CAGCTG, primer #09) and the TATA-CorePromoter (P31/P32), with lower abundance at the DPE-CorePromoter (P35/P36). Net is a bHLH E-box-binding transcriptional repressor, binds E-boxes (CACGTG), and competes with Myc/Max binding to E-box (Brentrup et al., 2000). Its enrichment at Myc-CRMs points to a potential occupancy at E-box motif within these regions, suggesting an architectural regulatory or competing mechanism for tissue-specific gene gating.

*Hairless (H):* was strongly enriched at P1/P2, P29/P30, and P37/P38, and absent from nuclear input. Hairless is a DNA-binding Notch pathway co-repressor (Su(H)-associated) involved in transcriptional repression and developmental patterning. Its identification suggests association with Notch-linked regulatory complexes at Myc-CRMs, consistent with pathway crosstalk.

*Mothers against dpp (Mad):* a Dpp/BMP effector, was absent from nuclear input but detected at multiple Myc-CRMs with low abundance (Tables S6 and S12). Mad Possesses established role in repressing *Myc* expression during G1/S transition (Firth and Baker, 2005) and in mediating Dpp-dependent transcriptional repression via Mad/Medea/Schnurri complexes (Pyrowolakis et al., 2004). Mad represents a notable regulator associated with Myc-CRMs within our dataset.

*Groucho (Gro)*: was enriched at all Myc-CRMs at low abundance (Tables S6, S7, and S12). Groucho is a conserved transcriptional co-repressor and has been shown to form a physical complex with Myc to regulate a subset of Myc target genes (Orian et al., 2007), integrating EGF and Notch signaling inputs. Its association suggests broad participation in repressive regulatory complexes linked to *Myc cis*-regulatory architecture.

*DOC3:* was associated selectively with the *Myc* Late-Enhancer (P1/P2) and TATA-CorePromoter (P31/P32). DOC3 is a T-box transcription factor downstream of Dpp/TGF-β controlling proliferation arrest and differentiation initiation (Hatton-Ellis et al., 2007; Pflugfelder et al., 2017). DOC3 enrichment at the *Myc* Late-enhancer suggests association with tissue-specific regulatory inputs affecting *Myc* patterning during cell differentiation and larval development.

DOC3 binding to *Myc* Late-Enhancer confirms the specificity of our protocol SSMEP in protein sample affinity purification and administration by LC-MS/MS protein identification.

*Lid/Kdm5:* (Little imaginal discs/Lysine demethylase 5), a nuclear chromatin histone demethylase, was predominantly enriched at the Late-Enhancer (P1/P2) and the DPE-Enhancer (P29/P30), indicating a role as an enhancer-associated factor regulating *Myc* transcription (Secombe et al., 2007). Notably, several proteins identified in our SSMEP–LC–MS/MS pulldown do not directly bind the *Myc cis*-regulatory modules (CRMs), despite being well-established regulators of *Myc* expression. This indicates that their enrichment is likely mediated through indirect or chromatin-based regulatory states rather than strict, sequence-specific binding, highlighting a broader network where non-DNA-binding factors still contribute to *Myc* regulation.

*Atrophin-Grunge (Gug/Atro):* was enriched at the TATA/DPE-Core Promoter (P31/P32, P35/P36) and the Oocyte Element (P37/P38). Grunge is a SANT-MYB domain transcriptional regulator that contains a DNA-binding HTH motif, a protein-protein interaction SANT domain, and ELM2 signature capable of both (Charroux et al., 2006; Zhang et al., 2013). It functions downstream od EGFR and Hedgehog signaling during body patterning. Its presence suggests association with promoter- and enhancer-linked regulatory complexes responding to signaling pathway inputs.

Scribbler (*sbb, mtv, brakeless*): a C2H2 zinc finger corepressor, was highly enriched at multiple Myc-CRMs. Regulated by Ultrabithorax (*Ubx*), Scribbler modulates DPP receptor Thickveins levels to fine-tune morphogen gradient during tissue size determination (Crickmore and Mann, 2006). Scribbler has also been reported to interact with Grunge and to associate with DNA-bound repressive complexes involved in developmental cofactor regulation (Zhang et al., 2002). Because *Myc* is highly responsive to developmental signaling inputs—including DPP, Hedgehog, and EGFR (Fig. 12)—the co-enrichment of Grunge and Scribbler at *Myc*-CRMs suggests their joint participation in indirect, chromatin-based repressive complexes that influence *Myc* expression.

**Fig. 12.**
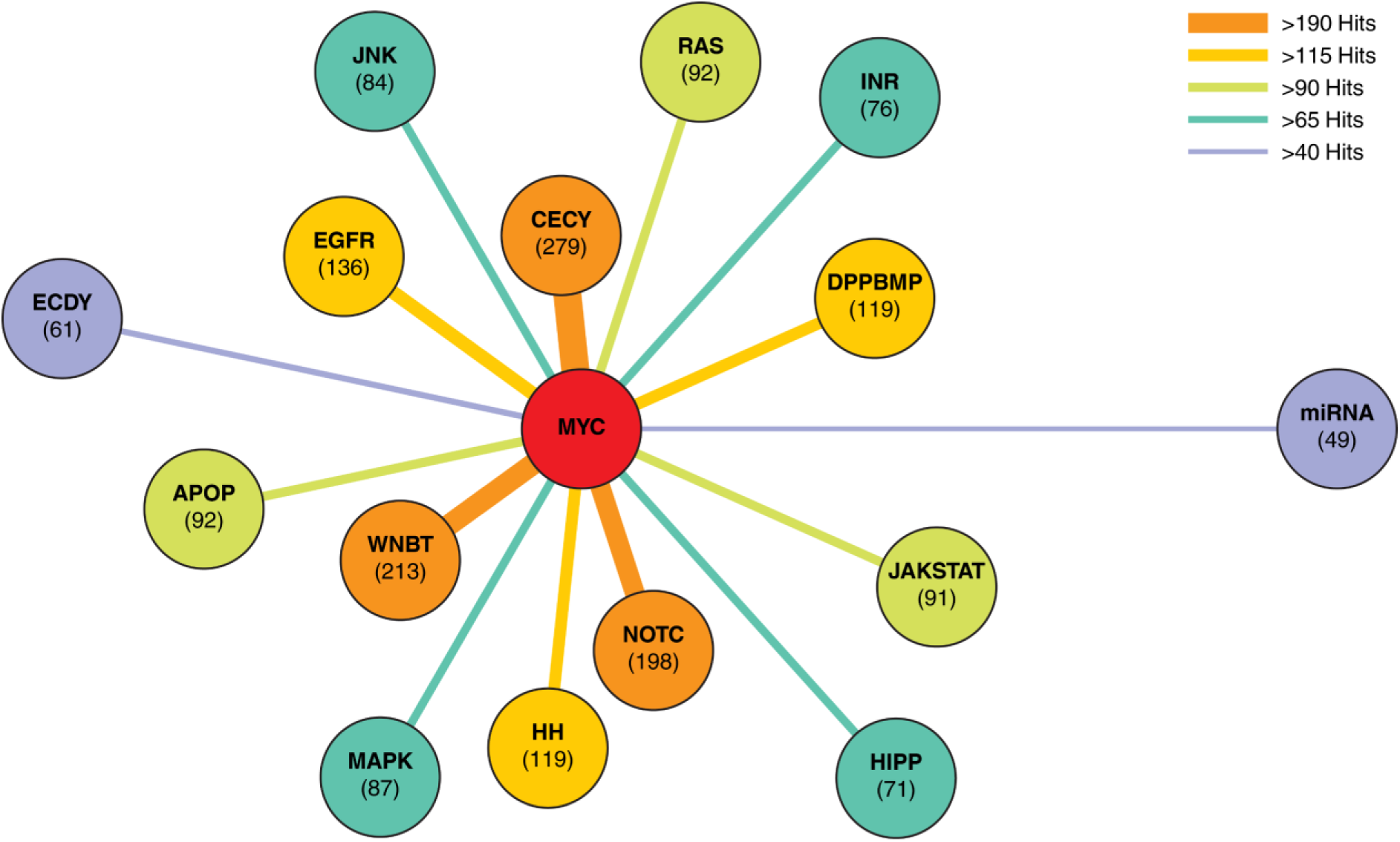
Cell cycle and signaling pathways as potential regulators of *Myc* and *wingless/wg* expression. Identified factors predominantly belong to cell cycle regulators and signaling pathways, including Wingless, Notch, Hedgehog (Hh), EGFR, BMP/DPP, MAPK, JAK/STAT, FGFR, and others. Most of the tested Myc-CRMs and the positive control (W-CRM) show enriched association with these factors (see Table S11a–S11f). Numbers indicate protein counts per cell cycle category or pathway. Abbreviations: CECY, Cell cycle; WNBT, canonical Wnt signaling (Wnt/β-Catenin/TCF); NOTC, Notch; EGFR, Epidermal Growth Factor; DPPBMP, DPP/BMP; HH, Hedgehog; RAS, RAS GTPase; APOP, Apoptosis; JAKSTAT, JAK/STAT; JNK, Janus kinase; INR, Insulin-like receptor; HIPP, Hippo; MAPK, Mitogen-activated protein kinase; ECDY, Ecdysteroid; miRNA, miRNA-mediated signaling.

##### Summary

The enrichment of signaling-associated repressors and activators across Myc-CRMs supports a complex regulatory environment integrating multiple developmental pathways (Dpp, Notch, EGFR, and Hedgehog). These findings are in line with the ability of the SSMEP-LC-MS/MS approach to capture *Myc*-associated regulatory proteins and protein complexes, including both direct DNA-binding factors and chromatin-associated regulators.

#### 3.8.8. Indirect regulators associated with the Myc-CRMs (**Yellow group**)

The factors in this group primarily function as cell cycle regulators, transcriptional cofactors, or RNA-processing regulators that indirectly support *Myc* transcription.

*Pontin (RUVBL1)*: participates in Wnt/β-catenin signaling and MYC-associated transcriptional regulation, and functions as part of the TIP60 chromatin remodeling complex. ATP-dependent helicase; co-activator in transcriptional activation and chromatin remodeling (Bauer et al., 2000).

*Reptin (RUVBL2)*: participates in Wnt/β-catenin signaling and MYC-associated transcriptional regulation, and functions as part of the TIP60 chromatin remodeling complex. Paralog of Pontin; often acts antagonistically or cooperatively in chromatin remodeling and transcription regulation (Bauer et al., 2000).

*Mael (Maelstrom)*: piRNA pathway / transposon silencing. Genome stability factor; represses transposable elements in germline via RNA-mediated mechanisms (Sienski et al., 2012).

*Piwi*: piRNA pathway.

Argonaute family protein; mediates transcriptional and post-transcriptional silencing via piRNAs; chromatin-linked repression (Sienski et al., 2012). Piwi and Mael were detected at the TATA-CorePromoter (P31/P32), with Piwi additionally at the Late-Enhancer (P1/P2) (Fig. 13; Table S6), indicating association of piRNA pathway components with *Myc* regulatory regions.

**Fig. 13.**
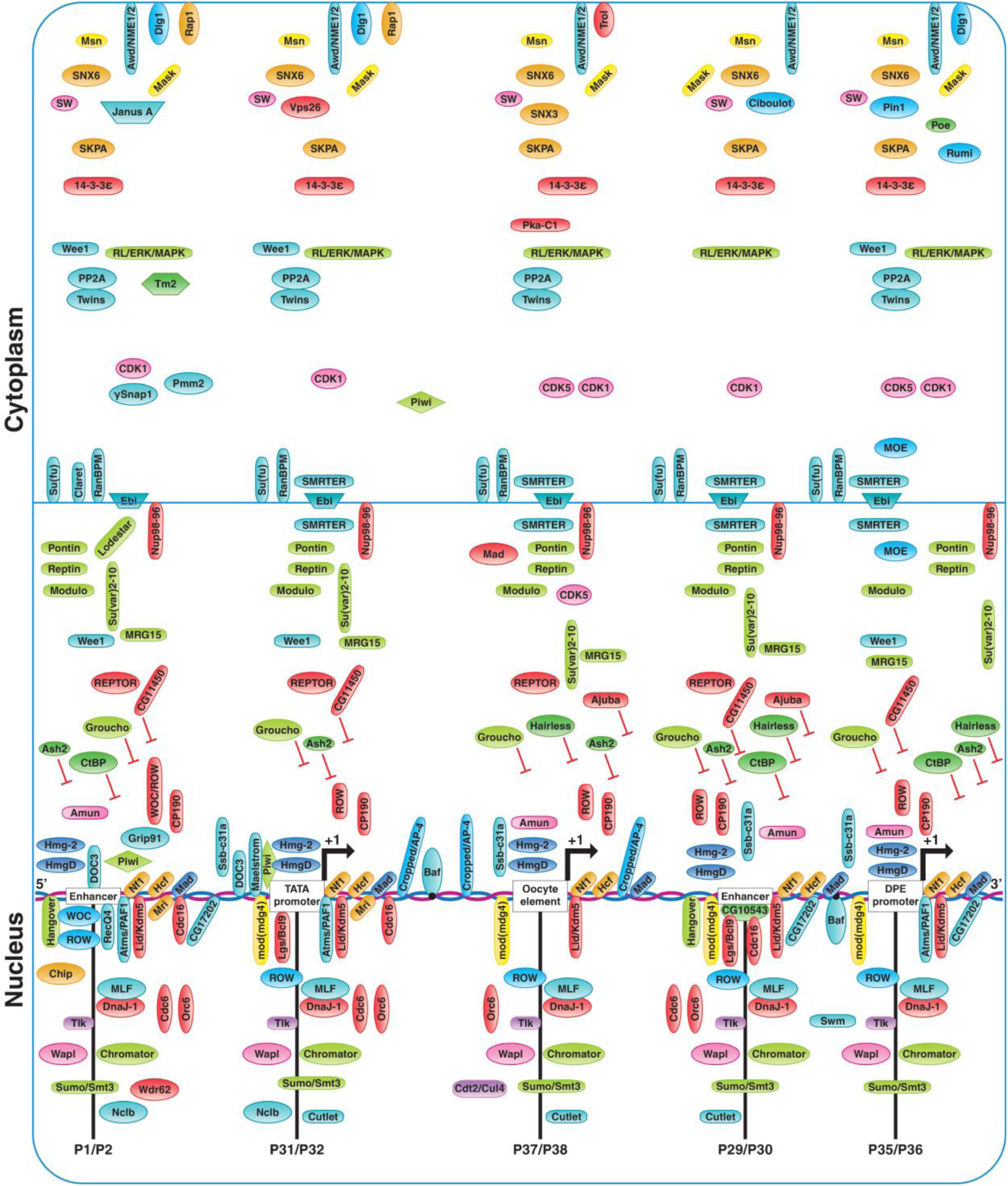
Representatives of regulatory factors across defined categories associated with Myc-CRMs. Doc3, an inhibitor of proliferation and pro-differentiation transcription factor, was identified with the distal Late-Enhancer (P1/P2) cluster and TATA-CorePromoter (P31/P32). Modulo (Myc partner for cell growth), HMG proteins (HmgD and Hmg-2), MLF, HCF, and transcription factor Mad were associated with all tested Myc-CRMs. Barrier to autointegration factor (BAF), was specifically linked to Oocyte Element (P37/P38) and DPE-CorePromoter (P35/P36). Piwi associated with the Late-Enhancer cluster and TATA-CorePromoter (P31/P32), whereas its interactor Maelstrom was restricted to the TATA-CorePromoter. Hairless, a Notch antagonist, associated with the DPE-CorePromoter/-Enhancer (P29/P30) and the Oocyte Element (P37/P38).

*Putzig (Pzg)*: Dpp/BMP, Notch; chromatin (NURF complex). Chromatin regulator required for proliferation control and gene regulation via nucleosome remodeling (Kugler and Nagel, 2007).

*Su(fu) (Suppressor of fused)*: Hedgehog signaling. Negative regulator of Ci/Gli transcription factors; controls transcriptional output of Hh pathway (Methot and Basler, 2000).

*Cropped / AP-4*: Myc network / E-box transcription. bHLH transcription factor; binds E-box motifs and is linked to growth control and *Myc*-related transcription programs (Jung et al., 2008).

*Psi*: RNA processing / transcriptional regulation (Myc-associated). KH-domain RNA-binding protein; regulates transcription and RNA processing, including *Myc* expression (Quinn et al., 2004).

*Hfp (Half pint)*: Splicing factor involved in Pol II transcription and RNA processing. It modulates *Myc* mRNA stability and processing. By interacting with Haywire (*hay*)—the *Drosophila* TFIIH subunit orthologous to human XPB—Hfp participates in promoter melting during transcription initiation (Quinn et al., 2004).

*Ebi*: F-box-like/WD repeat-containing; regulates EGFR, Notch; Wg signaling pathways.

Ubiquitin ligase cofactor; involved in protein turnover and transcriptional repression (Dong et al., 1999). *SMRTER*: Nuclear receptor co-repressor involved in Ecdysone and Notch signaling. It recruits HDACs to mediate hormone-dependent, developmental gene repression (Tsai et al., 2004).

Collectively, the identified regulators map onto major developmental, growth, and cell cycle–associated signaling pathways, including Wingless/Wnt, Notch, Hedgehog, EGFR, BMP/DPP, MAPK, JAK/STAT, Hippo, insulin, and cell cycle control (Fig. 12; Table 1; Table S11a–S11f). The enrichment of pathway-associated factors across both Myc-CRMs and the W-CRM control indicates that *Myc* regulatory regions integrate diverse signaling inputs through direct DNA-binding factors and broader cofactor-associated mechanisms. This supports a model in which *Myc* transcription functions as a central hub integrating developmental and proliferative signaling inputs.

**Table 1.**
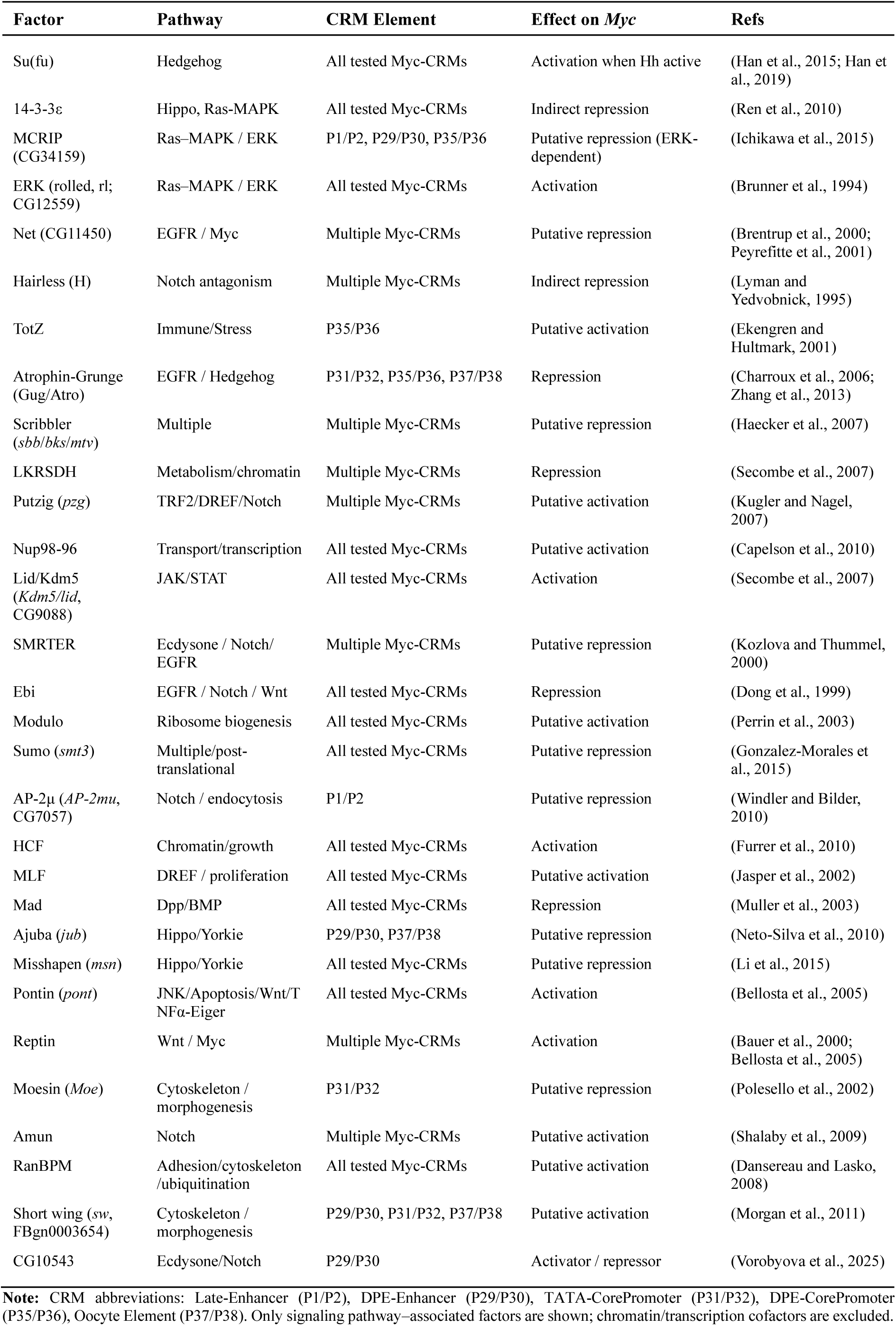
Signaling Pathway Components Recruited by *Myc cis*-Regulatory Modules (Myc-CRMs), LC-MS/MS identified.

#### 3.8.9. Unconfirmed and uncharacterized regulators of Myc expression identified at Myc-CRMs (Gray group)

Factors in this category include various kinases, phosphatases, RNA processing factors, ubiquitin ligases, and uncharacterized proteins. Numerous candidates are components of complexes involved in DNA synthesis, ribosomal biogenesis, cell cycle modulation, and signal transduction. Although they do not bind directly to Myc-CRMs, these factors may indirectly influence *Myc* expression at multiple levels.

I. *Unknown, cell cycle, and replication-associated regulators*

CG6026: putative chromatin-associated factor; transcriptional regulator. Uncharacterized protein enriched at P37/P38; associated with Pol II occupancy, suggesting a potential indirect role in transcriptional regulation at Myc-CRMs.

CG9641: DUF4780 domain-containing protein; unknown molecular function; potential regulatory role remains uncharacterized.

CG10543: Notch-associated transcriptional regulator; C2H2 zinc finger GGGA motif-binding protein (Wolfe et al., 2000); overlaps a Hairless binding site at the DPE enhancer (Table S1d, primer #23) and co-occurs with Groucho and CtBP, suggesting association with Notch-dependent repressive complex (Vorobyova et al., 2025). *Cdk1*: cell cycle kinase. Master mitotic kinase controlling G2/M transition (Lehner and O’Farrell, 1990); may indirectly influence *Myc* expression via cell cycle–dependent regulation.

*Cdk5*: Non-canonical kinase for cell cycle. It drives neuronal differentiation (e.g., *Drosophila* axon patterning). In mammals, Cdk5-mediated phosphorylation of MYC at Ser-62 disrupts its BIN1 interaction, promoting tumorigenesis (Zhang et al., 2019). Enrichment of Bin1/SAP18 suggests Cdk5 modulates Myc activity and protein levels during post-mitotic neuronal development.

*Wee1*: cell cycle checkpoint kinase; CDK inhibitor; negative regulator of Cdk1 via inhibitory phosphorylation (Ghelli Luserna di Rora et al., 2020); may modulate *Myc* expression through cell cycle control.

*TLK*: Tousled-like kinase; chromatin assembly regulator; a conserved S phase kinase regulates chromatin assembly via ASF1 (Carrera et al., 2003); linked to developmental signaling pathways (Yeh et al., 2015; Zhang et al., 2016), suggesting potential indirect regulatory involvement at *Myc* locus.

*Rap1*: Ras GTPase, a component of telomere and signaling complexes implicated in transcriptional modulation (Shore and Bianchi, 2009).

*SkpA*: SCF ubiquitin ligase component core E3 ligase subunit regulating protein turnover (Roberts et al., 2012), may indirectly influence *Myc* regulation through effects on other regulatory proteins in the complex.

*Misshapen (msn)*: MAP4K kinase upstream JNK pathway (Su et al., 1998); may influence *Myc* expression via stress-responsive signaling pathways.

*Cutlet (cutlet)*: CTF18-RFC complex replication clamp loader; DNA replication and cohesion factor (Hanna et al., 2001); indirectly linked to transcriptional regulation through replication-coupled chromatin states.

II. *Canonical RNA biogenesis factors with well-established roles*

*pAbp*: poly(A)-binding protein; mRNA stability

*Cpsf5 / CPSF6 / CPSF160*: 3′ end processing factors

*Su(f)*: transcription termination / 3′-end formation; poly(A) site selection (Jablonowski et al., 2023)

*Prp31*: spliceosome component (pre-mRNA splicing)

*NMD and the EJC Core (eIF4A3, Ddx1, Mago/Y14/Tsunagi)*: splicing-dependent mRNP marking, export, and Nonsense-Mediated mRNA Decay (NMD). It functions as a positional signal, enabling the surveillance machinery to identify premature stop codons and trigger the degradation of aberrant or regulatory transcripts.

*Barentsz (btz)*: mRNA export and splicing / translation coupling (Palacios et al., 2004)

*Abstrakt*: RNA helicase; splicing and oocyte polarity (Kara et al., 2023)

These factors regulate mRNA processing, export, and stability, suggesting indirect modulation of *Myc* transcript fate.

III. *mRNA processing and fate regulators*

*Pumilio / Pop2*: translational repression + mRNA decay control (Arvola et al., 2020)

*CNBP*: IRES-dependent translation regulator of *Myc* (e.g., during wing development) (Antonucci et al., 2014).

*Spenito/Nito*: splicing and Wnt-linked regulation (Abou Faycal et al., 2016)

*How / Struthio*: RNA-binding translational control in development

*Larp4B*: translational repression of *Myc* mRNA and a negative regulator of growth, resulting in reduced organ size specifically through the reduction of cell size (Funakoshi et al., 2018)

These proteins contribute to post-transcriptional regulation of *Myc* expression.

IV. *Post-translational regulatory modifiers*

*Sumo (smt3)*: Small ubiquitin like modifier; SUMOylation pathway modifier; modulates Toll, Dpp, Hedgehog, and Ras/MAPK pathways during embryonic patterning and mitosis (Gonzalez-Morales et al., 2015)

*Cdt2 (l(2)dtl)*: Cul4-DDB1 E3 ligase adapter; S-phase protein degradation regulator (Sloan et al., 2012); exclusively enriched at Oocyte Element (P37/P38), suggesting possible role in Myc protein turnover in developmental contexts

*Su(sable)*: Suppressor of sable; RNA processing / termination factor; regulates non-canonical 3′-end processing and termination in collaboration with Wdr82 (Brewer-Jensen et al., 2016)

*Ogt/sxc*: O-GlcNAc transferase / Super sex combs; O-GlcNAcylates signaling proteins and stabilizes human MYC (Gambetta et al., 2009; Itkonen et al., 2013); enrichment at all Myc-CRMs suggests conserved association with Myc stability regulation.

*Nmt*: N-myristoyltransferase; regulates myristoylation of developmental proteins during tissue patterning; MAX myristoylation prior to degradation supports proper MYC activity during development (Ntwasa et al., 1997; Martin et al., 2011)

*Pin1/dod/*: Peptidylprolyl isomerase Dodo (human Pin1); regulates MYC stability via post-translational modifications (Farrell et al., 2013)

*Rumi*: O-glucosyltransferase; contains EGF-domain; Notch pathway regulator (Acar et al., 2008); exclusive enrichment at the DPE-CorePromoter (P35/P36) suggests potential pathway-level coupling with *Myc* regulation

##### Summary

These findings indicate that cell cycle regulators, RNA processing factors, and post-translational modifiers may indirectly contribute to *Myc* regulatory networks through chromatin state, transcript processing, or protein stability pathways, influencing developmental proliferation and differentiation programs.

Further details are provided in Results S3.14 and Fig. S6.

Table S7 presents the complete set of candidate regulatory proteins identified after sequential filtering of the proteomic dataset, including ribosomal proteins and ribosome-associated complexes. To prioritize candidates with potential direct or indirect roles in chromatin and transcriptional regulation, as well as uncharacterized factors with potential regulatory functions, a curated subset excluding ribosomal proteins is provided in Table S19. The potential contribution of these candidate proteins to *Myc* regulation during developmental growth is considered further in the Discussion.

## 4. Discussion

Using the SSMEP protocol we developed, proteins associated with Myc-CRMs were isolated using *Drosophila* embryonic nuclear extracts, and candidate *Myc* regulators were identified by LC-MS/MS. The SSMEP provides an efficient and sensitive approach to isolate low-abundance, CRM-associated proteins from complex nuclear extracts, enabling systematic identification of proteins associated with defined regulatory DNA elements such as those controlling *Drosophila Myc*. Application of SSMEP to Myc-CRMs identified a high-confidence set of direct regulators (orange group), including TRF2/DREF, TRL/GAF, Net, and Chip. These candidates exhibit CRM-specific enrichment patterns, absence from SNF input where relevant, and co-clustering in quantitative heat maps, together with established mechanistic roles that support their direct involvement in *Myc* transcriptional regulation. These include factors controlling *Myc* transcription, post-transcriptional processing, and mRNA and protein stability.

The analyzed CRMs (summarized in Table 2)—Late-Enhancer (P1/P2), DPE-Enhancer (P29/P30), TATA-CorePromoter (P31/P32), DPE-CorePromoter (P35/P36), and Oocyte Element (P37/P38)—were selected based on conserved motifs and functional validation in reporter assays, as demonstrated in previous studies and further examined here (Figs. 1–2; Table S1d; Kharazmi et al., 2012).

**Table 2.** Investigated *Myc cis*-regulatory modules (Myc-CRMs).

| Original Name | New Name | Description |
| --- | --- | --- |
| P1/P2 | Late-Enhancer | Distal upstream enhancer cluster active during larval and pupal development stages. |
| P31/P32 | TATA-CorePromoter | Core promoter containing the TATA box, essential for transcription initiation. |
| P37/P38 | Oocyte Element | Proximal 5'-UTR conserved sequence block functioning independently of DPE and TATA, active during oogenesis and early embryogenesis. |
| P29/P30 | DPE-Enhancer | Downstream enhancer immediately upstream of and required for the function of the DPE-containing core promoter. |
| P35/P36 | DPE-CorePromoter | Core promoter containing the DPE, essential for transcription initiation, positioned downstream of the TATA element. |

**Note:** CRMs are listed from the 5' to 3' end of the *Myc* gene. “Original Name” refers to experimental IDs; “New Name” reflects functional classification. Descriptions summarize functional activity identified in reporter assays.

Our findings indicate that mitogenic and developmental signals assemble transcription-related proteins at *Myc* promoters and enhancers, enabling PIC formation and integration of trans-acting factors from multiple enhancer clusters. This architecture likely underlies the precise spatiotemporal control of *Myc* during proliferation, growth, and tissue patterning. Together, these results support a model in which *Myc* regulation emerges from coordinated enhancer–promoter integration of signaling inputs.

To benchmark specificity, we applied the approach to a validated Wnt *cis*-regulatory module (W-CRM) containing a TCF/LEF site (Fig. S5; Archbold et al., 2014), identifying known and candidate Wingless (Wg)-associated regulatory proteins alongside components of the Notch, Hedgehog, Dpp, EGFR, and MAPK pathways (Fig. 1o; Tables S6; Supplementary Data, Table S9b). This is consistent with established developmental crosstalk, e.g., cooperative Wg/Notch in wing patterning (Couso et al., 1995) and further supports the broader conclusion that the method captures pathway-associated regulatory complexes at comparable *cis*-regulatory elements.

These CRM-specific protein ensembles indicate that *Myc* transcription is not governed by a single uniform initiation mechanism but instead depends on promoter architecture–specific coordination between pre-initiation complex (PIC) assembly and productive elongation.

### 4.1. Affinity purification–coupled proteomics reveals proteins associated with Myc cis-regulatory elements involved in transcription regulation

Our combined reporter and proteomic analyses demonstrate that *Myc* transcription is not governed by a single uniform initiation mechanism but instead depends on promoter architecture–specific coupling between pre-initiation complex (PIC) assembly and productive elongation. Recovery of the core RNA polymerase II machinery, TFIID, TFIIH, TFIIE, Mediator, and associated RNA-processing factors across *Myc cis*-regulatory modules establishes SSMEP–LC-MS/MS as a versatile affinity purification platform for capturing native regulatory protein complexes. Importantly, the recovery of PIC components together with promoter- and enhancer-specific differences in elongation factor distribution demonstrates that SSMEP captures transcriptionally relevant complexes rather than nonspecific CRM pull-down proteins.

Notably, PIC components are also detected at the Oocyte Element (P37/P38), which retains full *lacZ* activity upon deletion of the adjacent TATA promoter (Fig. 14; Fig. 1C), indicating that this element is capable of directly engaging the basal transcription machinery and supporting transcription in a TATA-independent manner. However, the differential distribution of elongation-associated factors indicates that PIC formation alone does not predict transcriptional output, highlighting a key limitation of purely compositional proteomic datasets.

**Fig. 14.**
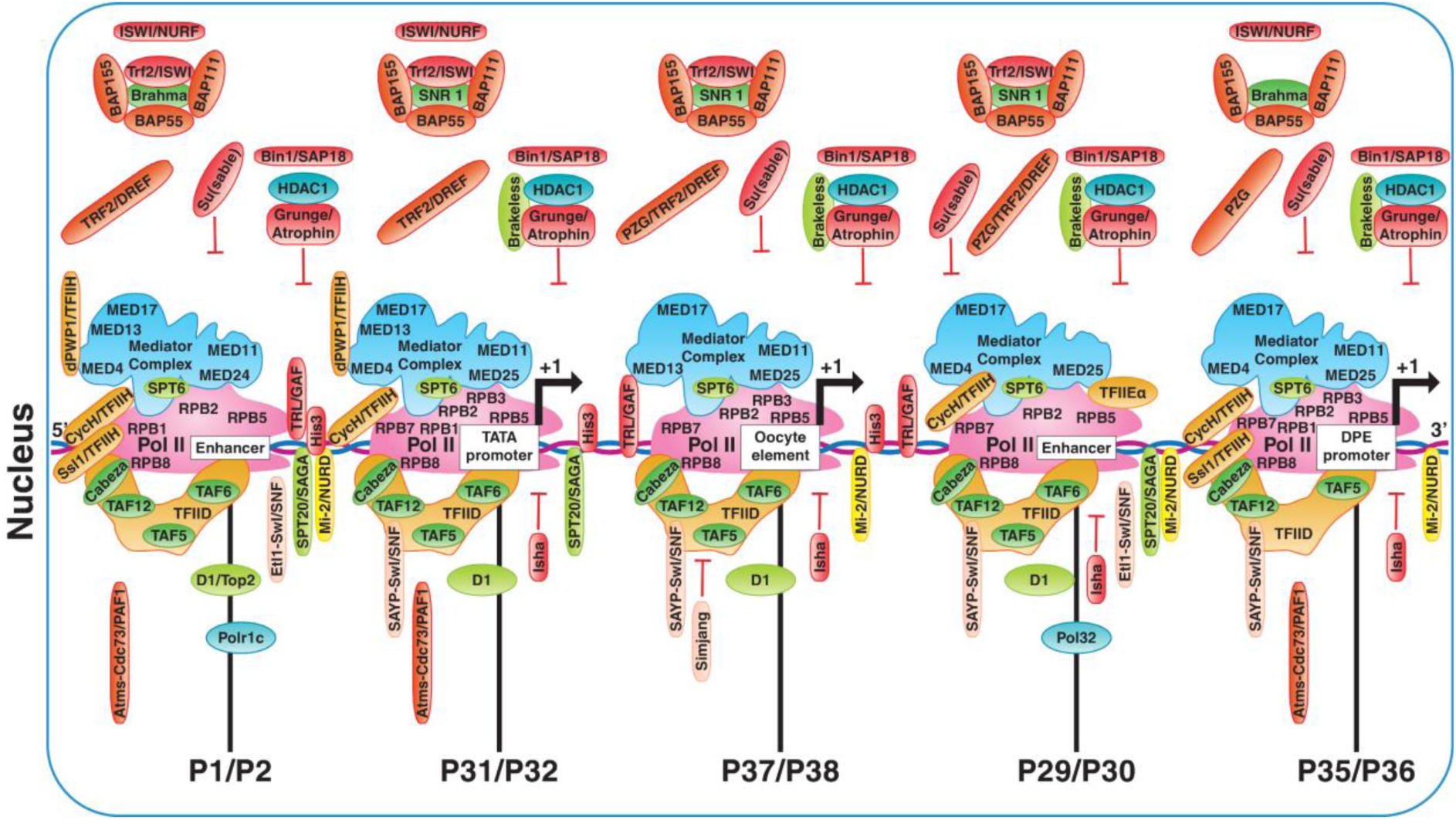
Core promoter machinery, basal transcription factors, and chromatin regulators identified with Myc-CRMs. Mediator complex, TFIID, and RNA Pol II subunits were identified in SSMEP assemblies recovered with the TATA-CorePromoter (P31/P32), Oocyte Element (P37/P38), and DPE-CorePromoter (P35/P36) baits. MED13/Skuld was absent from the DPE-CorePromoter (P35/P36) and the functionally linked enhancer P29/P30, and MED24 was uniquely identified with the Late-Enhancer (P1/P2) bait. Components of chromatin and histone modification complexes, insulator-related factors, and additional transcriptional cofactors identified by SSMEP are also included. See Tables S3, S6, S8, and S12 for factor associations and abundance ratios.

At the TATA promoter, PIC assembly is efficiently coupled to elongation, consistent with enrichment of TFIIH and PAF1C components and with truncation reporter assays showing retained activity (Fig. 1A). In contrast, the DPE-CorePromoter (P35/P36) assembles a PIC but requires coupling to its upstream enhancer to achieve productive transcription. In this context, SPT6 is enriched at the DPE-Enhancer (P29/P30), together with TRF2/DREF, inline with elongation-associated factors being supplied in trans from enhancer regulatory regions rather than being stably encoded at the core promoter (Fig. 14). These findings suggest a mechanistic distinction in which TATA promoters are intrinsically PIC-competent with efficient elongation coupling, whereas DPE promoters require enhancer-mediated licensing to achieve productive elongation.

The Oocyte Element (P37/P38) further extends this framework by functioning as a transcribing developmental enhancer that directly recruits Pol II and supports productive transcription in ovaries and early embryos in the absence of a canonical core promoter. Its strong activity in these contexts suggests that enhancer-driven transcription can operate independently of canonical promoter architecture while maintaining full engagement of the transcriptional machinery.

The recovery of these regulatory complex components by SSMEP supports a broader landscape of transcriptional regulation at *Myc* initiation sites, extending beyond individual DNA-binding trans-acting factors to multiprotein regulatory assemblies. Together, these results define spatiotemporal regulatory switches in which *Myc* transcription is controlled by promoter–enhancer coupling and elongation licensing across developmental contexts.

### 4.2. The distal Late-Enhancer (P1/P2) acts as a Myc differentiation enhancer

The distal Late-Enhancer (P1/P2) is required for full *Myc* expression during larval and pupal development. *Myc-lacZ* reporters retaining only the TATA and proximal Oocyte Element (P37/P38) show abolished activity in larval brains, imaginal discs, and ovaries compared to the full 7.2 kb upstream region that includes P1/P2 and the intervening sequences. P1/P2 also supports expression when paired with the DPE promoter, although full activity depends on appropriate enhancer–promoter distance (Fig. 1F).

SSMEP affinity purification followed by LC-MS/MS identified a set of factors enriched with P1/P2 relative to negative control (Scm). These include T-box transcription factor DOC3, cofactor Chip (Chi), the E-box repressor Net (consistent with the E-box motif in Myc-CRM P1/P2), Groucho (Gro), Antimeros/Paf1, and multiple subunits of SWI/SNF complex, as well as Wnt pathway components (Fig. 13; Table S6). P1/P2 co-clusters with the W-CRM in the heat map, supporting its role in integrating differentiation and patterning inputs.

Among the enriched factors, Topoisomerase II (Top2) and RecQ4 showed notable enrichment with P1/P2. Given their established roles in DNA topology, replication, and genome maintenance, their identification raises the possibility that P1/P2 may recruit factors involved in coordinating enhancer activity with DNA metabolic processes.

These proteomic results highlight the specificity and sensitivity of the SSMEP method for identifying both known transcription regulatory factors and new candidates that align with P1/P2 tissue-specific *Myc* control during larval development. Together data from smaller truncations at the core promoters, these findings indicate distinct functional contributions of each element while demonstrating that all three initiation sites (TATA, DPE, and P37/P38) support PIC assembly, with varying requirements for enhancer-dependent licensing of productive elongation. Further studies will be required to determine the precise mechanisms and functional significance of these interactions.

### 4.3. Coupling of transcription with cotranscriptional RNA processing at Myc cis-regulatory modules

Our SSMEP-LC-MS/MS analysis also identified RNA-binding proteins, splicing factors, and RNA biogenesis components associated with Myc-CRMs, consistent with the well-established coupling of transcription and co-transcriptional RNA processing. This proteomic signature is most prominent at the transcribing Oocyte Element (P37/P38), aligning with the early cell cycle phases when these factors are highly abundant in the developing embryo and oocyte. The association of these components from native extracts with CRMs indicates that the SSMEP platform identifies known, context-specific expression dynamics (Fig. S6; Table S8).

Across modules, a subset of factors (Larp4B, Pumilio, Pop2, CNBP, How/Struthio) modulates mRNA stability, localization, and translation, suggesting a potential buffering layer that stabilizes Myc output against fluctuating transcriptional inputs. Module-specific enrichments are also evident: the TATA (P31/P32) and Late-Enhancer (P1/P2) CRMs preferentially associate with RNA processing, splicing, and mRNA surveillance components (e.g., Prp31, EJC subunits), whereas the Oocyte Element recruits regulators such as Effete/UbcD1 tailored to oogenesis and early embryogenesis. These associations suggest a coordinated transcriptional–post-transcriptional feedback loop that ensures precise Myc dosing during rapid developmental growth.

Collectively, these data support a predictive regulatory network wherein *Myc cis*-acting elements share core PIC formation capacity but differ in enhancer-dependent elongation licensing and post-transcriptional coupling. This regulatory architecture may enable context-specific tuning of Myc levels, as observed in both normal development and oncogenesis.

While high-resolution, locus-specific crosslinking assays will be required to fully map the precise spatial topology of these complexes, these proteomic profiles provide a valuable resource for future functional studies exploring control of *Myc* expression during development.

#### Strengths and Limitations

The *Drosophila* crude nuclear embryonic extracts used in this study contain the full protein pools and are minimally manipulated to preserve the native condition of proteins, thereby reducing artifacts that can arise from more isolated protein samples. This means that our results represent interactions in a biologically relevant context. However, the complexity of these extracts may result in the enrichment of CRM-associated protein complexes in addition to direct DNA-binding transcription factors. Nevertheless, the CRMs identified gene-specific proteins, including known and putative transcriptional regulators, as well as diverse regulatory proteins.

The use of CRMs as baits for affinity purification is a well-established approach for identifying regulatory factors associated with gene expression. Although this strategy may not recover all factors involved in Myc-dependent transcription, it enables the direct enrichment of CRM-associated proteins and regulatory protein complexes from native embryonic extracts. The identified candidates represent a valuable dataset of known and putative transcriptional regulators, as well as diverse post-transcriptional regulatory proteins.

Although LC-MS/MS is a powerful tool for proteomics, it has limited sensitivity for low-abundance or transient proteins. The affinity purification and magnetic enrichment steps employed here—using a long-arm PEG linker (>20 repeats) to minimize steric hindrance—were specifically designed to enhance detection of such proteins while reducing non-specific background. Experimental conditions were optimized to mimic the in vivo environment and maintain protein integrity; however, some losses or alterations inherent to crude extracts cannot be entirely eliminated.

Finally, although the SSMEP protocol was successfully applied to *Drosophila* embryonic extracts, its performance in more complex vertebrate systems, which contain greater protein diversity and additional regulatory layers, remains to be validated. Such studies will be essential to assess the broader applicability of this protocol and to determine whether additional regulatory factors can be recovered in other biological contexts.

Together, these considerations support distinguishing the complete proteomic dataset from the subset prioritized for mechanistic follow-up. Table S7 represents the complete set of candidate regulatory proteins identified after sequential filtering, whereas Table S19 provides a curated subset prioritizing proteins with stronger evidence for chromatin-associated and transcriptional regulatory functions. The curated set provides a focused resource of candidate factors that may contribute directly or indirectly to *Myc* transcriptional regulation and for future functional studies.

#### Concluding remarks

The Myc-CRMs analyzed—Late-Enhancer (P1/P2), DPE-Enhancer (P29/P30), TATA-CorePromoter (P31/P32), DPE-CorePromoter (P35/P36), and Oocyte Element (P37/P38)—may harbor overlapping or adjacent sites allowing simultaneous, sequential, or competitive binding of transcriptional and regulatory factors. The precise sequences within each CRM recognized by transcription factors or other gene-specific factors, their interactions with coregulators and adaptors, and the contributions of Mediator subunits remain to be elucidated.

Using the Solid Surface Magnetic Enrichment Protocol (SSMEP), we generated magnetic Bead–PEG_(23)_–DNA constructs for low-background profiling of nuclear factors across multiple *Myc* regulatory sites. These analyses support the existence of a DPE element in *Drosophila Myc*, first mapped by our group in downstream gene-body sequences, and show that it synergizes with the nearby enhancer (P29/P30) to drive *Myc* expression in response to mitogenic and stress/immunity stimuli. Minimal TATA-less sequences containing the Oocyte Element (P37/P38) are sufficient to drive *Myc* expression in early embryos and maturing oocytes.

The dataset delineates multilayered regulatory control at the *Myc* locus, encompassing chromatin modulators, DNA/RNA modifiers, transcriptional and metabolic factors, and ‘Minute’ proteins, thereby providing a framework for investigating growth, proliferation, and organismal size regulation. SSMEP provides a versatile protocol for identification of regulatory factors to interrogate gene regulatory mechanisms across diverse systems. Beyond the identification of *Myc cis*-regulatory module-binding complexes, the SSMEP–LC-MS/MS platform may be adapted using miRNA mimics to investigate post-transcriptional regulation and miRNA-associated protein complexes or extended to other affinity-based applications, such as antigen–antibody interactions, broadening its utility across diverse biological systems.

Collectively, these findings likely extend the regulatory architecture of the *Myc* locus to promoter-specific control at the TATA-CorePromoter and long-range enhancer-dependent regulation mediated by the Late-Enhancer (P1/P2). The TATA element supports efficient coupling of PIC assembly to elongation, while the Late-Enhancer integrates developmental inputs with chromatin architecture and DNA topology control, ensuring robust transcription during differentiation. Together, these elements coordinate *Myc* expression across developmental contexts. Future locus-specific chromatin analyses will be required to further resolve factor occupancy at individual regulatory elements.

## Supporting information

Supplementary_Table_S13_Volcano_P1P2_Scm

Supplementary_Table_S14_Volcano_Positive_Scm

Supplementary_Table_S15_Volcano_P29P30_Scm

Supplementary_Table_S16_Volcano_P37P38_Scm

Supplementary_Table_S17_Volcano_P31P32_Scm

Supplementary_Table_S18_Volcano_P35P36_Scm

Supplementary_Methods_Results_Refs

README_20260905

## Acknowledgements

The authors thank C. Kerscher (Licor Biosciences) for gel shift image analysis; V. Schwämmle and A. Rogowska-Wrzesinska (University of Southern Denmark) for ComplexBrowser guidance; R. Dubey and M. Calcagni for laboratory facilities; C. Scavenius Sonne-Schmidt (Danish Technological Institute) for LC-MS/MS support; D. Ackermann and C. Winiger for novel oligonucleotide synthesis; H.-P. Lipp for critical manuscript review; and the BroadPharm team for synthesis of the free-end PEG-linker BP-25103 DBCO–PEG_(23)_– amine.

## CRediT authorship contribution statement

**Jasmine Kharazmi:** Conceptualization, Data curation, Formal analysis, Funding acquisition, Investigation, Methodology, Project administration, Resources, Supervision, Validation, Visualization, Writing – original draft, Writing – review & editing.

**Cameron Moshfegh:** Data curation, Formal analysis, Funding acquisition, Methodology, Software, Visualization, Writing – review & editing.

**Thomas Brody:** Formal analysis, Writing – review & editing.

## Funding

J.K. received partial support through a scholarship from the Department of Education of the Canton of Zurich. T.B. is affiliated with the National Institutes of Health (NIH); this research received no specific funding from the NIH. C.M. was supported by ETH Zurich and the ETH Zurich Foundation (University Medicine Zurich Seed Grant).

## Competing interests

The authors declare no competing interests.

## Data Availability Statement

The data that support the findings of this study are available from the corresponding author, Jasmine Kharazmi, upon reasonable request.

## Abbreviations

SSMEP: solid surface magnetic enrichment protocol
EMSA: electrophoretic-mobility shift assay
CRM: cis-regulatory module
DPE: downstream promoter element
SCF: soluble cytoplasmic fraction
SNF: soluble nuclear fraction
Myc-CRM: Drosophila Myc cis-regulatory module
MZT: maternal-to-zygotic transition
W-CRM: Wnt cis-regulatory module
pmol: picomole
DBCO–PEG_(23)_–amine: DiBenzoCycloOctyne-Poly Ethyl Glycol-amine
LC-MS/MS: liquid chromatography mass spectrometry
Scm: scrambled negative control
ORF: open reading frame
CE: capping enzyme
CBC: cap-binding complex
PRO-seq: Precision Run-On sequencing

## Notes

### Competing Interest Statement

The authors have declared no competing interest.

### Summary of Updates

A new supplementary table, Table S19, has been added to provide a curated subset of proteins associated with Myc CRMs from the final proteomic dataset presented in Table S7. The table excludes mitochondrial, ribosomal, and most cytosolic proteins and focuses on candidates with stronger evidence for potential direct or indirect roles in chromatin and transcriptional regulation. Uncharacterized factors with potential transcriptional regulatory functions are also included. The Table S19 legend has been added to clearly describe the selection criteria and scope of the curated dataset. The Results and Discussion sections have been revised to explain the distinction between the complete candidate dataset in Table S7 and the curated subset in Table S19 and to highlight the latter as a focused resource for future functional studies of Myc regulators.

## References

Abou Faycal, C., Gazzeri, S., Eymin, B., 2016. RNA splicing, cell signaling, and response to therapies. Curr Opin Oncol 28, 58–64.

Acar, M., Jafar-Nejad, H., Takeuchi, H., Rajan, A., Ibrani, D., Rana, N.A., Pan, H., Haltiwanger, R.S., Bellen, H.J., 2008. Rumi is a CAP10 domain glycosyltransferase that modifies Notch and is required for Notch signaling. Cell 132, 247–258.

Antonucci, L., D’Amico, D., Di Magno, L., Coni, S., Di Marcotullio, L., Cardinali, B., Gulino, A., Ciapponi, L., Canettieri, G., 2014. CNBP regulates wing development in Drosophila melanogaster by promoting IRES-dependent translation of dMyc. Cell Cycle 13, 434–439.

Archbold, H.C., Broussard, C., Chang, M.V., Cadigan, K.M., 2014. Bipartite recognition of DNA by TCF/Pangolin is remarkably flexible and contributes to transcriptional responsiveness and tissue specificity of wingless signaling. PLoS Genet 10, e1004591.

Arvola, R.M., Chang, C.T., Buytendorp, J.P., Levdansky, Y., Valkov, E., Freddolino, P.L., Goldstrohm, A.C., 2020. Unique repression domains of Pumilio utilize deadenylation and decapping factors to accelerate destruction of target mRNAs. Nucleic Acids Res 48, 1843–1871.

Bauer, A., Chauvet, S., Huber, O., Usseglio, F., Rothbacher, U., Aragnol, D., Kemler, R., Pradel, J., 2000. Pontin52 and reptin52 function as antagonistic regulators of beta-catenin signalling activity. EMBO J 19, 6121–6130.

Bellosta, P., Hulf, T., Balla Diop, S., Usseglio, F., Pradel, J., Aragnol, D., Gallant, P., 2005. Myc interacts genetically with Tip48/Reptin and Tip49/Pontin to control growth and proliferation during Drosophila development. Proc Natl Acad Sci U S A 102, 11799–11804.

Bhambhani, C., Chang, J.L., Akey, D.L., Cadigan, K.M., 2011. The oligomeric state of CtBP determines its role as a transcriptional co-activator and co-repressor of Wingless targets. EMBO J 30, 2031–2043.

Brentrup, D., Lerch, H., Jackle, H., Noll, M., 2000. Regulation of Drosophila wing vein patterning: net encodes a bHLH protein repressing rhomboid and is repressed by rhomboid-dependent Egfr signalling. Development 127, 4729–4741.

Brewer-Jensen, P., Wilson, C.B., Abernethy, J., Mollison, L., Card, S., Searles, L.L., 2016. Suppressor of sable [Su(s)] and Wdr82 down-regulate RNA from heat-shock-inducible repetitive elements by a mechanism that involves transcription termination. RNA 22, 139–154.

Brunner, D., Ducker, K., Oellers, N., Hafen, E., Scholz, H., Klambt, C., 1994. The ETS domain protein pointed-P2 is a target of MAP kinase in the sevenless signal transduction pathway. Nature 370, 386–389.

Capelson, M., Liang, Y., Schulte, R., Mair, W., Wagner, U., Hetzer, M.W., 2010. Chromatin-bound nuclear pore components regulate gene expression in higher eukaryotes. Cell 140, 372–383.

Carrera, P., Moshkin, Y.M., Gronke, S., Sillje, H.H., Nigg, E.A., Jackle, H., Karch, F., 2003. Tousled- like kinase functions with the chromatin assembly pathway regulating nuclear divisions. Genes Dev 17, 2578–2590.

Carroll, P.A., Freie, B.W., Mathsyaraja, H., Eisenman, R.N., 2018. The MYC transcription factor network: balancing metabolism, proliferation and oncogenesis. Front Med 12, 412–425.

Charroux, B., Freeman, M., Kerridge, S., Baonza, A., 2006. Atrophin contributes to the negative regulation of epidermal growth factor receptor signaling in Drosophila. Dev Biol 291, 278–290.

Couso, J.P., Knust, E., Martinez Arias, A., 1995. Serrate and wingless cooperate to induce vestigial gene expression and wing formation in Drosophila. Curr Biol 5, 1437–1448.

Crickmore, M.A., Mann, R.S., 2006. Hox control of organ size by regulation of morphogen production and mobility. Science 313, 63–68.

Dansereau, D.A., Lasko, P., 2008. RanBPM regulates cell shape, arrangement, and capacity of the female germline stem cell niche in Drosophila melanogaster. J Cell Biol 182, 963–977.

Davidson, E.H., Erwin, D.H., 2006. Gene regulatory networks and the evolution of animal body plans. Science 311, 796–800.

Dhanasekaran, R., Deutzmann, A., Mahauad-Fernandez, W.D., Hansen, A.S., Gouw, A.M., Felsher, D.W., 2022. The MYC oncogene - the grand orchestrator of cancer growth and immune evasion. Nat Rev Clin Oncol 19, 23–36.

Dong, X., Tsuda, L., Zavitz, K.H., Lin, M., Li, S., Carthew, R.W., Zipursky, S.L., 1999. ebi regulates epidermal growth factor receptor signaling pathways in Drosophila. Genes Dev 13, 954–965.

Dragan, A.I., Klass, J., Read, C., Churchill, M.E., Crane-Robinson, C., Privalov, P.L., 2003. DNA binding of a non-sequence-specific HMG-D protein is entropy driven with a substantial non-electrostatic contribution. J Mol Biol 331, 795–813.

Eeftens, J.M., van der Torre, J., Burnham, D.R., Dekker, C., 2015. Copper-free click chemistry for attachment of biomolecules in magnetic tweezers. BMC Biophys 8, 9.

Ekengren, S., Hultmark, D., 2001. A family of Turandot-related genes in the humoral stress response of Drosophila. Biochem Biophys Res Commun 284, 998–1003.

Farrell, A.S., Pelz, C., Wang, X., Daniel, C.J., Wang, Z., Su, Y., Janghorban, M., Zhang, X., Morgan, C., Impey, S., Sears, R.C., 2013. Pin1 regulates the dynamics of c-Myc DNA binding to facilitate target gene regulation and oncogenesis. Mol Cell Biol 33, 2930–2949.

Fiedler, M., Graeb, M., Mieszczanek, J., Rutherford, T.J., Johnson, C.M., Bienz, M., 2015. An ancient Pygo-dependent Wnt enhanceosome integrated by Chip/LDB-SSDP. Elife 4.

Fioresi, R., Demurtas, P., Perini, G., 2022. Deep learning for MYC binding site recognition. Front Bioinform 2, 1015993.

Firth, L.C., Baker, N.E., 2005. Extracellular signals responsible for spatially regulated proliferation in the differentiating Drosophila eye. Dev Cell 8, 541–551.

Funakoshi, M., Tsuda, M., Muramatsu, K., Hatsuda, H., Morishita, S., Aigaki, T., 2018. Overexpression of Larp4B downregulates dMyc and reduces cell and organ sizes in Drosophila. Biochem Biophys Res Commun 497, 762–768.

Furrer, M., Balbi, M., Albarca-Aguilera, M., Gallant, M., Herr, W., Gallant, P., 2010. Drosophila Myc interacts with host cell factor (dHCF) to activate transcription and control growth. J Biol Chem 285, 39623–39636.

Gallant, P., Shiio, Y., Cheng, P.F., Parkhurst, S.M., Eisenman, R.N., 1996. Myc and Max homologs in Drosophila. Science 274, 1523–1527.

Gambetta, M.C., Oktaba, K., Muller, J., 2009. Essential role of the glycosyltransferase sxc/Ogt in polycomb repression. Science 325, 93–96.

Gaskill, M.M., Gibson, T.J., Larson, E.D., Harrison, M.M., 2021. GAF is essential for zygotic genome activation and chromatin accessibility in the early Drosophila embryo. Elife 10.

Ghelli Luserna di Rora, A., Cerchione, C., Martinelli, G., Simonetti, G., 2020. A WEE1 family business: regulation of mitosis, cancer progression, and therapeutic target. J Hematol Oncol 13, 126.

GO Reference Genome Project,-. 2011. Phylogenetic annotation using the Gene Ontology.

Goedhart, J., Luijsterburg, M.S., 2020. VolcaNoseR is a web app for creating, exploring, labeling and sharing volcano plots. Sci Rep 10, 20560.

Gonzalez-Morales, N., Geminard, C., Lebreton, G., Cerezo, D., Coutelis, J.B., Noselli, S., 2015. The Atypical Cadherin Dachsous Controls Left-Right Asymmetry in Drosophila. Dev Cell 33, 675–689.

Goodman, J.K., Zampronio, C.G., Jones, A.M.E., Hernandez-Fernaud, J.R., 2018. Updates of the In-Gel Digestion Method for Protein Analysis by Mass Spectrometry. Proteomics 18, e1800236.

Grandori, C., Cowley, S.M., James, L.P., Eisenman, R.N., 2000. The Myc/Max/Mad network and the transcriptional control of cell behavior. Annu Rev Cell Dev Biol 16, 653–699.

Guo, L., Zaysteva, O., Nie, Z., Mitchell, N.C., Amanda Lee, J.E., Ware, T., Parsons, L., Luwor, R., Poortinga, G., Hannan, R.D., Levens, D.L., Quinn, L.M., 2016. Defining the essential function of FBP/KSRP proteins: Drosophila Psi interacts with the mediator complex to modulate MYC transcription and tissue growth. Nucleic Acids Res 44, 7646–7658.

Haecker, A., Qi, D., Lilja, T., Moussian, B., Andrioli, L.P., Luschnig, S., Mannervik, M., 2007. Drosophila brakeless interacts with atrophin and is required for tailless-mediated transcriptional repression in early embryos. PLoS Biol 5, e145.

Han, Y., Shi, Q., Jiang, J., 2015. Multisite interaction with Sufu regulates Ci/Gli activity through distinct mechanisms in Hh signal transduction. Proc Natl Acad Sci U S A 112, 6383–6388.

Han, Y., Wang, B., Cho, Y.S., Zhu, J., Wu, J., Chen, Y., Jiang, J., 2019. Phosphorylation of Ci/Gli by Fused Family Kinases Promotes Hedgehog Signaling. Dev Cell 50, 610–626 e614.

Hanna, J.S., Kroll, E.S., Lundblad, V., Spencer, F.A., 2001. Saccharomyces cerevisiae CTF18 and CTF4 are required for sister chromatid cohesion. Mol Cell Biol 21, 3144–3158.

Hatton-Ellis, E., Ainsworth, C., Sushama, Y., Wan, S., VijayRaghavan, K., Skaer, H., 2007. Genetic regulation of patterned tubular branching in Drosophila. Proc Natl Acad Sci U S A 104, 169–174.

Hochheimer, A., Zhou, S., Zheng, S., Holmes, M.C., Tjian, R., 2002. TRF2 associates with DREF and directs promoter-selective gene expression in Drosophila. Nature 420, 439–445.

Ichikawa, K., Kubota, Y., Nakamura, T., Weng, J.S., Tomida, T., Saito, H., Takekawa, M., 2015. MCRIP1, an ERK substrate, mediates ERK-induced gene silencing during epithelial-mesenchymal transition by regulating the co-repressor CtBP. Mol Cell 58, 35–46.

Ihry, R.J., Bashirullah, A., 2014. Genetic control of specificity to steroid-triggered responses in Drosophila. Genetics 196, 767–780.

Itkonen, H.M., Minner, S., Guldvik, I.J., Sandmann, M.J., Tsourlakis, M.C., Berge, V., Svindland, A., Schlomm, T., Mills, I.G., 2013. O-GlcNAc transferase integrates metabolic pathways to regulate the stability of c-MYC in human prostate cancer cells. Cancer Res 73, 5277–5287.

Jablonowski, C., Quarni, W., Singh, S., Tan, H., Bostanthirige, D.H., Jin, H., Fang, J., Chang, T.C., Finkelstein, D., Cho, J.H., Hu, D., Pagala, V., Sakurada, S.M., Pruett-Miller, S.M., Wang, R., Murphy, A., Freeman, K., Peng, J., Davidoff, A.M., Wu, G., Yang, J., 2023. Metabolic reprogramming of cancer cells by JMJD6-mediated pre-mRNA splicing is associated with therapeutic response to splicing inhibitor. bioRxiv.

Jasper, H., Benes, V., Atzberger, A., Sauer, S., Ansorge, W., Bohmann, D., 2002. A genomic switch at the transition from cell proliferation to terminal differentiation in the Drosophila eye. Dev Cell 3, 511–521.

Jewett, J.C., Bertozzi, C.R., 2010. Cu-free click cycloaddition reactions in chemical biology. Chem Soc Rev 39, 1272–1279.

Johnston, L.A., Prober, D.A., Edgar, B.A., Eisenman, R.N., Gallant, P., 1999. Drosophila myc regulates cellular growth during development. Cell 98, 779–790.

Jung, P., Menssen, A., Mayr, D., Hermeking, H., 2008. AP4 encodes a c-MYC-inducible repressor of p21. Proc Natl Acad Sci U S A 105, 15046–15051.

Kara, E., McCambridge, A., Proffer, M., Dilts, C., Pumnea, B., Eshak, J., Smith, K.A., Fielder, I., Doyle, D.A., Ortega, B.M., Mukatash, Y., Malik, N., Mohammed, A.R., Govani, D., Niepielko, M.G., Gao, M., 2023. Mutational analysis of the functional motifs of the DEAD-box RNA helicase Me31B/DDX6 in Drosophila germline development. FEBS Lett 597, 1848–1867.

Kharazmi, J., Moshfegh, C., 2013. Investigation of dmyc Promoter and Regulatory Regions. Gene Regul Syst Bio 7, 85–102.

Kharazmi, J., Moshfegh, C., Brody, T., 2012. Identification of cis-Regulatory Elements in the dmyc Gene of Drosophila Melanogaster. Gene Regul Syst Bio 6, 15–42.

Kozlova, T., Thummel, C.S., 2000. Steroid regulation of postembryonic development and reproduction in Drosophila. Trends Endocrinol Metab 11, 276–280.

Kugler, S.J., Nagel, A.C., 2007. putzig is required for cell proliferation and regulates notch activity in Drosophila. Mol Biol Cell 18, 3733–3740.

Lehner, C.F., O’Farrell, P.H., 1990. Drosophila cdc2 homologs: a functional homolog is coexpressed with a cognate variant. EMBO J 9, 3573–3581.

Leitner, A., Walzthoeni, T., Kahraman, A., Herzog, F., Rinner, O., Beck, M., Aebersold, R., 2010. Probing native protein structures by chemical cross-linking, mass spectrometry, and bioinformatics. Mol Cell Proteomics 9, 1634–1649.

Li, S., Cho, Y.S., Yue, T., Ip, Y.T., Jiang, J., 2015. Overlapping functions of the MAP4K family kinases Hppy and Msn in Hippo signaling. Cell Discov 1, 15038.

Lyman, D.F., Yedvobnick, B., 1995. Drosophila Notch receptor activity suppresses Hairless function during adult external sensory organ development. Genetics 141, 1491–1505.

Martin, D.D., Beauchamp, E., Berthiaume, L.G., 2011. Post-translational myristoylation: Fat matters in cellular life and death. Biochimie 93, 18–31.

Methot, N., Basler, K., 2000. Suppressor of fused opposes hedgehog signal transduction by impeding nuclear accumulation of the activator form of Cubitus interruptus. Development 127, 4001–4010.

Michalak, W., Tsiamis, V., Schwammle, V., Rogowska-Wrzesinska, A., 2019. ComplexBrowser: A Tool for Identification and Quantification of Protein Complexes in Large-scale Proteomics Datasets. Mol Cell Proteomics 18, 2324–2334.

Morgan, J.L., Song, Y., Barbar, E., 2011. Structural dynamics and multiregion interactions in dynein-dynactin recognition. J Biol Chem 286, 39349–39359.

Muller, B., Hartmann, B., Pyrowolakis, G., Affolter, M., Basler, K., 2003. Conversion of an extracellular Dpp/BMP morphogen gradient into an inverse transcriptional gradient. Cell 113, 221–233.

Neto-Silva, R.M., de Beco, S., Johnston, L.A., 2010. Evidence for a growth-stabilizing regulatory feedback mechanism between Myc and Yorkie, the Drosophila homolog of Yap. Dev Cell 19, 507–520.

Neves, A., Eisenman, R.N., 2019. Distinct gene-selective roles for a network of core promoter factors in Drosophila neural stem cell identity. Biol Open 8.

Ntwasa, M., Egerton, M., Gay, N.J., 1997. Sequence and expression of Drosophila myristoyl-CoA: protein N-myristoyl transferase: evidence for proteolytic processing and membrane localisation. J Cell Sci 110 (Pt 2), 149–156.

Orian, A., Delrow, J.J., Rosales Nieves, A.E., Abed, M., Metzger, D., Paroush, Z., Eisenman, R.N., Parkhurst, S.M., 2007. A Myc-Groucho complex integrates EGF and Notch signaling to regulate neural development. Proc Natl Acad Sci U S A 104, 15771–15776.

Palacios, I.M., Gatfield, D., St Johnston, D., Izaurralde, E., 2004. An eIF4AIII-containing complex required for mRNA localization and nonsense-mediated mRNA decay. Nature 427, 753–757.

Park, Y., Rangel, C., Reynolds, M.M., Caldwell, M.C., Johns, M., Nayak, M., Welsh, C.J., McDermott, S., Datta, S., 2003. Drosophila perlecan modulates FGF and hedgehog signals to activate neural stem cell division. Dev Biol 253, 247–257.

Perrin, L., Benassayag, C., Morello, D., Pradel, J., Montagne, J., 2003. Modulo is a target of Myc selectively required for growth of proliferative cells in Drosophila. Mech Dev 120, 645–655.

Peyrefitte, S., Kahn, D., Haenlin, M., 2001. New members of the Drosophila Myc transcription factor subfamily revealed by a genome-wide examination for basic helix-loop-helix genes. Mech Dev 104, 99–104.

Pflugfelder, G.O., Eichinger, F., Shen, J., 2017. T-Box Genes in Drosophila Limb Development. Curr Top Dev Biol 122, 313–354.

Polesello, C., Delon, I., Valenti, P., Ferrer, P., Payre, F., 2002. Dmoesin controls actin-based cell shape and polarity during Drosophila melanogaster oogenesis. Nat Cell Biol 4, 782–789.

Port, F., Kuster, M., Herr, P., Furger, E., Banziger, C., Hausmann, G., Basler, K., 2008. Wingless secretion promotes and requires retromer-dependent cycling of Wntless. Nat Cell Biol 10, 178–185.

Pyrowolakis, G., Hartmann, B., Muller, B., Basler, K., Affolter, M., 2004. A simple molecular complex mediates widespread BMP-induced repression during Drosophila development. Dev Cell 7, 229–240.

Quinn, L.M., Dickins, R.A., Coombe, M., Hime, G.R., Bowtell, D.D., Richardson, H., 2004. Drosophila Hfp negatively regulates dmyc and stg to inhibit cell proliferation. Development 131, 1411–1423.

Rahl, P.B., Lin, C.Y., Seila, A.C., Flynn, R.A., McCuine, S., Burge, C.B., Sharp, P.A., Young, R.A., 2010. c-Myc regulates transcriptional pause release. Cell 141, 432–445.

Ren, F., Wang, B., Yue, T., Yun, E.Y., Ip, Y.T., Jiang, J., 2010. Hippo signaling regulates Drosophila intestine stem cell proliferation through multiple pathways. Proc Natl Acad Sci U S A 107, 21064–21069.

Roberts, D.M., Pronobis, M.I., Alexandre, K.M., Rogers, G.C., Poulton, J.S., Schneider, D.E., Jung, K.C., McKay, D.J., Peifer, M., 2012. Defining components of the ss-catenin destruction complex and exploring its regulation and mechanisms of action during development. PLoS One 7, e31284.

Samuels, T.J., Jarvelin, A.I., Ish-Horowicz, D., Davis, I., 2020. Imp/IGF2BP levels modulate individual neural stem cell growth and division through myc mRNA stability. Elife 9.

Secombe, J., Li, L., Carlos, L., Eisenman, R.N., 2007. The Trithorax group protein Lid is a trimethyl histone H3K4 demethylase required for dMyc-induced cell growth. Genes Dev 21, 537–551.

Shalaby, N.A., Parks, A.L., Morreale, E.J., Osswalt, M.C., Pfau, K.M., Pierce, E.L., Muskavitch, M.A., 2009. A screen for modifiers of notch signaling uncovers Amun, a protein with a critical role in sensory organ development. Genetics 182, 1061–1076.

Shore, D., Bianchi, A., 2009. Telomere length regulation: coupling DNA end processing to feedback regulation of telomerase. EMBO J 28, 2309–2322.

Sienski, G., Donertas, D., Brennecke, J., 2012. Transcriptional silencing of transposons by Piwi and maelstrom and its impact on chromatin state and gene expression. Cell 151, 964–980.

Sloan, R.S., Swanson, C.I., Gavilano, L., Smith, K.N., Malek, P.Y., Snow-Smith, M., Duronio, R.J., Key, S.C., 2012. Characterization of null and hypomorphic alleles of the Drosophila l(2)dtl/cdt2 gene: Larval lethality and male fertility. Fly (Austin) 6, 173–183.

Smith-Bolton, R.K., Worley, M.I., Kanda, H., Hariharan, I.K., 2009. Regenerative growth in Drosophila imaginal discs is regulated by Wingless and Myc. Dev Cell 16, 797–809.

Su, Y.C., Treisman, J.E., Skolnik, E.Y., 1998. The Drosophila Ste20-related kinase misshapen is required for embryonic dorsal closure and acts through a JNK MAPK module on an evolutionarily conserved signaling pathway. Genes Dev 12, 2371–2380.

Tsai, C.C., Kao, H.Y., Mitzutani, A., Banayo, E., Rajan, H., McKeown, M., Evans, R.M., 2004. Ataxin 1, a SCA1 neurodegenerative disorder protein, is functionally linked to the silencing mediator of retinoid and thyroid hormone receptors. Proc Natl Acad Sci U S A 101, 4047–4052.

van Meyel, D.J., O’Keefe, D.D., Jurata, L.W., Thor, S., Gill, G.N., Thomas, J.B., 1999. Chip and apterous physically interact to form a functional complex during Drosophila development. Mol Cell 4, 259–265.

van Meyel, D.J., O’Keefe, D.D., Thor, S., Jurata, L.W., Gill, G.N., Thomas, J.B., 2000. Chip is an essential cofactor for apterous in the regulation of axon guidance in Drosophila. Development 127, 1823–1831.

Vorobyova, N.E., Nikolenko, I.V., Krasnov, A.N., 2025. CG10543 Protein Is Involved in the Regulation of Transcription of Ecdysone-Dependent Genes. Molecular Biology 59, 646–654.

Wang, L., Lam, G., Thummel, C.S., 2010. Med24 and Mdh2 are required for Drosophila larval salivary gland cell death. Dev Dyn 239, 954–964.

Windler, S.L., Bilder, D., 2010. Endocytic internalization routes required for delta/notch signaling. Curr Biol 20, 538–543.

Wolfe, S.A., Nekludova, L., Pabo, C.O., 2000. DNA recognition by Cys2His2 zinc finger proteins. Annu Rev Biophys Biomol Struct 29, 183–212.

Yeh, T.H., Huang, S.Y., Lan, W.Y., Liaw, G.J., Yu, J.Y., 2015. Modulation of cell morphogenesis by tousled-like kinase in the Drosophila follicle cell. Dev Dyn 244, 852–865.

Zhang, S., Xu, L., Lee, J., Xu, T., 2002. Drosophila atrophin homolog functions as a transcriptional corepressor in multiple developmental processes. Cell 108, 45–56.

Zhang, X., Wang, J., Jia, Y., Liu, T., Wang, M., Lv, W., Zhang, R., Shi, J., Liu, L., 2019. CDK5 neutralizes the tumor suppressing effect of BIN1 via mediating phosphorylation of c-MYC at Ser-62 site in NSCLC. Cancer Cell Int 19, 226.

Zhang, Y., Cai, R., Zhou, R., Li, Y., Liu, L., 2016. Tousled-like kinase mediated a new type of cell death pathway in Drosophila. Cell Death Differ 23, 146–157.

Zhang, Z., Feng, J., Pan, C., Lv, X., Wu, W., Zhou, Z., Liu, F., Zhang, L., Zhao, Y., 2013. Atrophin-Rpd3 complex represses Hedgehog signaling by acting as a corepressor of CiR. J Cell Biol 203, 575–583.

