## Supplementary_Table_S1 for "Functional dissection of *Drosophila Myc cis*-regulatory modules (Myc-CRMs) reveals developmentally active DNA-protein interactions"

| **No.** | **Construct Names** | **Promoter Type** | **Promoter Size (kb) and X Chr. Coordinates** | **Plasmid Size (kb)** | **Myc-CRMs Contained** | **lacZ Activity: Brain, Discs, Emb, Ova** |
| --- | --- | --- | --- | --- | --- | --- |
| 01 | pJ8 | *MYC* In2 | 8.161; ChrX:3,376,232..3,384,387 | 19.819 | P29/P30; P35/P36 | all tested tissues |
| 02 | pJ8.4 | pJ8 enhancer cluster, upstream of DPE | 0.840; ChrX:3,379,408..3,380,247 | 12.497 | P29/P30 | None |
| 03 | pJ8.5 | 3’ end of In2 containing DPE Element | 2.102; ChrX:3,382,334..3,384,387 | 13,760 | P35/P36 | None |
| 04 | pJ2.1 | *MYC* noncoding 5’ end | 7.158; ChrX:3,367,866..3,375,005 | 18.816 | P1/P2; P31/P32; P37/P38 | all tested tissues |
| 05 | pJ7 | -100bp + 5’UTR | 1.928; ChrX:3,373,088..3,375,005 | 13.5 | P31/P32; P37/P38 | Emb, Ova |
| 06 | pJ7.1 | 5’UTR, (-100bp contained in pJ7 deleted) | 1.806;  ChrX:3,373,200..3,375,005 | 13.4 | P37/P38 | Emb, Ova |
| 07 | pJ7.2 | 5’-UTR (In1+non-coding E2) | 1.298; ChrX:3,373,710..3,375,005 | 13 | P37/P38 | Emb, Ova |
| 08 | pJ7.3 | 5’UTR (TATA region + Exon1) | 0.647; ChrX:3,373,088..3,373,728 | 12.3 | P31/P32 | None |
| 09 | pJ7.4 | pJ2.1 Enhancer region fused to pJ7 promoter | 3.498; composite promoter ChrX:3,369,648..3,371,174  ChrX:3,373,044..3,375,005 | 15.158 | P1/P2; P31/P32; P37/P38 | discs weaker than in pJ2.1 |
| 10 | pJ7.5 | J8 Enhancer region fused to J7 promoter | 2.896; composite promoter ChrX: 3,379,324..3,380,254  ChrX:3,373,044..3,375,005 | 14.518 | P29/P30; P31/P32; P37/P38 | all tested tissues |

**Table S1a. Reporter Constructs Used in This Study.**

pJ2.1, pJ7, and pJ8 constructs, along with derived truncations, contains at least one investigated Myc-CRM P1/P2, P29/P30, P31/P32, P35/P36, and P37/P38. pJ7.4 and pJ7.5 include composite promoters.

**Abbreviations:** **bp**, base pair; **DPE**, Downstream Promoter Element; E2, Exon 2; **In1**, Intron 1; **UTR**, Untranslated Region; **Emb**, Embryo; **Ova**, Ovary.

| **No.** | **Primer names** | **Sequence (5′→3′)** | **Size (bp)** | **Usage** |
| --- | --- | --- | --- | --- |
| 01 | J7-F | AGCTCCTATTGGTACCACTTAAAGCGAATT C | 31-mer | PCR - pJ7 |
| 02 | J7-R | CTGGCGAAAGGGGGATGTGCT | 21-mer |  |
| 03 | J1.4-F | AACGCGGCCTTTTTACGGTTC | 21-mer | PCR - pJ7.1-template |
| 04 | J1.4-R | CACCCACCTTTCGGTACCACTTTCACTTTC AC | 31-mer |  |
| 05 | J7.2-F | TAAATAGAGACGGATACCGCGGCTATG TTCAG | 32-mer | PCR - pJ7.2 |
| 06 | J7.2-R | GCTGGCGACCTGCGTTTCAC | 20-mer |  |
| 07 | J7.3-F | CCCGCAGTGAGAGCGCAGAGAG | 22-mer | PCR - pJ7.3 |
| 08 | J7.3-R | CTCGCTGAACATAGTTTCCGTATCCGTC | 28-mer |  |
| 09 | J7.4-F | TCTAGTGTGCCGTTATAAAACCGCGGCGCTGC | 32-mer | PCR - pJ7.4 |
| 10 | J7.4-R | GCGGGTACCTGGAAAGTGTGCATGGCGAGAT | 31-mer |  |
| 11 | In2E-F-Not | GCACTCTTAACAAGCGGCCGCATGTTCTTTTT | 32-mer | PCR - pJ7.5 |
| 12 | In2E-R-Asp | AACAGGTACCACCGCTGTGTTATAAACAATTGCG | 34-mer |  |
| 13 | SV40-F | GATTAGGGCCGCAAGAAAACTATCC | 25-mer | PCR- SV40 poly (A) |
| 14 | SV40-R | TAGGCACCCCAGGCTTTACACTTTAT | 26-mer |  |
| 15 | SV40 | GTTCAGGGGGAGGTGTGGG | 19 | Sequences SV40 P(A) |
| 16 | pCAF | GACGGCGATATTTCTGTGGAC | 21-mer | Forward and Reverse sequencing of pCaSpeR4 |
| 17 | pCAR | CCTTAGCATGTCCGTGGGGTTTGA | 24-mer |  |
| 18 | Acc651_F | GCCGAGAAAATCAAATGGAC | 20-mer | Forward and Reverse sequencing of pJ7.5 |
| 19 | PZR | CGGGCCTCTTCGCTATTACG | 20-mer |  |

**Table S1b. Analytical Primers Used for Plasmid Cloning and Sequencing.**

Oligonucleotides were designed using LaserGene Primer Select (Bioinformatics tool) and synthesized by Microsynth AG, Switzerland. Primers were used to amplify and sequence *Myc* non-coding regions, cloning vectors and injection plasmids. Primer orientation is 5´→3´; length is in base pair (bp). Forward primers are indicated by “F” and the reverse primers by “R.”

| **No.** | **Stock # (Source)** | **Name** | **Genotype / Description** |
| --- | --- | --- | --- |
| 01 | 6598 (Bloomington) | y w | [y^1^](http://flybase.org/reports/FBal0018607.html) [w^1118^](http://flybase.org/reports/FBal0018186.html) |
| 02 | 8412 (Bloomington) | dpp-lacZ | [y^1^](http://flybase.org/reports/FBal0018607.html) [w^1118^](http://flybase.org/reports/FBal0018186.html); [P{dpp-lacZ.Exel.2}3](http://flybase.org/reports/FBti0040759.html) |
| 03 | 2475 (Bloomington) | double balancer | w^*^; [T(2;3)ap^Xa^](http://flybase.org/reports/FBab0007867), [ap^Xa^](http://flybase.org/reports/FBal0000663)/[In(2L)Cy](http://flybase.org/reports/FBab0004410), [In(2R)Cy](http://flybase.org/reports/FBab0004960), [Duox^Cy^](http://flybase.org/reports/FBal0002196); [TM3](http://flybase.org/reports/FBba0000047), [Sb^1^](http://flybase.org/reports/FBal0015145) |
| 04 | 11108 (Kyoto 108991) | blue balancer | Cyo, P{lArB}A66.2F2/[b^1^](http://flybase.org/reports/FBal0018607.html) [Adh^*^](http://flybase.org/reports/FBal0018186.html) [cn^*^](http://flybase.org/reports/FBal0018607.html) l([2)^**^](http://flybase.org/reports/FBal0018186.html); r[y^506^](http://flybase.org/reports/FBal0018607.html) |
| 05 | 64349 (originally Stk#1) (Bloomington) | Canton-S | Canton-S |
| 06 | — (This study) | J7.1 | P{J7.1-lacZ} in y¹ w¹¹¹⁸ (random P-element insertion) |
| 07 | — (This study) | J7.2 | P{J7.2-lacZ} in y¹ w¹¹¹⁸ (random P-element insertion) |
| 08 | — (This study) | J7.3 | P{J7.3-lacZ} in y¹ w¹¹¹⁸ (random P-element insertion) |
| 09 | — (This study) | J7.4 | P{J7.4-lacZ} in y¹ w¹¹¹⁸ (random P-element insertion) |
| 10 | — (This study) | J7.5 | P{J7.5-lacZ} in y¹ w¹¹¹⁸ (random P-element insertion) |
| 11 | — (This study) | J8.4 (new insertion) | Independent P-element insertion in y¹ w¹¹¹⁸ (this study; reproduced construct from Kharazmi et al., 2012) |
| 12 | — (This study) | J8.5 (new insertion) | Independent P-element insertion in y¹ w¹¹¹⁸ (this study; reproduced construct from Kharazmi et al., 2012) |

**Table S1c. Fly Stocks Used in This Study.**

Embryos from *y[1] w[1118]* were used for random P-element transgenesis. *Canton-S* (Bloomington # 64349) embryos 0-72 h AEL) were used for cytoplasmic and nuclear protein extraction. The fly “blue balancer” line expressing *lacZ* in embryos and ovaries, was used as positive control for embryo/ovary staining. The *dpp-lacZ* line, expressing *lacZ* in the pattern of the *dpp* gene, was used as positive control for third instar larval brain and imaginal disc staining. *y[1] w[1118]* flies served as negative control. Transgenic reporter lines (J7.1–J7.5) were generated in this study by random P-element insertion in the y[1] w[1118] background. Multiple independent insertion lines were established and analyzed for each construct (see Table S1c and text for details).

| **No.** | **Target Name** | **Sequence (5′→3′)** | **Size (bp)** | **Identity** | **Modifications** |
| --- | --- | --- | --- | --- | --- |
| 01 | IR5-Scm1 | TGTGTGTGTGTGCGCACGCGCGTGTGTGTGCGTGCATGCGTGCAT | 45 | Non-Dmel scrambled sequence (origin: Mus musculus *Mdm2* gene) | 5’ Dyo 781 |
| 02 | Az5-Scm2-IR3 | ATGCACGCATGCACGCACACACACGCGCGTGCGCACACACACACA | 45 |  | 3’ Dyo781, 5’ Az |
| 03 | Scm2-IR3 | ATGCACGCATGCACGCACACACACGCGCGTGCGCACACACACACA | 45 | The same as above | 3’ Dyo781 |
| 04 | IR5-Scm1-Az3 | TGTGTGTGTGTGCGCACGCGCGTGTGTGTGCGTGCATGCGTGCAT | 45 |  | 5’ Dyo781, 3’ Az |
| 05 | IR5-Pos1 | TACGAT**AAGATCAAAGG**TGCAGT**GCCGCCA**GTCGTA | 36 | HMG & Helper sites configurations in W-CRM | 5’ Dyo781 |
| 06 | Az5-Pos2-IR3 | TACGAC**TGGCGGC**ACTGCA**CCTTTGATCTT**ATCGTA | 36 |  | 3’ Dyo781, 5’ Az |
| 07 | Pos2-IR3 | TACGAC**TGGCGGC**ACTGCA**CCTTTGATCTT**ATCGTA | 36 | The same as above | 3’ Dyo781 |
| 08 | IR5-Pos1-Az3 | TACGAT**AAGATCAAAGG**TGCAGT**GCCGCCA**GTCGTA | 36 |  | 5’ Dyo781, 3’ Az |
| 09 | IR5-P1 | GAAGCTTT**TTGTTG**T**TTTTGCAGCTGC**C**GTTT**TCGCCCTT | 40 | J2.1 enhancer cluster | 5’ Dyo 781 |
| 10 | Az5-P2-IR3 | AAGGGCGA**AAAC**G**GCAGCTGCAAAA**A**CAACAA**AAAGCTTC | 40 |  | 3’ Dyo781, 5’ Az |
| 11 | P2-IR3 | AAGGGCGA**AAAC**G**GCAGCTGCAAAA**A**CAACAA**AAAGCTTC | 40 | The same as above | 3’ Dyo781 |
| 12 | IR5-P1-Az3 | GAAGCTTT**TTGTTG**T**TTTTGCAGCTGC**C**GTTT**TCGCCCTT | 40 |  | 5’ Dyo781, 3’ Az |
| 13 | P3 | GAAGCTTT**TTGTTG**T**TTTTGCAGCTGC**C**GTTT**TCGCCCTT | 40 | The same as above | 5’-biot |
| 14 | P4 | AAGGGCGA**AAAC**G**GCAGCTGCAAAA**A**CAACAA**AAAGCTTC | 40 |  | 3’-biot |
| 15 | IR5-P29 | ATG**CGCGTGGGAAAA**TCTT**ACA**GT**GCAGCGGAAGCA**A**TGTTT**TTC | 45 | Dead Ringer in J8 enhancer region | 5’ Dyo 781 |
| 16 | IR3-P30-Az5 | GAA**AAACA**T**TGCTTCCGCTGC**AC**TGT**AAGA**TTTTCCCACGCG**CAT | 45 |  | 3’ Dyo781, 5’ Az |
| 17 | P30-IR3 | GAA**AAACA**T**TGCTTCCGCTGC**AC**TGT**AAGA**TTTTCCCACGCG**CAT | 45 | Enhancer cluster in J8 enhancer region | 3’ Dyo 781 |
| 18 | IR5-P29-Az3 | ATG**CGCGTGGGAAAA**TCTT**ACA**GT**GCAGCGGAAGCA**A**TGTTT**TTC | 45 |  | 5’ Dyo 781, 3’ Az |
| 19 | IR5-P31 | GA**GCGCGGC**AGTCTGGTACGATAG**AAATTTTATTTAA**GCCACAG | 44 | GC1-TATA1-box region in J2.1, J7, and J7.3 - J7.5 Promoters | 5’ Dyo 781 |
| 20 | Az5-P32-IR3 | CTGTGGC**TTAAATAAAATTT**CTATCGTACCAGACT**GCCGCGC**TC | 44 |  | 3’ Dyo781, 5’ Az |
| 21 | P32-IR3 | CTGTGGC**TTAAATAAAATTT**CTATCGTACCAGACT**GCCGCGC**TC | 44 | The same as above | 3’ Dyo781 |
| 22 | IR5-P31-Az3 | GA**GCGCGGC**AGTCTGGTACGATAG**AAATTTTATTTAA**GCCACAG | 44 |  | 5’ Dyo781, 3’ Az |
| 23 | IR5-P35 | A**TCATTC**ATTCATTGTCTATCGAAAGCGCGGTGGTGGGCTT**GGTCG**C | 47 | DPE-containing region in J8 | 5’ Dyo 781 |
| 24 | Az5-P36-IR3 | G**CGACC**AAGCCCACCACCGCGCTTTCGATAGACAATGAAT**GAATGA**T | 47 |  | 3’ Dyo781, 5’ Az |
| 25 | P36-IR3 | G**CGACC**AAGCCCACCACCGCGCTTTCGATAGACAATGAAT**GAATGA**T | 47 | The same as above | 3’ Dyo 781 |
| 26 | IR5-P35-Az3 | A**TCATTC**ATTCATTGTCTATCGAAAGCGCGGTGGTGGGCTT**GGTCG**C | 47 |  | 5’ Dyo 781, 3’ Az |

**Table S1d continued**

| **No.** | **Target Name** | **Sequence (5′→3′)** | **Size (bp)** | **Identity** | **Modifications** |
| --- | --- | --- | --- | --- | --- |
| 27 | IR5-P37 | A**CAA**CG**ATTT**CCGCCT**TA**TC**TATA**T**TTT**TCA**GACAGGC**ATA**TAACTCAGGAA**C | 53 | 3’ end of In1 & start of Ex2 noncoding region | 5’ Dyo781 |
| 28 | Az5-P38-IR3 | G**TTCCTGAGTTA**TATG**CCTGTC**TGA**AAA**A**TATA**GA**TAA**GGCGG**AAAT**CG**TTG**T | 53 |  | 3’ Dyo781, 5’ Az |
| 29 | P38-IR3 | G**TTCCTGAGTTA**TATG**CCTGTC**TGA**AAA**A**TATA**GA**TAA**GGCGG**AAAT**CG**TTG**T | 53 | The same as above | 3’ Dyo781 |
| 30 | IR5-P37-Az3 | A**CAA**CG**ATTT**CCGCCT**TA**TC**TATA**T**TTT**TCA**GACAGGC**ATA**TAACTCAGGAA**C | 53 |  | 5’ Dyo781, 3’ Az |

**Table S1d. Primer Pairs for EMSA and Identification of *Myc* Regulators.**

Primer pairs (P1/P2 to P37/P38) were derived from *Myc* non-coding Conserved *cis*-Regulatory Regions (Myc-CRMs). Bold nucleotides indicate sequences conserved in all or all but one of five other *Drosophila* species, (*D. sechellia, D. Yakuba, D. erecta, D. willistoni, D. virilis*); gray highlights mark conserved clusters. The Wingless *cis*-Regulatory Module (W-CRM, Archbold et al. 2014) was used as a positive control. A 45 bp scrambled sequence *Mus musculus Mdm2* was used as a negative control ;BLAST against FlyBase and NCBI revealed no matches. Oligonucleotides were synthesized at Microsynth AG (Balgach, Switzerland). Orientation is 5´→3´; lengths are in base pair (bp).

**Abbreviations:** **biot**, biotinylated; **DPE**, downstream promoter element; **Dyo 781**, Dyomics 781 (IR dye); **In**, Intron; **Ex**, Exon; **IR**, Infrared; **Pos**, positive; **Scm**, Scrambled; **Az**, Azide moiety end-modification.

| **No.** | **Annealed Oligo (Full Name)** | **Annealed Oligo (Short Name)** | **Coordinates on X Chromosome** |
| --- | --- | --- | --- |
| 01 | ds IR5-Scm1/Az5-Scm2-IR3 | IR-Scm1/Scm2 | NA |
| 02 | ds Scm2-IR3/IR5-Scm1-Az3 | IR-Scm2/Scm1 |  |
| 03 | ds IR5-Pos1/Az5-Pos2-IR3 | IR-Pos1/Pos2 | NA |
| 04 | ds Pos2-IR3/IR5-Pos1-Az3 | IR-Pos2/Pos1 |  |
| 05 | ds IR5-P1/Az5-P2-IR3 | IR-P1/P2 | ChrX:3,369,776..3,369,815 |
| 06 | ds P2-IR3/IR5-P1-Az3 | IR-P2/P1 |  |
| 07 | ds IR5-P29/Az5-P30-IR3 | IR-P29/P30 | ChrX:3,380,185..3,380,229 |
| 08 | ds P30-IR3/IR5-P29-Az3 | IR-P30/P29 |  |
| 09 | ds IR5-P31/Az5-P32-IR3 | IR-P31/P32 | ChrX:3,373,105..3,373,148 |
| 10 | ds P32-IR3/IR5-P31-Az3 | IR-P32/P31 |  |
| 11 | ds IR5-P35/Az5-P36-IR3 | IR-P35/P36 | ChrX:3,382,530..3,382,576 |
| 12 | ds P36-IR3/IR5-P35-Az3 | IR-P36/P35 |  |
| 13 | ds IR5-P37/Az5-P38-IR3 | IR-P37/P38 | ChrX:3,374,908..3,374,960 |
| 14 | ds P38-IR3/ IR5-P37-Az3 | IR-P38/P37 |  |

**Table S1e. Annealing Schemes for Single-Stranded Oligos to Generate Double-Stranded Myc Target and Control Sequences.**

Each oligonucleotide, (Myc-CRMs, positive, and negative controls) was annealed with its complementary strand to produce double-stranded targets. These were attached to Bead-PEG heterodimers via copper-free “click” chemistry reaction to generate Bead-PEG-DNA constructs for protein binding assays. Genomic coordinates of the Myc-CRMs are indicated on the X chromosome of *D. melanogaster*.

**Abbreviations:** **ds**, double-stranded; **Scm**, Scrambled; **IR**, Infrared; **NA**, not applicable; **P**, primer; **Pos**, positive; **Az**, azide; **biot**, biotinylated; **DPE**, downstream promoter element; **Dyo 781**, Dyomics 781 (IR dye); **In**, Intron; **Ex**, Exon.

| **No.** | **Lab Sample Name** | **Sample Description** | **Protein Subcellular Localization** | **MS Sample Name** |
| --- | --- | --- | --- | --- |
| 1 | Beads Only | Dynabeads Magnetic MyOne Carboxylic | None | 20210527_CSSS_ZUH_1.raw |
| 2 | SCF Only | Soluble Cytoplasmic Fraction | SCF | 20210527_CSSS_ZUH_2.raw |
| 3 | SNF Only | Soluble Nuclear Fraction | SNF | 20210527_CSSS_ZUH_3.raw |
| 4 | IR-Scm1/Scm2:SCF | ds IR5-Scm1/IR3-Az5-Scm2, Scm2 strand attached to Beads | SCF | 20210527_CSSS_ZUH_4.raw |
| 5 | IR-Scm2/Scm1:SCF | ds IR3-Scm2/IR5-Az3-Scm1, Scm1 strand attached to Beads | SCF | 20210527_CSSS_ZUH_5.raw |
| 6 | IR-Scm1/Scm2:SNF | ds IR5-Scm1/IR3-Az5-Scm2, Scm2 strand attached to Beads | SNF | 20210527_CSSS_ZUH_6.raw |
| 7 | IR-Scm2/Scm1:SNF | ds IR3-Scm2/IR5-Az3-Scm1, Scm1 strand attached to Beads | SNF | 20210527_CSSS_ZUH_7.raw |
| 8 | IR-Pos1/Pos2:SNF | ds IR5-Pos1/IR3-Az5-Pos2, Pos2 strand attached to Beads | SNF | 20210527_CSSS_ZUH_8.raw |
| 9 | IR-Pos2/Pos1:SNF | ds IR3-Pos2/IR5-Az3-Pos1, Pos1 strand attached to Beads | SNF | 20210527_CSSS_ZUH_9.raw |
| 10 | IR-P1/P2:SCF | ds IR5-P1/IR3-Az5-P2, P2 strand attached to Beads | SCF | 20210527_CSSS_ZUH_10.raw |
| 11 | IR-P2/P1:SCF | ds IR3-P2/IR5-Az3-P1, P1 strand attached to Beads | SCF | 20210527_CSSS_ZUH_11.raw |
| 12 | IR-P1/P2:SNF | ds IR5-P1/IR3-Az5-P2, P2 strand attached to Beads | SNF | 20210527_CSSS_ZUH_12.raw |
| 13 | IR-P2/P1:SNF | ds IR3-P2/IR5-Az3-P1, P1 strand attached to Beads | SNF | 20210527_CSSS_ZUH_13.raw |
| 14 | IR-P29/P30:SCF | ds IR5-P29/IR3-Az5-P30, P30 strand attached to Beads | SCF | 20210527_CSSS_ZUH_14.raw |
| 15 | IR-P30/P29:SCF | ds IR3-P30/IR5-Az3-P29, P29 strand attached to Beads | SCF | 20210527_CSSS_ZUH_15.raw |
| 16 | IR-P29/P30:SNF | ds IR5-P29/IR3-Az5-P30, P30 strand attached to Beads | SNF | 20210527_CSSS_ZUH_16.raw |
| 17 | IR-P30/P29:SNF | ds IR3-P30/IR5-Az3-P29, P29 strand attached to Beads | SNF | 20210527_CSSS_ZUH_17.raw |

**Table S1f continued**

| **No.** | **Lab Sample Name** | **Sample Description** | **Protein Subcellular Localization** | **MS Sample Name** |
| --- | --- | --- | --- | --- |
| 18 | IR-P31/P32:SCF | ds IR5-P31/IR3-Az5-P32, P32 strand attached to Bead | SCF | 20210527_CSSS_ZUH_18.raw |
| 19 | IR-P32/P31:SCF | ds IR3-P32/IR5-Az3-P31, P31 strand attached to Bead | SCF | 20210527_CSSS_ZUH_19.raw |
| 20 | IR-P31/P32:SNF | ds IR5-P31/IR3-Az5-P32, P32 strand attached to Bead | SNF | 20210527_CSSS_ZUH_20.raw |
| 21 | IR-P32/P31:SNF | ds IR3-P32/IR5-Az3-P31, P31 strand attached to Bead | SNF | 20210527_CSSS_ZUH_21.raw |
| 22 | IR-P35/P36:SCF | IR5-P35/IR3-Az5-P36, P36 strand attached to Bead | SCF | 20210527_CSSS_ZUH_22.raw |
| 23 | IR-P36/P35:SCF | ds IR3-P36/IR5-Az3-P35, P35 strand attached to Beads | SCF | 20210527_CSSS_ZUH_23.raw |
| 24 | IR-P35/P36:SNF | IR5-P35/IR3-Az5-P36, P36 strand attached to Bead | SNF | 20210527_CSSS_ZUH_24.raw |
| 25 | IR-P36/P35:SNF | ds IR3-P36/IR5-Az3-P35, P35 strand attached to Beads | SNF | 20210527_CSSS_ZUH_25.raw |
| 26 | IR-P37/P38:SCF | IR5-P37/IR3-Az5-P38, P38 strand attached to Beads | SCF | 20210527_CSSS_ZUH_26.raw |
| 27 | IR-P38/P37:SCF | ds IR3-P38/IR5-Az3-P37, P37 strand attached to Beads | SCF | 20210527_CSSS_ZUH_27.raw |
| 28 | IR-P37/P38:SNF | IR5-P37/IR3-Az5-P38, P38 strand attached to Beads | SNF | 20210527_CSSS_ZUH_28.raw |
| 29 | IR-P38/P37:SNF | ds IR3-P38/IR5-Az3-P37, P37 strand attached to Beads | SNF | 20210527_CSSS_ZUH_29.raw |

**Table S1f. DNA–Protein Complex Samples for Mass Spectrometry Analysis.**

Two replicates were prepared per sample. Each double-stranded oligo was labeled with Dyomics 781: 5´ on the top strands and 3´ on the complementary bottom strand. In each replicate, only one strand carried an azide modification for copper-free “click” chemistry with PEG23-amine attached to Dynabeads® MyOne™ Carboxylic Acid (Invitrogen, cat. #65001). This design minimized steric hindrance and allowed protein complexes to bind the Bead-PEG-linked DNA efficiently.

**Abbreviations:** **IR**, Infrared; **Dyo 781**, Dyomics 781 (IR dye); **P**, primer; **SCF**, Soluble Cytoplasmic Fraction; **SNF**, Soluble Nuclear Fraction; **Scm**, scrambled; **Pos**, positive; **Az**, azide end-modification.

| **No.** | **Replicate Pair** | **Replicate dsDNA** | **Coordinates on X Chromosome** |
| --- | --- | --- | --- |
| 01 | P1/P2 | 5′-GAAGCTTTTTGTTGTTTTTGCAGCTGCCGTTTTCGCCCTT-3′  3′-CTTCGAAAAACAACAAAAACGTCGACGGCAAAAGCGGGAA-5′ | ChrX:3,369,776..3,369,815 |
| 02 | P29/P30 | 5′-ATGCGCGTGGGAAAATCTTACAGTGCAGCGGAAGCAATGTTTTTC-3′  3′-TACGCGCACCCCTTTTAGAATGTCACGTCGCCTTCGTTACAAAAAG-5′ | ChrX:3,380,185..3,380,229 |
| 03 | P31/P32 | 5′-GAGCGCGGCAGTCTGGTACGATAGAAATTTTATTTAAGCCACAG-3′  3′-CTCGCGCCGTCAGACCATGCTATCTTTAAAATAAATTCCGTGTC-5′ | ChrX:3,373,105..3,373,148 |
| 04 | P35/P36 | 5′-ATCATTCATTCATTGTCTATCGAAAGCGCGGTGGTGGGCTTGGTCGC-3′  3′-TAGTAAGTAAGTAACAGATAGCTTTCGCGCCACCACCCGAACCAGCG-5′ | ChrX:3,382,530..3,382,576 |
| 05 | P37/P38 | 5′-ACAACGATTTCCGCCTTATCTATATTTTTCAGACAGGCATATAACTCAGGAAC-3′  3′-TGTTGCTAAAGGCGGAATAGATATAAAAAGTCTGTCCGTATATTGAGTCCTTG-5′ | ChrX:3,374,908..3,374,960 |
| 06 | **Scm*** (not Scm1/Scm2) | 5′-TGTGTGTGTGTGCGCACGCGCGTGTGTGTGCGTGCATGCGTGCAT-3′  3′-ACACACACACACGCGTGCGCGCACACACACGCACGTACGCACGTA-5′ | N/A |
| 07 | **Pos*** (not Pos1/Pos2) | 5′-TACGATAAGATCAAAGGTGCAGTGCCGCCAGTCGTA-3′  3′-ATGCTATTCTAGTTTCCACGTCACGGCGGTCAGCAT-5′ | N/A |

**Table S1g. Mapping of *Myc* Regulatory Elements, Genomic Coordinates, Oligonucleotide Sequences, Replicate Pairs, and MS**

**Sample Identifiers.**

Oligonucleotide sequences correspond to the sense (5′→3′) strand of annealed double-stranded DNA constructs representing *Myc* regulatory elements (Myc-CRMs) and controls (W-CRM positive control and Scm negative control). Constructs are grouped by replicate pairs (e.g., P1/P2), each consisting of two independent experimental replicates under identical conditions. Each oligo is analyzed using the same sequence with two MS labeling positions (Az at 5′ and Az at 3′), which serve as technical identifiers for traceability in the proteomics dataset. Table S3 contains the corresponding ungrouped MS dataset, including intermediate sample identifiers (e.g., Az3, Az5) for linkage to raw proteomics input files (Table S1f), while Tables S6, S7, and S12 contain the corresponding grouped MS dataset.

**Pos* and Scm***: For simplicity, Pos1/Pos2 and Scm1/Scm2 are designated as Pos and Scm, respectively, throughout data processing and in the manuscript.
