## Supplementary_Data_S2 for "Functional dissection of *Drosophila Myc cis*-regulatory modules (Myc-CRMs) reveals developmentally active DNA-protein interactions"

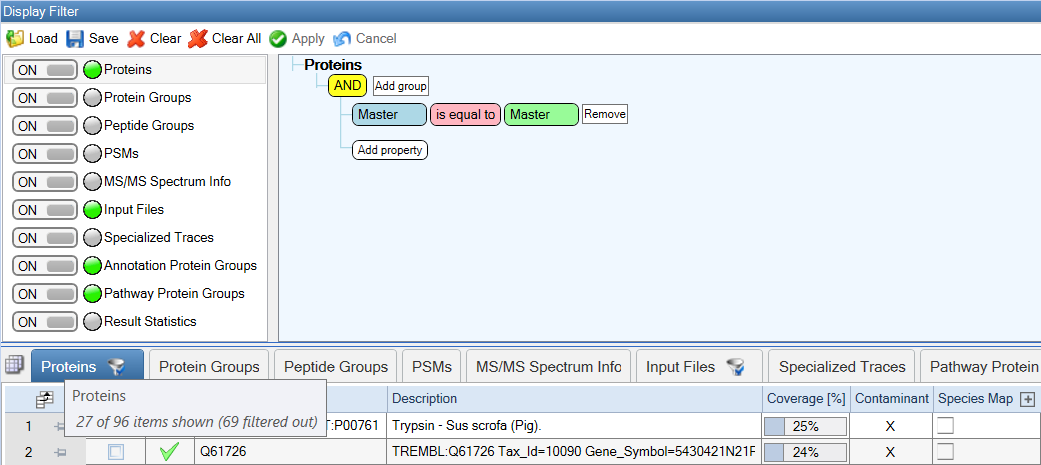

**Fig. S7. Analysis of raw (blank) magnetic beads.**

This figure shows quality control filtering applied to Dynabeads® MyOne™ Carboxylic Acid using the same “Master = Master” filtering criteria as for experimental samples. Twenty-six *Drosophila melanogaster* proteins were detected associated with the raw beads. The complete list of identified proteins is provided in Supplementary Table S2.
