## Supplementary_Data_S3 for "Functional dissection of *Drosophila Myc cis*-regulatory modules (Myc-CRMs) reveals developmentally active DNA-protein interactions"

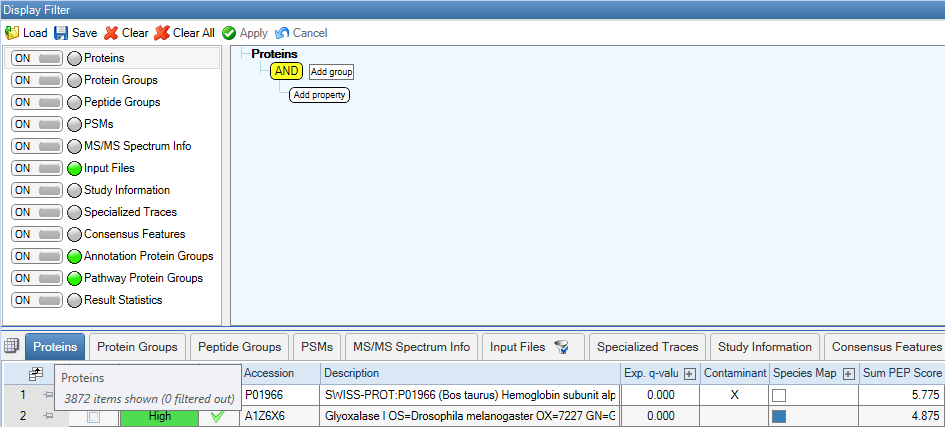

**Fig. S8. Original dataset screenshot.**

No filtering was applied to the dataset. The screenshot shows the original Excel file prior to processing, with biological replicates ungrouped, containing 3,872 proteins. No contaminants were excluded at this stage. The complete protein list is provided in Supplementary Table S3.
