## Supplementary_Data_S4 for "Functional dissection of *Drosophila Myc cis*-regulatory modules (Myc-CRMs) reveals developmentally active DNA-protein interactions"

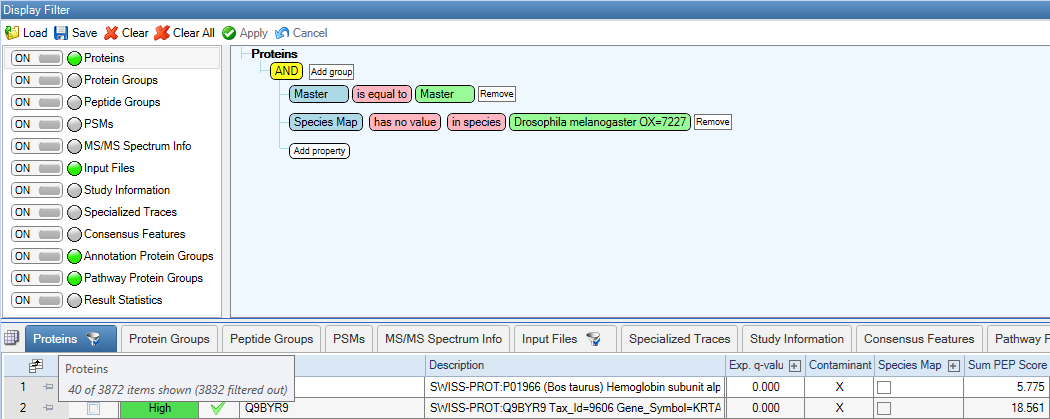

**Fig. S9.** **Applied filter for contaminant exclusion.**

This filtering step removed 40 proteins not belonging to *Drosophila melanogaster*. The complete list of excluded contaminant proteins is provided in Supplementary Table S4.
