## Supplementary_Data_S5 for "Functional dissection of *Drosophila Myc cis*-regulatory modules (Myc-CRMs) reveals developmentally active DNA-protein interactions"

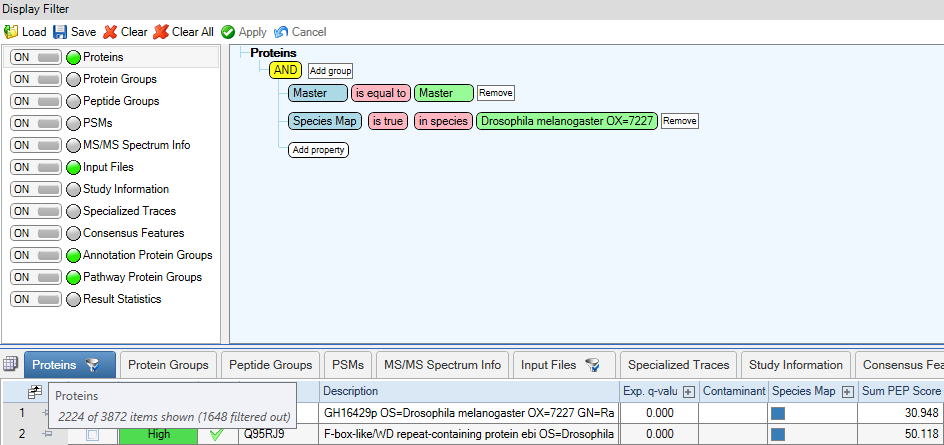

**Fig. S10.** **Application of species filter and contaminant removal in ungrouped replicates.**

A species-specific filter for *Drosophila melanogaster* was applied, removing 1,648 non-*D. melanogaster* proteins and retaining only *D. melanogaster* factors with ungrouped replicates. The complete list of filtered proteins is provided in Supplementary Table S5.
