## Supplementary_Data_S6 for "Functional dissection of *Drosophila Myc cis*-regulatory modules (Myc-CRMs) reveals developmentally active DNA-protein interactions"

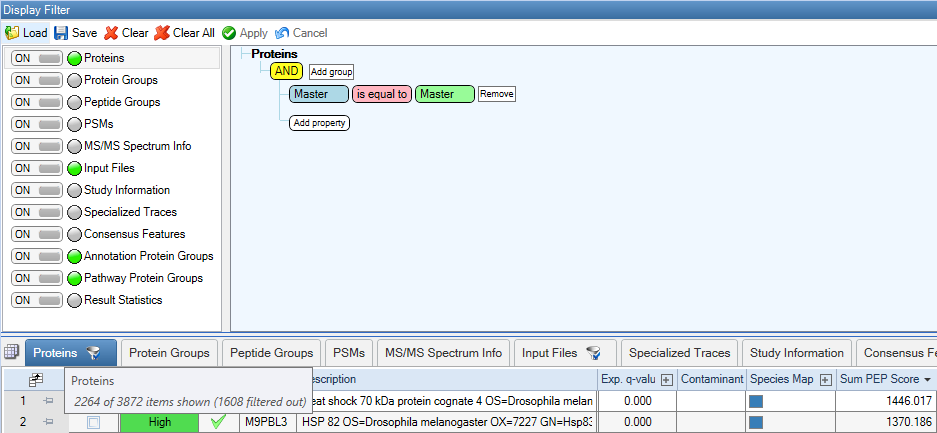

**Fig. S11. Grouping of replicates in the unfiltered proteome.**

Replicates in the original dataset (Supplementary Table S3) were grouped, resulting in the exclusion of 1,608 proteins. The full list of excluded proteins is provided in Supplementary Table S6.
