## Supplementary_Data_S7 for "Functional dissection of *Drosophila Myc cis*-regulatory modules (Myc-CRMs) reveals developmentally active DNA-protein interactions"

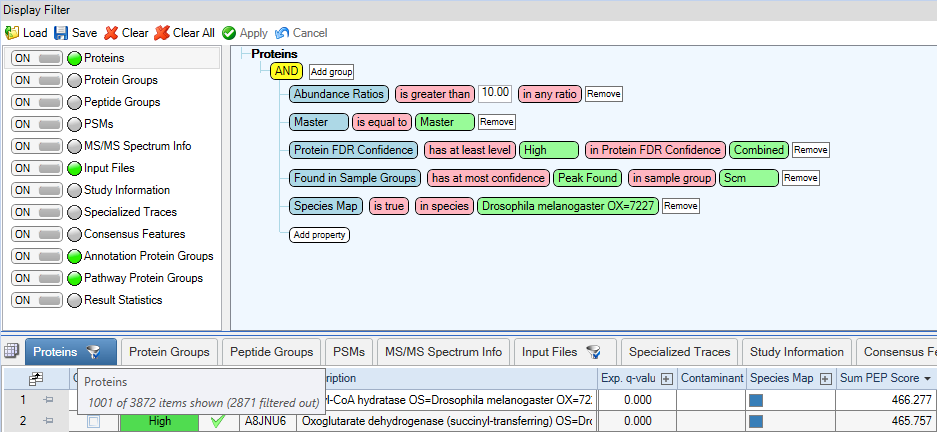

**Fig. S12. Application of quality control filters to unfiltered dataset.**

A series of stringent quality control filters were applied to the unfiltered dataset to identify putative *Myc* regulators. The full list of retained proteins is provided in Supplementary Table S7.
