## Supplementary_Methods_Results_Refs for "Functional dissection of *Drosophila Myc cis*-regulatory modules (Myc-CRMs) reveals developmentally active DNA-protein interactions"

**Article Type: Research Article**

**Title:**

### Contents

- Supplementary Methods
- Supplementary Results
- Supplementary References

### Supplementary Methods

#### S1. Generation of *lacZ* Reporter Fly Strains

Previously described constructs pJ2.1, pJ7, pJ8, pJ8.4, pJ8.5, J2.1, J7, J8, J8.4, J8.5 (Table S1a) and intermediate vectors pCaSpeR4, pCaSpeR4-NLSlacZ, SKII-dmyc5, SKII-In2 were (Kharazmi et al., 2012). New transgenes J7.1–J7.5 were generated by subcloning *Myc* genomic fragments into pCaSpeR4-NLSlacZ upstream of *lacZ*.

The J7 promoter region (ChrX:3,373,200..3,375,005) includes 100 bp upstream of the main TSS, noncoding exon 1, intron 1, and noncoding exon 2 (Table S1a).

J7.1 lacks upstream TATA1-GC1-Inr1 elements (contained in P31/P32); the 1,806 bp genomic 5′-SacII/Acc65I-3′ insert (ChrX:3,373,200..3,375,005) from SKII-dmyc5′ was ligated into SacII/Acc65I-digested, dephosphorylated pCaSpeR4-NLSlacZ.

For J7.2, a 2,164 bp fragment was PCR-amplified from SKII-dmyc5′ with primers J7.2-F (adding SacII) and J7.2-R (Table S1b; 98°C/30 s, 30x [98°C/10 s, 58°C/30 s, 72°C/2 min], 72°C/5 min, and 4 °C hold). The 1,295 bp SacII–Acc65I digest (ChrX:3,373,710..3,375,005; 14 bp exon 1 3′ end, intron 1, noncoding exon 2) was subcloned into SacII/Acc65I-digested, dephosphorylated pCaSpeR4-NLSlacZ vector upstream of the *lacZ* reporter to generate transgene J7.2.

For J7.3, a 641 bp fragment was PCR-amplified from the SKII-dmyc5′ with primers J7.3-F and J7.3-R (Table S1b). The resulting genomic fragment (ChrX:3,373,088..3,373,728) was blunt-ligated (using CIAP, Roche Diagnostics, Indianapolis, IN for vector preparation) into the Acc65I linearized, Klenow-treated, and dephosphorylated pCaSpeR4-NLSlacZ.

For the J7.4 intermediate plasmid, a 1,562 bp fragment was PCR-amplified from the *Myc* genomic sequences within the pJ2.1 enhancer region with primers J7.4-F (adding SacII) and J7.4-R (adding Acc65I; Table S1b). The 1,527 bp SacII-Acc65I digest (ChrX:3,369,648..3,371,174) was subcloned into SacII/Acc65I-digested, dephosphorylated pCaSpeR4-NLSlacZ to obtain the intermediate plasmid pJ7.4-temp. The pJ7 promoter was PCR-amplified with primers J7-F (adding Acc65I) and J7-R (adding Acc65I; Table S1b). The 1,958 bp Acc65I–Acc65I digest was subcloned into Acc65I-digested, dephosphorylated pJ7.4-temp to create the chimeric transgene pJ7.4.

For J7.5 intermediate plasmid, a 932 bp fragment from J8 enhancer (containing 5′-**CGCGTGGGAAAA**-3′ with Hairless (*H*) site **GTGGGAA**; ChrX: 3,379,324..3,380,254) was amplified with In2E-F-Not/In2E-R-Asp (Tables S1a, S1b), yielding NotI–Acc65I insert (ChrX:3,379,324..3,380,255). It replaced full intron 2 in J8 to create pJ7.5-temp. The pJ7 promoter (from Acc65I-digested pJ7.4) was ligated into Acc65I-digested pJ7.5-temp downstream of the enhancer.

For J7.5, a 932 bp fragment from J8 enhancer (containing 5′-**CGCGTGGGAAAA**-3′ with Hairless (*H*) site **GTGGGAA**) was amplified with In2E-F-Not/In2E-R-Asp (Tables S1b, S1d), yielding NotI–Acc65I insert (ChrX:3,379,324..3,380,255). It replaced full intron 2 in J8 to create pJ7.5-temp. The pJ7 promoter (from Acc65I-digested pJ7.4) was ligated into Acc65I-digested, dephosphorylated pJ7.5-temp downstream of the enhancer region.

Protocols for transfections, plasmid prep, ϕC31 integrase, and P-element transgenesis are in (Kharazmi et al., 2012). Transgenes J2.1, J7, J7.1–J7.5, J8, J8.4, J8.5 were sequenced distally (pcaF, Acc651_F) and proximally (pCAR, pZR) (Table S1b), oriented toward *Myc*, by Microsynth AG (Balgach, Switzerland), where the oligos were also synthesized.

### S2. *D. melanogaster* Stocks and X-Gal Staining Assays

Transgenic lines J2.1, J8, J8.4, and J8.5 were generated by random P-element transgenesis or ϕC31 integrase-mediated integration (Kharazmi et al., 2012). Lines J7.1–J7.5 were produced by random P-element insertion into *y[1] w[1118]* embryos (Rainbow Transgenic Flies Inc., CA). To account for position effects associated with random P-element insertion, multiple independent transgenic lines were established and analyzed for each J7 construct. Between 6 and 12 independent insertion lines were recovered per construct, of which 5–10 lines were subjected to X-gal staining. Reporter expression patterns were largely consistent across the majority of lines for each construct, with 4–7 lines per construct exhibiting reproducible staining patterns (see Table S1c for details). The *dpp-lacZ* stock (Kharazmi et al., 2012) served as positive control, and *w[1118]* as negative control (Table S1c).

All stocks were maintained on standard medium at 25°C; crosses were performed at the same temperature. Larval and adult female tissues were dissected from homozygous F2 progeny for *lacZ* analysis.

**X-Gal Staining Protocol:**

Homozygous F2 tissues were fixed in 0.2% glutaraldehyde for 20 min at room temperature.

Rinsed in PBS and incubated in X-Gal solution (1 mg/mL X-Gal, 5 mM $K_3Fe(CN)_6$, 5 mM $K_4Fe(CN)_6$, 2 mM $MgCl_2$, 0.01% sodium deoxycholate, 0.02% NP-40) at 37°C for 6–12 h.

Washed in PBS, post-fixed in 4% paraformaldehyde, and mounted for imaging.

Transgenic lines J2.1, J8, J8.4, and J8.5 were generated by both the random P-element and site-directed ϕC31 integrase–mediated transformation (Kharazmi et al., 2012). Lines J7.1–J7.5 were produced by P-element insertion into *y[1] w[1118]* embryos (Rainbow Transgenic Flies Inc., CA). *dpp-lacZ* and *w[1118]* served as positive and negative controls, respectively (Table S1c).

Flies and crosses were maintained on standard medium at 25 °C. X-Gal staining of *Myc-lacZ* lines J2.1, J8, J8.4, J8.5, and J7-J7.5 truncations from homozygous F2 tissues was performed using modified protocols based on previously described methods (Song et al., 2004; Kharazmi et al., 2012; Kharazmi and Moshfegh, 2013).

### S3. Protein Extraction from *Drosophila* Embryos

Wild-type *Canton-S* flies (BDSC #64349; Kyoto #1; Table S1c) were maintained at 25°C and 75-80% humidity on yeast-supplemented food in population cages.

Embryos were collected over 0-12h after egg laying (AEL) for up to 72h, stored at 4 °C, manually dechorionated, and suspended in cold **buffer I** (3ml/ g embryos: 15 mM HEPES-$K^+$ pH 7.6, 10 mM KCl, 5 mM $MgCl_2$, 0.1 mM EDTA, 0.5 mM EGTA, 350 mM sucrose, 1 mM DTT, 1 mM $Na_2S_2O_4$, 0.2 mM PMSF, 1 mM benzamidine-HCl). Embryos were homogenized on ice using a 40 mL Dounce homogenizer.

Filtered and centrifuged (8,000 rpm, 12 min, 4°C), supernatants were collected as soluble cytoplasmic fraction (SCF) and stored at -80 °C.

Nuclear pellet re-homogenized in buffer I, re-pelleted, and resuspended in **HEMG/0.1 M KCl buffer** (0.5 ml/g nuclei: 25 mM HEPES-$K^+$ pH 7.6, 0.1 mM KCl, 12.5 mM $MgCl_2$, 0.1 mM EDTA, 20% glycerol, 1.5 mM DTT, 1 mM $Na_2S_2O_4$, 0.1 mM PMSF fluoride, 1 mM benzamidine-HCl).

Incubated on ice 20–60 min with occasional swirling and shaking, ultracentrifuged (SW28 rotor, 24,000 rpm, 1 h, 4°C).
Uppermost lipid layer was removed, and the soluble nuclear fraction (SNF) underneath was collected, aliquoted, frozen in liquid $N_2$ at −80°C.
Protein levels were quantified ((BCA assay, Pierce™)) and confirmed via 10% SDS-PAGE, Coomassie staining, and infrared imaging at 700 nm using the Odyssey Classic system (Luo et al., 2006; Harris et al., 2007).

**S4. Oligonucleotides Comprising Myc-CRMs and Controls and Annealing to Double-stranded**

To visualize double-stranded (ds) oligonucleotides, one strand was labeled at either the 5′- or 3′-end with IRDye Dyomics 781; complementary strands were labeled oppositely so that fluorophores were juxtaposed after annealing. To minimize steric hindrance during protein binding to Magnetic Bead–$PEG_{(23)}$–DNA heterotrimers, each oligonucleotide pair was synthesized in two forms, containing an azide–NHS ester group at either the 5´- or 3´-end of one strand; these two forms also served as technical replicates for MS-based protein identification. Oligonucleotides were synthesized by Microsynth AG, Switzerland.
Annealing reactions contained complementary oligonucleotides at a total concentration of 1 pmol/µl, mixed at a 1:1 molar ratio in 50 mM NaCl. Thermal cycler comprised one cycle 95°C for 5 min; 40 cycles decreasing 1 °C per cycle (1 min each); 55 °C for 30 min; 20 cycles decreasing from 55 °C at 1 °C per cycle (1 min each); and 4 °C hold. The number of hybridization cycles was adjusted according to $T_m$ (protocol described is for a primer pair with $T_m$ 55 °C in 50 mM NaCl, pH 8.0).

**S5. Electrophoretic Mobility Shift Assay (EMSA) Protocol Using dsAzide-Dyo781 Oligonucleotides**

EMSA reactions (20 µl) contained 1.35 pmol of dsAzide-Dyomics 781-labeled oligos (Table S1e), 2 µl 10x binding buffer (100 mM Tris, 500 mM KCl, 10 mM DTT, pH 7.5), 2 µl 25 mM DTT with 2.5% Tween-20, 1 µl poly(dI.dc) (1 µl in 10 mM Tris, 1 mM EDTA, pH 7.5), 1 µl 100 mM $MgCl_2$, 1 µl 1% NP-40, 200 µg protein (SCF or SNF), and ultrapure water to volume. Reactions were incubated in the dark for 30 min.
After adding 2 µl 10x Orange Loading Dye (Odyssey EMSA Kit), samples and PageRuler protein marker were resolved on 5% native TBE-PAGE gels for 1 h at 80 V using the Mini-PROTEAN system. Wet gels were imaged at 800 nm on an Odyssey Classic Infrared Imager, and images were processed by adjusting brightness and contrast in Adobe Photoshop CS6.

**S6. Creation of Magnetic Bead–$PEG_{(23)}$–DNA Constructs**

Magnetic beads (Dynabeads®) were washed twice in 100 mM MES buffer (pH 6.0) and resuspended at 10 µg/µL. EDC (N-Ethyl-N'-(3-dimethylaminopropyl) carbodiimide hydrochloride, MW 191.7) (Thermo Fisher, cat.# 77149) activation and amide coupling were performed by mixing 140 µL beads with 84 µL 1250 mM EDC (75 nmol/µg bead) and 42 µL 100 mM DBCO–$PEG_{(23)}$–amine (3 nmol/µg bead) in PBS/0.1% Triton X-100 (final 300 µL). The reaction proceeded overnight (12–16 h) at room temperature in the dark with gentle rotation (7 rpm) to yield DBCO–$PEG_{(23)}$–coated beads. Beads were washed 3x in storage buffer (100 mM PBS, 0.15 M NaCl, pH 7.5, 0.1% Tween-20), quenched with 50 mM Tris (pH 7.4)/0.01% Tween-20 (>30 min) to remove unbound carboxylic sites on beads, washed again 3x, and stored at 10 µg/µL in storage buffer (4°C, <1 week).

Copper-free click chemistry reaction, a highly efficient strain-promoted azide–alkyne cycloaddition between DBCO and azide groups, was used to covalently attach azide-modified double-stranded DNA oligonucleotide. 140 µL DBCO–$PEG_{(23)}$–Beads were reacted with 7 µL dsDNA (0.45 pmol/µL) in 300 µL storage buffer for 4.5 h at room temperature (dark, 7 rpm). The resulting Bead–$PEG_{(23)}$–DNA construct was washed 4x in storage buffer, resuspended in 500 µL, and stored at 4°C (<2 days) until use in protein binding experiments.

**S7. Protein Binding to Magnetic Bead–$PEG_{(23)}$–DNA Constructs and Sample Preparation for LC-MS/MS**

The magnetic Bead–$PEG_{(23)}$–DNA construct in storage buffer was carefully resuspended. After placement of the tube on a magnetic rack, the supernatant was aspirated. The beads were then rinsed once with storage buffer and resuspended in 140 µL of the same buffer to a final concentration of 10 µg/µL. We set up protein binding reactions for the soluble cytoplasmic fraction (SCF) and soluble nuclear fraction (SNF) samples using components from the Odyssey® Infrared EMSA Kit (Part #829-07910) with the following components per reaction:

**a)** Magnetic Bead–$PEG_{(23)}$–DNA construct (10 µg/ µL): **1400 µg**; **b)** 10x binding buffer: **10 µL**; **c)** 25 mM DTT, 2.5% Tween®- 20: **10 µL**; **c)** poly(dI.dc), 1 µg/ µL: **5 µL**; **d)** 1% NP-40: **5 µL**; **e)** 100 mM MgCl2: **5 µL**; **f)** Protein (SCF in cytoplasmic samples and SNF in nuclear samples): **200 µg**; **g)** ultrapure water was added to a final volume of **100 µL**. Sample tubes were incubated in the dark for 30 minutes by occasional manual flipping to keep the mixture in suspension. After incubation and aspiration of the supernatants, the Bead–$PEG_{(23)}$–DNA/protein complexes were gently washed four times for 5 min with storage buffer, and residual liquid was removed once more.

Bead–$PEG_{(23)}$–DNA/protein samples were stored at -80°C until processed by liquid chromatography–tandem mass spectrometry (LC-MS/MS)–based identification of DNA-associated proteins.

**S8. Sample Processing and Protein Identification by LC-MS/MS**

**Sample Preparation:**

1 Beads (unmodified and Bead–$PEG_{(23)}$-DNA/protein complexes) were resuspended in 8 M urea, 100 mM ammonium bicarbonate with 5 mM DTT and incubated for 1 h at room temperature.

2 Cysteines were alkylated with 15 mM iodoacetamide for 1 h.

3 Samples were diluted eightfold with 100 mM ammonium bicarbonate, centrifuged at 20,000 × g for 5 min, and supernatants were digested with 0.2 µg trypsin for 16 h at 37°C.

4 Tryptic peptides were isolated and desalted by reverse-phase micropurification.

5 Samples included Myc-CRMs, SCF and SNF, scrambled oligo (negative control, Scm), W-CRM (positive control), and raw beads to identify contaminants (Table S1f).

6 Peptides were dissolved in 0.1% formic acid, loaded onto a 2 cm × 100 µm trap column, and separated on a 15 cm × 75 µm C18 column (ReproSil-Pur C18-AQ, 3 µm).

7 Peptides were eluted at 250 nL/min over a 30 min gradient from 5% to 35% phase B (0.1% formic acid, 90% acetonitrile).

**LC-MS/MS:**

1 MS analysis was performed on a Thermo Scientific Orbitrap Eclipse Tribrid mass spectrometer coupled to an EASY-nLC 1200 system.

2 Raw data were processed using Proteome Discoverer 2.5 and searched against the *Drosophila melanogaster* proteome and a contaminant database.
3 Search parameters included trypsin specificity, minimum peptide length of 6 residues, precursor mass tolerance of 10 ppm, fragment mass tolerance of 0.02 Da, and dynamic modifications of methionine oxidation and N-terminal acetylation.
4 Precursor ion quantification was based on summed abundances of unique peptides without normalization.

**S9. Proteomic Data Processing and Analysis**

Raw LC–MS/MS data were processed using Proteome Discoverer™ v2.5 (Thermo Fisher Scientific) and searched against the *Drosophila melanogaster* reference proteome, supplemented with a contaminant database. Peptide-spectrum matches were filtered at a false discovery rate (FDR) <1% at peptide and protein levels. Protein abundance was quantified using unique peptides, and values were normalized across samples within Proteome Discoverer prior to downstream analysis.

Filtered and normalized datasets were imported into ComplexBrowser (Michalak et al., 2019; https://computproteomics.bmb.sdu.dk/app_direct/ComplexBrowser/) for quality control and protein complex–level analysis. QC metrics included normalized log2 abundance distributions, coefficient of variation (CV) across biological replicates, and pairwise Pearson correlation coefficients between all sample pairs. Analyses included negative control (Scm), experimental conditions (Myc-CRMs), and positive control samples.

Volcano plot analysis was performed using VolcaNoseR (Goedhart and Luijsterburg, 2020; https://huygens.science.uva.nl/VolcaNoseR2/). Input data consisted of log2 abundance ratios and corresponding q-values. Significance thresholds were set at $q < 0.01$ and absolute abundance ratio > 2.

Protein complex annotation and interpretation were performed by integrating ComplexBrowser outputs with manual curation using UniProt, FlyBase, STRING, and The Interactive Fly.

**For supplementary methods:**

Quality control and exploratory data analysis were performed in ComplexBrowser (Michalak et al., 2019; https://computproteomics.bmb.sdu.dk/app_direct/ComplexBrowser/). This included generation of normalized log2 abundance boxplots, coefficient of variation (CV) histograms, and pairwise scatter plots with Pearson correlation coefficients to assess reproducibility between biological replicates. Analyses were performed across all sample pairs, including negative control (Scm), experimental conditions (Myc-CRMs), and positive control samples.

### Supplementary Results

*Supplementary Results S3.1. Reporter Constructs and Detailed Expression Patterns*

To establish the functional relevance of *Myc cis*-regulatory modules (Myc-CRMs), we previously analyzed the reporter activity of the J2.1 and J7 transgenes. J2.1, containing 7.2 kb of upstream regulatory sequence, drives *Myc* expression in larval and adult female tissues, whereas the smallest J7 construct (1.928 kb; ChrX:3,373,008..3,375,005) activates reporter expression in embryos and ovaries (Fig. 1A–B; Table S1a). These constructs provided the baseline framework for subsequent dissection of *Myc* regulatory elements.

Building on this prior work, a series of truncations was generated in the present study to define the contribution of individual *cis*-regulatory modules (Fig. 1C–E; J7.1–J7.3). Removal of the P1 promoter region harboring the Inr–TATA–box-containing Myc-CRM P31/P32 in J7.1, and further truncation of sequences downstream of P31/P32 in J7.2—while retaining P37/P38 in both constructs—did not alter *Myc-lacZ* expression in embryos and ovaries in these three transgenes (J7, J7.1, J7.2). In contrast, removal of intron 1 and noncoding exon 2 sequences including Myc-CRM P37/P38 in J7.3—while retaining the upstream sequences containing the Myc-CRM P31/P32—abolished expression in adult female tissues (Fig. 1E). These results indicate that endogenous *Myc* expression in adult female tissues operates independently of P1 promoter activity and that the P37/P38 enhancer—a 53-bp sequence (ChrX:3,374,908..3,374,960; Table S1d and S1g)—is sufficient for the endogenous *Myc* patterning in embryos and ovarian cells during early developmental stages.

Consistent with recent studies, CRMs such as P37/P38 may function not only as classical enhancers but also as transcriptional initiators through context-dependent recruitment of RNA Pol II and chromatin-modifying factors (Barral and Dejardin, 2023; Lindhorst and Halfon, 2023).

To test whether the activity of the upstream Myc-CRM P31/P32 could be restored, we added this element in-frame to the J7 promoter, generating J7.4. This construct restored reporter expression in larval and adult female tissues, although activity in eye and wing imaginal discs was reduced compared with J2.1 (Fig. 1F). This result suggests that the proper enhancer–promoter distance is required for strong transcriptional activation and that the P1/P2 element functions as a late developmental enhancer, active during the onset of differentiation in larval primordial tissues.

Previous work analyzed the 8 kb full-length intron 2 sequences of the *Myc* gene, which comprises a dense subset of conserved *cis*-regulatory modules (CRMs), positioned upstream of an Inr–DPE core promoter (Kharazmi et al., 2012; Kharazmi and Moshfegh, 2013). The full-length J8 transgene, containing both the upstream CRM-rich regulatory region and the downstream Inr–DPE promoter, faithfully reproduced endogenous *Myc* expression in the tested larval and adult female tissues. In contrast, truncations in which either the upstream regulatory region or the Inr–DPE core promoter was tested in isolation (J8.4 and J8.5, respectively) failed to drive *Myc-lacZ* expression, indicating that both elements are required for productive transcriptional activity.

In this study, we focused on a single conserved CRM within the J8 regulatory region (P29/P30), selected based on sequence conservation and features suggestive of transcription factor binding. We first asked whether this CRM is sufficient to recapitulate the *Myc* expression pattern observed with the full-length J8 transgene. In addition, we tested whether its activity is dependent on an Inr–DPE core promoter or whether it can also function with a TATA-containing promoter.

To address these questions, we fused the P29/P30 CRM upstream of the J7 TATA-containing core promoter, generating a composite construct (J7.5). This configuration restored reporter expression in all tested tissues, demonstrating that P29/P30 is sufficient to recapitulate J8-like *Myc* expression and can function with either Inr–DPE or Inr–TATA-containing core promoters, revealing both promoter-dependent and promoter-flexible properties of *Myc cis*-regulatory modules.

Tissue abbreviations: a, brain; b, wing disc; c, eye-antennal disc; d, leg disc; e-f, embryos; g, ovary. β-Gal staining was performed overnight at 25°C on third-instar larval and adult female tissues. Scale bar in (a–g) indicates 100 μm. Sequences and coordinates of the Myc-CRMs P1/P2. P29/P30, P31/P32, P35/P36, and P37/P38 are provided in Tables S1d, S1e, and S1g. For β-galactosidase staining of tissues from positive and negative control flies, see Fig. S1a–g in Supplementary Material.

*S3.2. Quality control of oligonucleotides and protein extracts for Myc regulator enrichment*

All *Myc* target sequences, control elements, and the scrambled oligonucleotide were annealed to generate double-stranded DNA probes (Table S1g). Probe integrity and duplex formation were assessed by native polyacrylamide gel electrophoresis, revealing discrete bands of the expected mobility for all oligonucleotides (Fig. 3A–C).

Annealed probes were pegylated via click chemistry reaction to enable magnetic bead (Shchepinov et al., 1997)–based enrichment (Fig. 3D). PEG modification was verified by a reproducible mobility shift on native gels, with pegylated probes migrating more slowly than unmodified DNA (Fig. 3E). Electrophoretic mobility shift assays using the positive control confirmed that PEG conjugation did not impair protein-binding efficiency (Fig. 3F).

DNA–protein binding conditions were optimized by titrating increasing amounts of embryonic protein extract at constant oligonucleotide concentration. Both soluble cytoplasmic (SCF) and nuclear (SNF) fractions exhibited increased formation of shifted and super-shifted bands, with near-complete complex formation at 2.25 pmol DNA for the positive control (Fig. 3G, lanes 2–9). *Myc* targets such as the TATA-CorePromoter (P31/P32) probe displayed a greater proportion of unbound DNA at equivalent probe mass (Fig. 3H, lanes 2–7), consistent with reduced binding efficiency. Based on these analyses, 200 μg of SCF or SNF was selected for subsequent protein identification experiments (Methods, Section 2.5).

Titration experiments using the Oocyte Element probe (P37/P38) confirmed comparable trends: decreasing SCF and SNF concentrations reduced super-shift intensity. With SCF, no free oligo was observed, suggesting complete binding (Fig. 3I, lanes 1–3). In contrast, SNF showed residual unbound DNA at lower protein concentrations (Fig. 3I, lanes 4–6). These results indicate 0.45 pmol oligo is sufficient for specific protein complex formation.

Binding specificity was evaluated using the Late-Enhancer probe and scrambled control sequence (Scm). SCF produced shifted complexes with both probes (Fig. 3J, lanes 2–5), indicating nonspecific binding. In contrast, SNF showed concentration-dependent binding to the Late-Enhancer probe, with no binding to the scrambled control (Fig. 3J, lanes 6–9), consistent with sequence-specific interactions mediated by nuclear proteins. Specificity was further confirmed by competition electrophoretic mobility shift assays, in which an IR-labeled Late-Enhancer probe was challenged with increasing molar excess of an unlabeled competitor of identical sequence (Primer pair P3/P4; Table S1d). Increasing competitor concentrations resulted in progressive loss of the labeled probe signal, with complete competition observed at 30-fold molar excess (Fig. 3K, lanes 5–6).

Based on these controls, SNF—enriched in transcription factors—was selected as the input for all protein identifications. The scrambled oligonucleotide and W-CRM element (Schweizer et al., 2003; Archbold et al., 2014) were used as negative and positive controls, respectively, throughout the study.

Finally, Magnetic Bead–PEG(23)–DNA constructs were generated for all Myc-CRM targets and control probes as described in Methods. Infrared-labeled oligonucleotides in these constructs (e.g., Oocyte Element P37/P38) were visualized using a LI-COR infrared imager (Fig. 3L, panels 1a-1d and 2a-2d) prior to protein binding to confirm successful bead conjugation and probe integrity. Each validated construct was subsequently used for isolation of DNA-bound protein complexes for mass spectrometry analysis (Fig. S2A–D).

#### *S3.3. Magnetic Solid Surface Enrichment of Myc Regulators: Detailed Workflow and Assay Conditions*

DNA–protein interactions occur via sequence-specific or non-specific binding and are stabilized by $H^+$ bonds, low ionic strength, appropriate pH, and divalent cations such as $Zn^{2+}$ and $Mg^{2+}$ (Phizicky and Fields, 1995; Dey et al., 2012; Nilkanta and Bagchi, 2018). To preserve native interactions, assays were performed under non-denaturing conditions. Both soluble nuclear (SNF) and cytoplasmic (SCF) fractions were initially evaluated as protein sources. Non-specific binding observed with SCF led to the selection of SNF, enriched in transcription factors, for all subsequent MS-based discovery experiments.

Prior to incubation with *Myc cis*-regulatory modules (CRMs), increased SNF concentrations were assessed for quality and quantity on non-denaturing 5% PAGE gels run in low-salt 0.5x TBE buffer. Coomassie staining revealed a graded protein smear consistent with increasing input (Fig. S2E, lanes 1–9). In-gel digestion following gel-based separation often results in co-migrating background proteins, reducing sensitivity and selectivity in mass spectrometry (Goodman et al., 2018).

To overcome these limitations, a magnetic solid surface enrichment protocol (MSSEP) was developed and used Carboxylic Magnetic Bead–$PEG_{(23)}$–DNA constructs (Fig. 4; Methods 2.5). Amine-$PEG_{(23)}$-DBCO was covalently linked to carbodiimide-activated Dynabeads® MyOne™ Carboxylic Acid beads; the hydrophilic PEG spacer increases solubility, reduces steric hindrance, and spaces DNA from the bead to facilitate protein–DNA interactions (Shchepinov et al., 1997). Copper-free click chemistry was used to covalently conjugate the DBCO-modified beads to azide-functionalized DNA oligonucleotides (Jewett and Bertozzi, 2010; Eeftens et al., 2015).

DNA-associated proteins were immobilized, washed, and recovered by biomagnetic separation from *Drosophila* nuclear crude extracts. This highly selective fraction of proteins, associated with Myc-CRMs and enriched in transcription factors and other regulatory proteins, reduces background noise, increases sensitivity in mass spectrometric analyses, and delivers datasets with high-confidence. The MSSEP protocol is readily adaptable to other experimental contexts, including antigen–antibody detection or protein–RNA/microRNA interactions, by substituting the DNA oligonucleotide with appropriate RNA sequences or structural motifs. Detailed buffers, concentrations, incubation times, and replicates are provided in the Materials and Methods section (Methods 2.6).

he MSSEP protocol is readily adaptable to other experimental contexts, including antigen–antibody detection or

#### *S3.4. MS analysis of raw beads, controls, and dataset filtering*

The exact same data analysis was performed for the blank sample (raw beads), both control samples "the Scrambled (Scm) oligonucleotide" (Negative Control), the positive control (Pos), and the Myc-CRM samples. Analysis of blank beads identified 26 D. melanogaster proteins (Supplementary Data S2), 12 of which showed low scores with low-confidence identifications, and 10 were based on a single unique peptide, which would normally be excluded prior to publication. Three proteins (actin, albumin, and ATPase) were classified as non-*Drosophila* contaminants, and collagen alpha I(III) (P04258; 138.4 kDa, 1466 aa) was detected exclusively in blank bead samples, consistent with leaching from bead coatings during sample preparation (e.g., boiling). Bead-derived contaminants were independently assessed by boiling beads in 8 M urea followed by SDS–PAGE, revealing faint protein bands under UV illumination (Fig. S2F). These contaminants were computationally filtered prior to downstream analysis, demonstrating that raw beads can be effectively used for biomagnetic separation and affinity purification of proteins and nucleic acids.

The initial MS dataset (replicates ungrouped) contained 3,872 proteins (Supplementary Data S3). Following removal of bead-associated proteins (Supplementary Data S4) and filtering for *Drosophila*-specific identifications, 2,224 proteins were retained using ungrouped replicate data required for ComplexBrowser analysis (Supplementary Data S5), or 2,264 proteins when grouped replicates were analyzed (Supplementary Data S6). The grouped-replicate dataset was subjected to further quality filtering, yielding 1,001 proteins enriched on Myc-CRM targets (Supplementary Data S7), which were subsequently examined to identify *Myc* regulators, with validation provided by enrichment of known W-CRM regulators in the positive control.

#### *S3.5. Heat-map clustering and functional interpretation*

Heat-map analysis clustered proteins by enhancer or promoter activity, reflecting differences in binding specificity, affinity, and abundance relative to the starting nuclear extract (SNF). *Myc* targets and the positive control segregated by functional class (promoters, early enhancers, late enhancers), consistent with overlapping protein recruitment by similar *cis*-regulatory modules.

The *Myc* distal Late-Enhancer (P1/P2) clustered with the positive control dTCF/Pan, consistent with loss of P1/P2 abolishing *Myc* expression in larval imaginal tissues and brain (Kharazmi et al., 2012; Fig. 1B) and supporting a role in larval development and organ morphogenesis.

The SNF fraction (0–72 h nuclear embryonic extract) is a subset containing broad protein assemblies—lineage-specific transcription factors, co-activators, chromatin remodelers, and mediators—recruited by late differentiation enhancers (Spitz and Furlong, 2012). Under abundance-correlated label-free quantification, Late-Enhancer and positive control protein profile intensities align closely with SNF, explaining their clustering/proximity. The Scrambled control (Scm) clustered near promoters, which primarily bind general transcription factors with limited stage-specific occupancy in the extract. TATA- (P31/P32) and DPE-CorePromoter (P35/P36) clustered near the scrambled negative control, reflecting shared low-enrichment signals from general machinery and background/non-specific binding (often abundant "sticky" proteins like histones and chaperones) (Mellacheruvu et al., 2013).

The DPE-Enhancer (P29/P30) and Oocyte Element (P37/P38) clustered together. P37/P38 is sufficient for *Myc* expression in ovaries and embryos (Figs. 1C–D), whereas P29/P30 is required for DPE promoter activity (Fig. 2A and 2C). Their shared profiles suggest that the two early enhancer modules may support rapid *Myc* mRNA production during early embryonic proliferation, in contrast to Late-Enhancer, which activates *Myc* predominantly in lineage-specific manner during morphogenesis.

*S3.6. Functional Classification of Proteins Associated with W-CRM and Myc-CRMs*

Following quality control and clustering of functionally related samples (Figs. 5–8), proteins were grouped by CRM class to facilitate comparative analysis across *Myc* targets. The analyzed groups included the positive control (W-CRM); Late-Enhancer constructs (P1/P2); TATA-containing core promoters (P31/P32); DPE-containing core promoters (P35/P36); and strong developmental enhancers such as the Oocyte Element (P37/P38) and the downstream DPE-Enhancer (P29/P30).

This classification distinguished proteins unique to individual *Myc* targets or CRM classes from those shared across multiple targets (Tables S9a–S9h). In addition to well-characterized *Myc*-associated factors, the dataset revealed several newly recognized or poorly characterized proteins that represent promising candidates for future functional validation and structural studies.

*S3.7. Additional Factors and Details on Regulatory Factors Associated with the Positive Control W-CRM*

***CKIIα***: modulates Hedgehog/Wnt during proliferation and patterning (Jia et al., 2010).
Snx6: retromer component, positive regulator of Wnt secretion (Zhang et al., 2011).
***Galla-1***: CGX complex member, essential for mitosis, proliferation, and polarity in wing discs (Yeom et al., 2015).
**Cul1**: SCF ubiquitin ligase adapter, negatively regulates Wnt and controls cell cycle (Roberts et al., 2012).
***Twins (tws)***: PP2A component, stabilizes Arm/TCF during Wnt transcription, influences wing disc cell fate (Bajpai et al., 2004).
***Atu***: involved in Wg transactivation, Pol II initiation/elongation, and hemocyte proliferation (Sinenko et al., 2010).
***Rap1***: regulates planar cell polarity, morphogenesis, imaginal disc patterning, and follicle migration (Chang et al., 2018).

Signaling Components:
***Mad*** (BMP/Dpp effector): regulates cell cycle arrest, morphogenesis; represses transcription via Schnurri/Medea complexes; intersects with Wg in wing disc patterning/boundary formation (Sekelsky et al., 1995; Pyrowolakis et al., 2004; Zeng et al., 2008).
***Sumo (smt3)***: post-translationally modifies chromatin proteins; modulates Dpp/Hh signaling; controls transcriptional silencing at compartment boundaries (Miles et al., 2008; Lv et al., 2016; Jox et al., 2017; Cappadocia and Lima, 2018).
***Misshapen/Msn (MAP4K)***: integrates Wnt/Wg cues for cytoskeletal dynamics, epithelial organization (Lewellyn et al., 2013).
***Mlf and Su(fu)***: Mlf modulates transcription through interactions with DNA-bound transcription factors and chaperones, while the Hedgehog regulator Su(fu) contributes to morphogenesis, proliferation, and hematopoiesis.

Chromatin-associated and remodeling factors:
***Wapl***: controls sister chromatid cohesion and chromatin organization; modulates Wnt during wing development (Verni et al., 2000; Cho et al., 2013).
***NuRF components (NURF-38, Iswi)***: recruitment via Arm; Iswi represses transcription via TCF motifs (Badenhorst et al., 2002).
***SWI/SNF (Brahma) components (Snr1, Moira)***: remodel chromatin, contribute to morphogenesis and interact with Wnt signaling (Collins and Treisman, 2000; Parker et al., 2008).
***Cdc73/PAF1 (atms, Atu)***: mediates histone modifications and Pol II transcription; Atu participates in Wnt transactivation (Adelman et al., 2006).

**ECM and transport proteins:**
***Perlecan (trol)***: heparan sulfate proteoglycan; modulates Wnt, Hh, TGF-β signaling, linked to wg expression (Park et al., 2003; Schneider et al., 2006; Lindner et al., 2007).
***AP-2μ (CG7057)***: ortholog of AP2M1; potential role in Frizzled trafficking (Yu et al., 2007).
***Vps26***: retromer component; maintains Wg/Dpp gradients (Port et al., 2008).
***Pmm2***: glycosylation-related; influences wing positioning via Dlp, Wg, and Mmp2 (Parkinson et al., 2016).

These enrichments support biologically relevant interactions with wingless/wg regulatory networks.

*S3.10. Extended analysis of Pol II– and chromatin-associated factors at Myc-CRMs*

***RNA Pol II and TFIID complexes***: RNA Pol II largest subunit A (RPII215/RPB1), functions as a structural scaffold for the assembly of basal transcription factors and is essential for transcription initiation and elongation (Brickey and Greenleaf, 1995). Subunit B (Polr2B/RpII140/RPB2) is required for Pol II transcription at nearly all transcription start sites, underscoring its central role in promoter recognition and transcriptional activity (Hamilton et al., 1993). Subunit C (Polr2C/RPB3) has been implicated in stress-responsive transcription, including activation of Hsp70 expression during heat shock (Yao et al., 2007). Cabeza/dFUS (caz) the sole *Drosophila* homolog of human TLS/FUS and a TFIID-associated factor binds ssDNA/RNA and integrates transcription initiation with mRNA processing. Additionally, Cabeza as a member of TET/FET protein family functions in transcription regulation during development and disease (Chau et al., 2016). Through interactions with N-Myc downstream regulated gene-1 (NDRG1) and the AT-hook transcription factor Xrp1, Cabeza may form specialized TFIID subcomplexes that regulate *Myc* transcription in response to specific activators and TAFs (Li et al., 2023).
***TRL/GAF***: The Trithorax-like GAGA factor (TRL/GAF) recruits Polycomb group (PcG) complexes to mediate chromatin remodeling. Both trithorax group (TrxG) and PcG proteins are essential for chromatin-based regulation of developmental transcription, including the expression of homeotic genes (Matharu et al., 2010). TRL/GAF is also required for follicular cell morphogenesis and germline differentiation, and mutations in Trl cause female sterility (Ogirenko et al., 2008).
***Ssb-c31a***: a single-stranded DNA binding protein was identified at key *cis*-regulatory modules—the TATA- and DPE-CorePromoters, DPE-Enhancer, and Oocyte Element—each representing a functional transcriptional initiation site, with the two DPE elements acting cooperatively. Ssb-c31a binds sequence-specifically to the NssBF

element in the 1731 retrotransposon LTR, repressing retrotransposon promoter activity in *Drosophila* (Lacoste et al., 1995).

***CDK5RAP3 (CG30291)***: CG30291 is the *Drosophila* ortholog of mammalian CDK5RAP3 that influence co-activation of STAT3-dependent transcription and modulation of p53 and ERK (Wamsley et al., 2017; Egusquiaguirre et al., 2020; Li et al., 2020). CG30291 as a broadly Myc-associated factor, may be linked to PIC-assembled, paused transcriptional states at promoters and enhancers of *Drosophila Myc*.

***HmgD and Hmg-2***: HmgD and Hmg-2 are high mobility group (HMG) proteins that modulate chromatin structure and act as downstream effectors of Wingless (Wnt/Wg) and TGF-β/Dpp signaling, in part through interactions with the corepressors Groucho and CtBP (Dragan et al., 2003; Helman et al., 2011). These factors enable context-dependent transcriptional regulation by facilitating the transmission of signaling inputs to promoter-associated transcriptional complexes.

***Isha (CG4266)***: Isha is a suppressor of hairy wing su(Hw) mRNA adaptor protein that recruits su(Hw) mRNA to chromatin insulator complexes and negatively regulates initiating and elongating RNA Pol (Bag et al., 2022). Isha was detected at (P29/P30, P31/P32, P35/P36, and P37/P38), including functional transcription initiation sites, with P29/P30 and P35/P36 acting cooperatively within the transcribing unit. Although its association with Myc-CRMs suggests a regulatory role, direct evidence linking Isha to *Myc* transcription is currently lacking.

***SAYP/Enhancer of yellow 3 [e(y)3]***: SAYP is a component of the SWI/SNF (Brahma) complex and TFIID that promotes transcription initiation by facilitating assembly of the initiation machinery (Vorobyeva et al., 2009). SAYP was detected at multiple Myc-CRMs, including transcription initiation sites, suggesting a contribution to *Myc* promoter activity.

***RNA Pol II elongation- and pause-release–associated factors***: In addition to the core elongation machinery discussed in the main text, several additional RNA Pol II elongation- and pause-release–associated factors were detected at *Myc* core promoters. The male fertility–specific protein tPlus3a was identified as a low-input, high-specificity hit at both DPE- and TATA-CorePromoters (Table S6). tPlus3a is a component of the Cdc73/PAF1 complex and has been implicated in binding phosphorylated Pol II CTD phosphoserine residues (Hundertmark et al., 2019), suggesting a role in Pol II elongation on *Myc*-like genes.

Other elongation-associated factors, including TFIIA-S, TFIIS, Rtf1, NELF-A, Spt4/Spt5 (DSIF), and EloC, were detected but not enriched, consistent with stage-specific or transient association during early development.

***mod(mdg4) and CP190***: mod(mdg4) is a multifunctional chromatin-associated protein involved in enhancer blocking, transcriptional regulation, and chromosome segregation (Gerasimova et al., 1995; Matsui et al., 2011). CP190, a BTB/POZ domain transcription factor, contributes to insulator function and heterochromatin boundary formation via interactions with CTCF, Su(Hw), and mod(mdg4) (Pai et al., 2004; Mohan et al., 2007). Both proteins were associated with multiple tested *Myc cis*-regulatory modules, although mod(mdg4) was absent from the Late-Enhancer (P1/P2) cluster. Given the presence of a gypsy element within the 5′-UTR of *Myc* (Results, Section 3.9), the CP190/CTCF/Su(Hw)/mod(mdg4) complex may contribute to developmental regulation of *Myc* transcription.

***Relative of woc (ROW)***: a putative zinc finger transcription factor, associates with HP1c and WOC to form a chromatin-bound complex at active genes, supporting Pol II-mediated transcription (Font-Burgada et al., 2008; Abel et al., 2009). ROW was detected at all tested Myc-CRMs, suggesting that the ROW/WOC/HP1c complex may influence *Myc* transcription through chromatin-based regulatory mechanisms.

***Bin1 (Bin1/Sin3A/HDAC1)***: Bicoid interacting protein 1 (Bin1/SAP18), a subunit of the Sin3/HDAC1 histone deacetylase complex, mediates transcriptional repression during *Drosophila* embryogenesis and associates with Polycomb group response elements via GAGA factor (Espinas et al., 2000; Matyash et al., 2009). Bin1 was detected at all tested Myc-CRMs, potentially contributing to modulation of *Myc* transcription via histone deacetylation.

***Caf1-55 potential regulator of Myc via chromatin assembly***: Caf1-55 (Nurf55/p55), a chromatin assembly factor, participates in DNA replication–coupled nucleosome assembly and transcriptional regulation (Tyler et al., 1996), and is linked to Polycomb-mediated H3K27me3 deposition (Anderson et al., 2011). Caf1-55 was associated with all tested Myc-CRMs, consistent with a potential role in epigenetic regulation of *Myc* transcription.

***Iswi and MRG15 may regulate Myc***: Iswi, the ATPase subunit of the NuRF chromatin remodeling complex (Corona et al., 2000), was enriched at the Late-Enhancer (P1/P2) and TATA-/DPE-CorePromoters. NuRF interacts with Myc/Max heterodimers in *Drosophila* (Xia et al., 2014), suggesting Iswi may influence *Myc* transcription via chromatin modulation. MRG15, a component of the Tip60/NuA4 histone acetyltransferase complex (Kusch et al., 2004), was highly abundant across all Myc-CRMs, potentially contributing to *Myc* regulation through histone acetylation.

***Snr1/BAP45 regulates Myc transcription***: Snf-related 1 (Snr1/BAP45) functions as a tumor suppressor during S phase, repressing *Myc* target genes such as Cyclin E (Brumby et al., 2002). Enriched at multiple Myc-CRMs, Snr1 may maintain transcription of differentiation-associated genes while limiting Myc activity via histone lysine methylation through its SET-domain interactions (Curtis et al., 2011).

***BAP111/BAF57 may regulate Myc transcription***: BAP111/BAF57, a DNA-binding component of the Brahma and Polybromo complexes, interacts with HMG-box transcription factors to modulate chromatin remodeling and activate developmental promoters (Collins and Treisman, 2000; Papoulas et al., 2001; Mohrmann et al., 2004). BAP111 was detected at all tested Myc-CRMs, suggesting a role in coordinating chromatin accessibility at *Myc* regulatory regions.

***BAP155/Moira, potentially with SPT20, may regulate Myc***: BAP155/Moira, a core SWI/SNF (Brahma) and TrxG subunit, is essential for developmental patterning and proliferation, including G1/S control via Cyclin E (Terriente-Felix and de Celis, 2009). BAP155/Moira associates with all Myc-CRMs, supporting a role in *Myc* regulation. SPT20, a SAGA subunit mediating histone acetylation and Pol II transcription, was detected at the Late-Enhancer (P1/P2), DPE-Enhancer (P29/P30), and TATA-CorePromoter (P31/P32), consistent with functions in *Myc* transcription.

***Set1/Ash2 regulates Myc transcription*****:** Ash2, a core TrxG histone methyltransferase complex subunit, catalyzes H3K4 trimethylation and cooperates with other chromatin factors to promote transcription during development (Perez-Lluch et al., 2011). Ash2 was associated with all tested Myc-CRMs, consistent with a role in establishing transcriptionally permissive chromatin states for *Myc* in response to mitogenic stimuli.

#### *S3.11. Functional context of factors at the Myc TATA-CorePromoter (P31/P32)*

***Basal transcription machinery***: RPB1, the largest RNA polymerase II subunit, is essential for transcription initiation and elongation (Zehring et al., 1988; Brickey and Greenleaf, 1995). RPB2 (Polr2B) is required for transcription at nearly all start sites (Aoyagi and Wassarman, 2000). RPB3 supports heat-induced Hsp70 expression (Yao et al., 2007); Hsp70, together with the Myc target Bag1, forms a chaperone complex that counteracts Myc-induced apoptosis (Gennaro et al., 2019). RPB7 binds ssDNA/RNA and links open chromatin to transcription termination, translation initiation, and RNA processing (GO Reference Genome Project, 2011). Collectively, these Pol II subunits support context-dependent assembly of a functional RNA Pol II complex at the *Myc* TATA-CorePromoter.

***Promoter melting, initiation, elongation, and chromatin regulation***: Cyclin H, the CAK subcomplex of TFIIH, phosphorylates the Pol II CTD during promoter escape (Larochelle et al., 2001). The TFIIH core subunit Ssl1/p44 interacts with XPD and contributes to promoter clearance (Seroz et al., 2000). e(y)3/SAYP, a shared component of SWI/SNF remodeling and TBP/TFIID initiation complexes facilitates developmental transcriptional activation (Shidlovskii et al., 2005), and was detected at multiple *Myc cis*-regulatory modules (CRMs), including the TATA promoter.

***Elongation factors***: include SPT6, a Pol II elongation factor and histone H3 chaperone that recruits SPT5/DSIF (Adelman et al., 2006), enriched at all Myc-CRMs. Topoisomerase II (Top2), associated exclusively with the Late-Enhancer (P1/P2), and Barren (Cap-H), a chromatin-binding protein, associated with both the P1/P2 cluster and the Oocyte Element (P37/P38), indicating element-preferential chromatin regulation rather than general TATA-linked activity. Chromatin-associated regulators include Bin1/SAP18, which recruits the Sin3A–HDAC1 complex and can collaborate with GAF at PRE-linked targets (Espinas et al., 2000). This is consistent with MYC recruiting Sin3A-HDAC1 at specific targets (Liu et al., 2023). Set1/Ash2 COMPASS components, together with *Myc* modulators Psi and Half pint were enriched across all tested CRMs (Guo et al., 2016).

***Transcription termination and 3′-end processing***: Cpsf160, a core AAUAAA recognition factor of the CPSF complex, showed enrichment at the distal Late-Enhancer (P1/P2), DPE-Enhancer (P29/P30), and Oocyte Element (P37/P38). It coordinates RNA 3′-end processing and polyadenylation via interactions with poly(A) polymerase (PAP) and other CPSF subunits, and associates with *MYC* transcripts during cleavage, polyadenylation, and FBW7-mediated turnover (Glover-Cutter et al., 2008; Sim et al., 2024).

**Poly(A)-binding protein (PABP) associated with all *Myc cis*-elements**: PABP stabilizes poly(A) tails, stimulates PAP activity, and links cleavage/polyadenylation to transcription termination, particularly during oogenesis and the maternal-to-zygotic transition (Clouse et al., 2008). Notably, PABP interacts with G-quadruplex–binding factors, relevant given G4-quadruplex structures in *MYC* promoters (Siddiqui-Jain et al., 2002).

*S3.12. Mechanistic details and factor associations at Myc DPE promoter and the DPE enhancer*

Elongation and termination machinery factors identified but not enriched included Wdr82, a component of Set1C/COMPASS (H3K4 methylation and Pol II negative elongation; Mohan et al., 2007; Brewer-Jensen et al., 2016), and CPSF100 (termination/3′-end processing; Brewer-Jensen et al., 2016), along with additional general elongation components (see Tables S3, S6 for full lists).

Additional associated coregulators included His2Av with ~25% nucleosome occupancy, and roles in transcription, DNA repair, and stem cell fate (Kusch et al., 2004; Tang et al., 2021); cropped/AP-4 (cell proliferation, tissue growth, muscle development; King-Jones et al., 1999; Dobi et al., 2014); Putzig (pzg) activates Notch, Ecdysone, and JAK/STAT in proliferating cells. Putzig–TRF2–DREF complex drives replication genes and may regulate *Myc* expression during G1/S phase of cell cycle. (Kugler et al., 2011; Kugler and Nagel, 2007).

*S3.13. Mechanistic description of regulatory factors associated with the Myc Oocyte Element (P37/P38)*

***Components of PIC assembly***: RPB7 (Polr2G) associates with RPB4 (Polr2D) to form the dissociable two-subunit protein RPB4/7.

***Chromatin remodelers and other regulators: Simjang (simj/p66)***: A subunit of Mi-2/NuRD complex, involved in transcriptional repression via chromatin remodeling and histone deacetylation. It recognizes methylated CpG islands and contributes to embryonic heart development via epigenetic control (Kim et al., 2004).
***BAF (baf)***: Drives chromosome condensation and karyosome formation in the oocyte nucleus, potentially modulating *Myc* locus accessibility through nuclear reorganization (Lancaster et al., 2007).
***Trol/Perlecan***: Terribly reduced optic lobes (trol) encodes the ECM protein Perlecan, a key modulator of multiple developmental signaling pathways—including Hedgehog, Wingless, EGFR, FGFR, and VEGFR—likely influencing fate specification and tissue patterning during development (Park et al., 2003; Dragojlovic-Munther and Martinez-Agosto, 2013).
***Spaghetti Squash (sqh)***: encodes the Myosin II light chain and is essential for key developmental morphogenetic processes, including mitosis, gastrulation, syncytial cellularization, and oogenesis, potentially influencing *Myc* during cell growth, proliferation, and differentiation (Somma et al., 2002; Royou et al., 2004).
***Cdt2 (l(2)dtl)***: A substrate adapter for the Cul4-DDB1 E3 ubiquitin ligase, controlling replication- and DNA damage-coupled proteolysis (Arias and Walter, 2006). Cdt2 regulates Myc stability (Dammai et al., 2003; Sloan et al., 2012) and was exclusively detected at the Oocyte Element (P37/P38) (Table 6), extending the Cdt2–Myc regulatory axis to oogenesis.
***Awd (awd)***: A nucleoside-diphosphate kinase (homolog of human Nm23/NME1/2), involved in vesicle trafficking, receptor internalization, and modulation of signaling pathways (e.g., Notch and EGFR, known *Myc* regulators). Its human homologs NmE1/2 are essential for tissue patterning (Ignesti et al., 2014). Its enrichment at all tested Myc-CRMs (including the Oocyte Element) may highlight a role in *Myc* regulation during early oocyte development.
***Smn (Smn)***: Survival motor neuron (Smn) is a subunit of SMN complex that has been implicated in c-MYC–dependent apoptosis and is required for oocyte maturation (Vyas et al., 2002; Borg et al., 2015). SMN was detected across multiple *Myc cis*-regulatory regions, indicating that Smn broadly associates with regulatory elements controlling *Myc* during early development.

**Post-transcriptional and post-translational factors at the Oocyte Element:**

***CPSF5 and CPSF160*** are components of the cleavage and polyadenylation specificity factor complex involved in mRNA 3′-end processing. In the context of *c-MYC*, they contribute to 3′-end processing and may also influence mRNA turnover via FBW7-dependent, proteasome-mediated degradation. In *Drosophila*, they may additionally contribute to *Myc* transcription termination and mRNA processing (Glover-Cutter et al., 2008; Sim et al., 2024).

***UNR*** is an RNA-binding protein regulating cap-independent translation of *c-MYC* mRNAs (Mitchell et al., 2001). It is developmentally regulated and may also modulate *Myc* mRNA stability and translation in a context-dependent manner. In *Drosophila*, UNR regulates *Myc* translation and contributes to dosage compensation and broader mRNA regulation (Militti et al., 2014).

***SBDS*** is required for 60S ribosomal subunit maturation and global translation. Its disruption alters *MYC* expression, likely via translational control. In this study, SBDS was enriched at the DPE Core-Promoter and Oocyte Element, suggesting coupling between transcriptional initiation sites and translational capacity during growth-associated programs (In et al., 2016).

***eIF3j*** is a component of the eIF3 complex essential for translation initiation in proliferating cells. Its selective enrichment at the Oocyte Element suggests a role in modulating *Myc* translation during oogenesis, linking local translational control to developmental regulation (Andersen and Leevers, 2007).

***Nucleostemin 3 (Ns3)*** is a conserved GTPase required for 60S ribosomal export and growth control (Hartl et al., 2013). It was strongly enriched at the Late-Enhancer (P1/P2), DPE-Core Promoter (P35/P36), and Oocyte Element (P37/P38), all active regulatory regions during oogenesis and early embryogenesis. This enrichment suggests a functional link between ribosome biogenesis, translational capacity, and Myc-driven growth programs (Zacarias-Fluck et al., 2024).

***Effete (eff, UbcD1)*** is an E2 ubiquitin-conjugating enzyme involved in protein turnover and germline stem cell maintenance. Its role in ubiquitin-mediated protein regulation suggests potential involvement in controlling Myc protein stability during developmental cell cycles (Herman-Bachinsky et al., 2007; Chen et al., 2009).

*S3.14. Extended details of posttranscriptional/-translational regulatory factors identified at Myc-CRMs*

Detailed quality control metrics, contaminant exclusion analyses, replicate peak-overlap statistics, abundance ratios, and statistical factor classification values are provided in Supplementary Data S2–S7 and S9–S11. The summaries below focus on regulator-specific contextualization and biological interpretation. See also Fig. S6 and Results Fig.14.

**Details on specific regulators:**

***Ogt/sxc***: Beyond MYC stabilization (Itkonen et al., 2013; Itkonen et al., 2019), Ogt/Sxc has been implicated in coupling metabolic state to chromatin regulation through dynamic O-GlcNAc cycling at Polycomb-associated loci. In *Drosophila*, Sxc functions within Polycomb repressive complexes and modulates histone-associated regulatory networks, thereby playing a key role in regulating epigenetic gene silencing and during developmental transitions (Gambetta et al., 2009; Akan et al., 2016). In our analysis, Ogt/Sxc occupancy was detected across multiple CRM subclasses, including both promoter-proximal and distal elements. Enrichment strength was consistent across developmental stages, suggesting stage-independent recruitment.

***Nmt (Nmt)***: In *Drosophila*, N-myristoyl transferase (Nmt) was detected in the nuclear extract and exclusively enriched at the Late-Enhancer (P1/P2) (100% abundance ratio), a differentiation-associated regulatory element, suggesting that Nmt recruitment is both CRM-specific and stage-associated, consistent with a potential role in modulating *Myc* expression during developmental transitions.

***Pin1/dod***: Dodo was associated with the Late-Enhancer (P1/P2) as well as TATA-Core (P31/P32) and DPE-Core Promoters (P35/P36), suggesting a potential role in regulating both TATA- and DPE-driven transcription during differentiation. In *Drosophila*, Dodo facilitates rhomboid expression by promoting degradation of Chorion factor 2 (Cf2), contributing to dorsoventral patterning of the follicular epithelium (Hsu et al., 2001). While the main text summarizes its role in regulating MYC stability and activity, these additional functional links highlight the broader developmental and regulatory context in which Dodo operates.

***Spaghetti/Spag***: Spag is a cochaperone and associates with HSP70 and HSP90 to mediate protein-protein interactions, facilitating the assembly of multiprotein complexes (Das et al., 1998). Within RNA Pol II complexes, Spag contributes to transcription machinery stability and transcriptional regulation (Rodriguez and Llorca, 2020). In ribonucleoprotein complexes, it mediates protein folding immediately upon translation completion (Boulon et al., 2008). Spag has additionally been implicated in regulation of circadian rhythm and apoptosis in optic lobes (Means et al., 2015). Here, our LC MS/MS analysis identified Spag as highly abundant at all tested Myc-CRMs, which suggests a potential role in regulating *Myc* expression, although this remains unresolved.

**CG7546 (*clone 2.45*)**: CG7546 is a large proline-rich protein and a member of the BAT3 complex, involved in ubiquitin-dependent degradation, DNA damage response, and apoptosis (Kobayashi et al., 2019). LC-MS/MS data analysis identified CG7546 at the DPE-Enhancer (P29/P30), TATA-Core Promoters (P31/P32), and Oocyte Element (P37/P38), highlighting CRM-specific recruitment patterns that may support future studies of *Myc* regulation.

***Rumi***: Is an EGF-domain serine O-glucosyl-/O-xylosyltransferase and O-glucosylates Notch in the ER (Acar et al., 2008; Takeuchi et al., 2011). Its exclusive enrichment at the DPE-CorePromoter (P35/P36), together with its link to Notch signaling—a known regulator of *Myc*—suggests a potential role for Rumi in regulating *Myc* expression during development.

***Poe***: Poe (Purity of essence, calmodulin-binding E3 ubiquitin ligase) is involved in protein metabolism and developmental processes (Xu et al., 1998; Ashton-Beaucage et al., 2016), and was exclusively enriched at the DPE-Core Promoter (P35/P36), highlighting a potential role for Poe in regulating Myc protein turnover via DPE transcription, and providing a framework for further investigation of protein stability and differentiation-associated regulatory networks.

**Details on processing and translation factors:**

Several RNA-binding, RNA processing, helicases, and translation factors were enriched at the Myc-CRMs, indicating roles in regulation of *Myc* mRNA surveillance and protein stability during *Drosophila* development (Fig. S6; Table S3).

***Larp4B***: La-related protein 4B (Larp4B) is an RNA-binding protein that mediates post-transcriptional inhibition of *Myc* mRNA translation, contributing to the downregulation of cell and organ growth (Funakoshi et al., 2018). Larp4B was associated with all the tested Myc-CRMs, suggesting potential mechanistic links between *Myc* translational regulation and developmental growth control.

Pumilio/Pop2: Pumilio (pum), RNA-binding protein and mRNA decay complex, modulates RNA stability, translation repression, and mRNA degradation (Dean et al., 2002; Kim et al., 2012). Target mRNAs deadenylated by Pop2 are recognized by repression domains of Pumilio for destruction during stem cell differentiation, neurogenesis, and embryogenesis (Arvola et al., 2020). Pumilio regulates Hunchback mRNA in early embryos, germline stem cell proliferation and migration (Asaoka-Taguchi et al., 1999; Wickens et al., 2002), and fate specification in sensory organ precursor (SOP) cells via EGFR signaling (Kim et al., 2012). In our dataset, Pumilio/Pop2 was detected at all tested Myc-CRMs, with the exception of Pop2, which was absent from the Late-Enhancer (P1/P2). This broad recruitment across CRM types indicates that the complex may contribute to fine-tuning *Myc* mRNA levels during developmental processes such as bristle and wing vein formation.

***Ddx1***: Dead-box RNA helicase 1 is involved in RNA trafficking, nucleic acid duplex unwinding (Gene Ontology Curators, 2002), translation, and protein synthesis (Rafti et al., 1996). In the early embryo, Ddx1 positively regulates cell proliferation and contributes to body size and germ cell development (Rafti et al., 1996; Germain et al., 2015). The human homolog of Ddx1 is amplified in MYCN-amplified tumors such as retinoblastoma (Godbout and Squire, 1993; Manohar et al., 1995; Godbout et al., 1998). Furthermore, in pediatric neuroblastoma, the MYC-binding partner Max shows strong interaction with Ddx1 (Jin et al., 2021). Ddx1 was associated with multiple Myc-CRMs, suggesting that Ddx1 may be indirectly involved in regulating Myc activity through Max.

***Abstrakt***: Abstrakt is a splicing factor that contributes to pre-mRNA processing and has been shown as direct target of MYC/MAX network with roles in proliferation, germline development, and determination of body size (Grandori et al., 1996; Herold et al., 2009). In LC-MS/MS dataset, Abstrakt was detected across Myc-CRMs; this broad occupancy suggests a potential role for Abstrakt in regulating *Myc* expression during development, potentially through RNA processing mechanisms, but functional validation is required.

**How/Struthio**: How/Struthio ('held out wings') is a conserved KH domain RNA-binding protein that regulates alternative splicing, with How(L) destabilizing RNA and How(S) stabilizing RNA during embryonic differentiation and morphogenesis (Walsh and Brown, 1998; Volohonsky et al., 2007). It integrates multiple developmental signals—including EGFR, MAPK/ERK, DPP/BMP, Hedgehog, Wnt-TCF, and Akt/Ras pathways—to control cell cycle progression and differentiation (Riese et al., 1997; Martin-Bermudo, 2000; Marsden & DeSimone, 2001; Edenfeld et al., 2006; Israeli et al., 2007; Toledano-Katchalski et al., 2007; Lobbardi et al., 2011; Nir et al., 2012). The mouse homolog Quaking is a tumor suppressor regulating similar processes (Biedermann et al., 2010). How/Struthio was detected at multiple Myc-CRMs, suggesting broad recruitment and a potential role in modulating *Myc* expression during cell cycle progression and fate specification. wg expression (Park et al., 2003; Schneider et al., 2006; Lindner et al., 2007)

***Smaug (smg)***: is the founding member of the SMAUG family of sequence-specific RNA-binding proteins. It functions as a translation repressor during the maternal-to-zygotic transition in early embryos (Dean et al., 2002; Kim and Bowie, 2003) and promotes the degradation of hundreds of maternal mRNAs by recruiting the CCR4-NOT deadenylase complex (Jeske et al., 2006; Zaessinger et al., 2006; Chartier et al., 2015). Through this activity, Smaug regulates embryonic patterning (Smibert et al., 1999), the activation of zygotic transcription, and early embryonic cell cycles (Liang et al., 2008). These functions, together with its specific identification at the Oocyte Element (P37/P38), suggest a potential role for Smaug in regulating *Myc* during early development.

***Obelus (obe)***: Obelus is a member of Ski2 family of DNA helicases (Vichas et al., 2015). It is required for adherence junction arrangement and for the alternative splicing of Crumbs (*crb*) mRNA, which encodes a transmembrane protein essential for apico-basal cell polarity during embryogenesis and oogenesis (Vichas et al., 2015). Myc, a master regulator of growth and proliferation, must be tightly controlled to maintain the balance between cell polarity establishment and proliferation rates, thereby preventing malignant transformation (Grifoni et al., 2013). Here, we identified Obelus, a regulator of apico–basal polarity, in association with the DPE-CorePromoter (P35/P36). These findings suggest that Obelus may act as a potential regulator of *Myc* mRNA and protein levels during early development.

***Saf-B***: Scaffold attachment factor B (Saf-B) is an RNA- and DNA-binding protein involved in regulating alternative mRNA splicing (Park et al., 2004) and pol II transcription (GO Reference Genome Project, 2011). In *Drosophila* SAF-B connects the nuclear matrix with chromatin and transcriptional complexes, contributing to higher-order nuclear organization (Alfonso-Parra and Maggert, 2010). The human *MYC* gene contains a nuclear matrix attachment region (MAR) within its 3′-UTR that binds regulatory proteins such as Sox2 (Lei et al., 2005). MARs function as boundary elements safeguard genomic regions during replication and transcription (Bode et al., 1996; BODE et al. 2003). Given its architectural role and MAR association, Saf-B may support nuclear organization during *Myc* transcriptional regulation.

***Exon Junction Complex (EJC)***: Core components eIF4A3 (CG7483, RNA helicase scaffold), Ddx1 (RNA processing factor), and Mago/Y14 (mRNP export adaptor), together with Barentsz (btz) (EJC assembly and stability factor) (Palacios et al., 2004), were enriched at all tested Myc-CRMs (Pym detected at TATA-CorePromoter; Mago showed signal but not enriched). Given the EJC's role in co-transcriptional mRNP assembly and downstream mRNA export and stability, their presence at *Myc* regulatory regions suggests coupling of *Myc* transcription with post-splicing mRNA processing steps.

**Additional specialized factors:**

***Bic/Btf (Bicaudal)***: bic/bicaudal encodes NACβ, a ribosome-associated factor supporting translational activity (Kogan et al., 2017). NAC has also been implicated in transcriptional regulation during development (Rospert et al., 2002). Proteomic analyses revealed the coexpression of BTF with MYCN and MYC targets including NDRG1 and ODC1 (Hogarty et al., 2008; Symes et al., 2013), supporting integration within *MYC* regulatory networks. Broad

***CNBP***: Conserved small CCHC-type zinc finger protein (CNBP) binds ssDNA/RNA and stabilizes G-rich mRNAs by preventing G-quadruplex formation (InterPro Project Members, 2004; GO Reference Genome Project, 2011). In *Drosophila*, CNBP promotes Myc protein synthesis via IRES-dependent translation and is essential for viability, as its loss results in early embryonic lethality; moreover, CNBP regulates wing disc morphogenesis and size by modulating Myc levels (Antonucci et al., 2014). In this study, CNBP was detected at the Late-Enhancer (P1/P2), DPE-CorePromoter (P35/P36), and Oocyte Element (P37/P38), and reporter analyses confirmed their functional contribution to *Myc* expression (Figs. 1–2), demonstrating that CNBP directly engages multiple *cis*-regulatory elements that shape *Myc* expression during *Drosophila* development.

***Prp31***: Pre-mRNA processing factor 31 (Prp31) is a splicing factor required for pre-mRNA processing and photoreceptor development (Mount and Salz, 2000; Herold et al., 2009; Ray et al., 2010). Loss or reduction of Prp31 leads to embryonic lethality in mice (Bujakowska et al., 2009). Given MYC's role in regulating splicing machinery during development and disease (Koh et al., 2015; Abou Faycal et al., 2016; Jablonowski et al., 2023; Phillips et al., 2020), Prp31 may contribute to *Drosophila Myc* regulation.

***Spenito (Nito)***: Spenito is an RNA-binding Wnt-TCF effector required for alternative mRNA splicing during compound eye development (Abou Faycal et al., 2016; Jablonowski et al., 2023). As a component associated with the m$^6$A writer-reader machinery (WMM complex), Spenito promotes m$^6$A methylation to facilitate splicing of targets such as Sex-lethal (Sxl) during sex determination and dosage compensation (Yan and Perrimon, 2015). LC-MS/MS specifically identified Spenito at the Late-Enhancer (P1/P2) cluster, a regulatory element required for *Myc* patterning in larval imaginal discs and brain (Fig. 1A, 1F). Given that Myc, Sxl, and transformer (tra) collectively regulate sex-specific body size dimorphism (Mathews et al., 2017), we hypothesize that Spenito modulates *Myc* mRNA splicing during larval development, potentially linking Wnt signaling, $m^6A$ modification, and growth control.

***Su (f)***: Suppressor of forked (Su[f]) is an essential component of the CSF cleavage complex involved in mRNA 3′-end formation and alternative poly(A) site selection, particularly in proliferating cells (Audibert and Simonelig, 1999; InterPro Project Members, 2004; GO Reference Genome Project, 2011). Su(f) was enriched at the Late-Enhancer (P1/P2) and Oocyte Element (P37/P38) enhancer clusters, suggesting a role in *Myc* transcription termination and mRNA processing. High Su(f) levels occur during mitosis, and the protein also negatively regulates the gypsy retrotransposon, which is inserted in the proximal 5′ region of the *Myc* gene near P37/P38 (Parkhurst and Corces, 1986; Mitchelson et al., 1993; Gallant et al., 1996).

Ribosomal and translational machinery components, encompassing multiple cytoplasmic and mitochondrial 40S/60S subunits as well as initiation, elongation, and termination factors, were also associated with Myc-CRMs, pointing to a potential translational feedback loop coordinating *Myc* expression during tissue growth.

Interpretation of all these factors in the context of a multi-layered and complex regulation of *Myc* expression is discussed in the main text.

### Supplementary References


Abel, J., Eskeland, R., Raffa, G.D., Kremmer, E., Imhof, A., 2009. Drosophila HP1c is regulated by an auto-regulatory feedback loop through its binding partner Woc. PLoS One 4, e5089.

Akan, I., Love, D.C., Harwood, K.R., Bond, M.R., Hanover, J.A., 2016. Drosophila O-GlcNAcase Deletion Globally Perturbs Chromatin O-GlcNAcylation. J Biol Chem 291, 9906-9919.

Alfonso-Parra, C., Maggert, K.A., 2010. Drosophila SAF-B links the nuclear matrix, chromosomes, and transcriptional activity. PLoS One 5, e10248.

Anderson, A.E., Karandikar, U.C., Pepple, K.L., Chen, Z., Bergmann, A., Mardon, G., 2011. The enhancer of trithorax and polycomb gene Caf1/p55 is essential for cell survival and patterning in Drosophila development. Development 138, 1957-1966.

Arias, E.E., Walter, J.C., 2006. PCNA functions as a molecular platform to trigger Cdt1 destruction and prevent re-replication. Nat Cell Biol 8, 84-90.

Asaoka-Taguchi, M., Yamada, M., Nakamura, A., Hanyu, K., Kobayashi, S., 1999. Maternal Pumilio acts together with Nanos in germline development in Drosophila embryos. Nat Cell Biol 1, 431-437.

Badenhorst, P., Voas, M., Rebay, I., Wu, C., 2002. Biological functions of the ISWI chromatin remodeling complex NURF. Genes Dev 16, 3186-3198.

Bag, I., Chen, Y., D'Orazio, K., Lopez, P., Wenzel, S., Takagi, Y., Lei, E.P., 2022. Isha is a su(Hw) mRNA-binding protein required for gypsy insulator function. G3 (Bethesda) 12.

Bajpai, R., Makhijani, K., Rao, P.R., Shashidhara, L.S., 2004. Drosophila Twins regulates Armadillo levels in response to Wg/Wnt signal. Development 131, 1007-1016.

Barral, A., Dejardin, J., 2023. The chromatin signatures of enhancers and their dynamic regulation. Nucleus 14, 2160551.

Biedermann, B., Hotz, H.R., Ciosk, R., 2010. The Quaking family of RNA-binding proteins: coordinators of the cell cycle and differentiation. Cell Cycle 9, 1929-1933.

Bode, J., Stengert-Iber, M., Kay, V., Schlake, T., Dietz-Pfeilstetter, A., 1996. Scaffold/matrix-attached regions: topological switches with multiple regulatory functions. Crit Rev Eukaryot Gene Expr 6, 115-138.

Brewer-Jensen, P., Wilson, C.B., Abernethy, J., Mollison, L., Card, S., Searles, L.L., 2016. Suppressor of sable [Su(s)] and Wdr82 down-regulate RNA from heat-shock-inducible repetitive elements by a mechanism that involves transcription termination. RNA 22, 139-154.

Brickey, W.J., Greenleaf, A.L., 1995. Functional studies of the carboxy-terminal repeat domain of Drosophila RNA polymerase II in vivo. Genetics 140, 599-613.

Brumby, A.M., Zraly, C.B., Horsfield, J.A., Secombe, J., Saint, R., Dingwall, A.K., Richardson, H., 2002. Drosophila cyclin E interacts with components of the Brahma complex. EMBO J 21, 3377-3389.

Bujakowska, K., Maubaret, C., Chakarova, C.F., Tanimoto, N., Beck, S.C., Fahl, E., Humphries, M.M., Kenna, P.F., Makarov, E., Makarova, O., Paquet-Durand, F., Ekstrom, P.A., van Veen, T., Leveillard, T., Humphries, P., Seeliger, M.W., Bhattacharya, S.S., 2009. Study of gene-targeted mouse models of splicing factor gene Prpf31 implicated in human autosomal dominant retinitis pigmentosa (RP). Invest Ophthalmol Vis Sci 50, 5927-5933.

Cadigan, K.M., 2012. TCFs and Wnt/beta-catenin signaling: more than one way to throw the switch. Curr Top Dev Biol 98, 1-34.

Cappadocia, L., Lima, C.D., 2018. Ubiquitin-like Protein Conjugation: Structures, Chemistry, and Mechanism. Chem Rev 118, 889-918.

Chang, Y.C., Wu, J.W., Hsieh, Y.C., Huang, T.H., Liao, Z.M., Huang, Y.S., Mondo, J.A., Montell, D., Jang, A.C., 2018. Rap1 Negatively Regulates the Hippo Pathway to Polarize Directional Protrusions in Collective Cell Migration. Cell Rep 22, 2160-2175.

Chartier, A., Klein, P., Pierson, S., Barbezier, N., Gidaro, T., Casas, F., Carberry, S., Dowling, P., Maynadier, L., Bellec, M., Oloko, M., Jardel, C., Moritz, B., Dickson, G., Mouly, V., Ohlendieck, K., Butler-Browne, G., Trollet, C., Simonelig, M., 2015. Mitochondrial dysfunction reveals the role of mRNA poly(A) tail regulation in oculopharyngeal muscular dystrophy pathogenesis. PLoS Genet 11, e1005092.

Cho, I.K., Chang, C.L., Li, Q.X., 2013. Diet-induced over-expression of flightless-I protein and its relation to flightlessness in Mediterranean fruit fly, Ceratitis capitata. PLoS One 8, e81099.

Collins, R.T., Treisman, J.E., 2000. Osa-containing Brahma chromatin remodeling complexes are required for the repression of wingless target genes. Genes Dev 14, 3140-3152.

Corona, D.F., Eberharter, A., Budde, A., Deuring, R., Ferrari, S., Varga-Weisz, P., Wilm, M., Tamkun, J., Becker, P.B., 2000. Two histone fold proteins, CHRAC-14 and CHRAC-16, are developmentally regulated subunits of chromatin accessibility complex (CHRAC). EMBO J 19, 3049-3059.

Curtis, B.J., Zraly, C.B., Marenda, D.R., Dingwall, A.K., 2011. Histone lysine demethylases function as co-repressors of SWI/SNF remodeling activities during Drosophila wing development. Dev Biol 350, 534-547.

Dean, K.A., Aggarwal, A.K., Wharton, R.P., 2002. Translational repressors in Drosophila. Trends Genet 18, 572-577.
Dey, B., Thukral, S., Krishnan, S., Chakrobarty, M., Gupta, S., Manghani, C., Rani, V., 2012. DNA-protein interactions: methods for detection and analysis. Mol Cell Biochem 365, 279-299.
Dragan, A.I., Klass, J., Read, C., Churchill, M.E., Crane-Robinson, C., Privalov, P.L., 2003. DNA binding of a non-sequence-specific HMG-D protein is entropy driven with a substantial non-electrostatic contribution. J Mol Biol 331, 795-813.
Egusquiaguirre, S.P., Liu, S., Tosic, I., Jiang, K., Walker, S.R., Nicolais, M., Saw, T.Y., Xiang, M., Bartel, K., Nelson, E.A., Frank, D.A., 2020. CDK5RAP3 is a co-factor for the oncogenic transcription factor STAT3. Neoplasia 22, 47-59.
Espinas, M.L., Canudas, S., Fanti, L., Pimpinelli, S., Casanova, J., Azorin, F., 2000. The GAGA factor of Drosophila interacts with SAP18, a Sin3-associated polypeptide. EMBO Rep 1, 253-259.
Font-Burgada, J., Rossell, D., Auer, H., Azorin, F., 2008. Drosophila HP1c isoform interacts with the zinc-finger proteins WOC and Relative-of-WOC to regulate gene expression. Genes Dev 22, 3007-3023.
Gene Ontology Curators, - 2002. Manual transfer of experimentally-verified manual GO annotation data to orthologs by curator judgment of sequence similarity.
Gennaro, V.J., Wedegaertner, H., McMahon, S.B., 2019. Interaction between the BAG1S isoform and HSP70 mediates the stability of anti-apoptotic proteins and the survival of osteosarcoma cells expressing oncogenic MYC. BMC Cancer 19, 258.
Gerasimova, T.I., Gdula, D.A., Gerasimov, D.V., Simonova, O., Corces, V.G., 1995. A Drosophila protein that imparts directionality on a chromatin insulator is an enhancer of position-effect variegation. Cell 82, 587-597.
GO Reference Genome Project, - 2011. Phylogenetic annotation using the Gene Ontology.
Godbout, R., Packer, M., Bie, W., 1998. Overexpression of a DEAD box protein (DDX1) in neuroblastoma and retinoblastoma cell lines. J Biol Chem 273, 21161-21168.
Godbout, R., Squire, J., 1993. Amplification of a DEAD box protein gene in retinoblastoma cell lines. Proc Natl Acad Sci U S A 90, 7578-7582.
Goedhart, J., Luijsterburg, M.S., 2020. VolcaNoseR is a web app for creating, exploring, labeling and sharing volcano plots. Sci Rep 10, 20560.
Grandori, C., Mac, J., Siebelt, F., Ayer, D.E., Eisenman, R.N., 1996. Myc-Max heterodimers activate a DEAD box gene and interact with multiple E box-related sites in vivo. EMBO J 15, 4344-4357.
Grifoni, D., Froldi, F., Pession, A., 2013. Connecting epithelial polarity, proliferation and cancer in Drosophila: the many faces of lgl loss of function. Int J Dev Biol 57, 677-687.
Hamilton, B.J., Mortin, M.A., Greenleaf, A.L., 1993. Reverse genetics of Drosophila RNA polymerase II: identification and characterization of RpII140, the genomic locus for the second-largest subunit. Genetics 134, 517-529.
Harris, L.R., Churchward, M.A., Butt, R.H., Coorssen, J.R., 2007. Assessing detection methods for gel-based proteomic analyses. J Proteome Res 6, 1418-1425.
Helman, A., Cinnamon, E., Mezuman, S., Hayouka, Z., Von Ohlen, T., Orian, A., Jimenez, G., Paroush, Z., 2011. Phosphorylation of Groucho mediates RTK feedback inhibition and prolonged pathway target gene expression. Curr Biol 21, 1102-1110.
Hogarty, M.D., Norris, M.D., Davis, K., Liu, X., Evageliou, N.F., Hayes, C.S., Pawel, B., Guo, R., Zhao, H., Sekyere, E., Keating, J., Thomas, W., Cheng, N.C., Murray, J., Smith, J., Sutton, R., Venn, N., London, W.B., Buxton, A., Gilmour, S.K., Marshall, G.M., Haber, M., 2008. ODC1 is a critical determinant of MYCN oncogenesis and a therapeutic target in neuroblastoma. Cancer Res 68, 9735-9745.
Hsu, T., McRackan, D., Vincent, T.S., Gert de Couet, H., 2001. Drosophila Pin1 prolyl isomerase Dodo is a MAP kinase signal responder during oogenesis. Nat Cell Biol 3, 538-543.
Hundertmark, T., Kreutz, S., Merle, N., Nist, A., Lamp, B., Stiewe, T., Brehm, A., Renkawitz-Pohl, R., Rathke, C., 2019. Drosophila melanogaster tPlus3a and tPlus3b ensure full male fertility by regulating transcription of Y-chromosomal, seminal fluid, and heat shock genes. PLoS One 14, e0213177.
Ignesti, M., Barraco, M., Nallamothu, G., Woolworth, J.A., Duchi, S., Gargiulo, G., Cavaliere, V., Hsu, T., 2014. Notch signaling during development requires the function of awd, the Drosophila homolog of human metastasis suppressor gene Nm23. BMC Biol 12, 12.
InterPro Project Members, - 2004. Gene Ontology annotation through association of InterPro records with GO terms.
Israeli, D., Nir, R., Volk, T., 2007. Dissection of the target specificity of the RNA-binding protein HOW reveals dpp mRNA as a novel HOW target. Development 134, 2107-2114.
Itkonen, H.M., Urbanucci, A., Martin, S.E., Khan, A., Mathelier, A., Thiede, B., Walker, S., Mills, I.G., 2019. High OGT activity is essential for MYC-driven proliferation of prostate cancer cells. Theranostics 9, 2183-2197.
Jeske, M., Meyer, S., Temme, C., Freudenreich, D., Wahle, E., 2006. Rapid ATP-dependent deadenylation of nanos mRNA in a cell-free system from Drosophila embryos. J Biol Chem 281, 25124-25133.

Jia, H., Liu, Y., Xia, R., Tong, C., Yue, T., Jiang, J., Jia, J., 2010. Casein kinase 2 promotes Hedgehog signaling by regulating both smoothened and Cubitus interruptus. J Biol Chem 285, 37218-37226.
Jin, Y., Shi, J., Wang, H., Lu, J., Chen, C., Yu, Y., Wang, Y., Yang, Y., Ren, D., Zeng, Q., Ni, X., Guo, Y., 2021. MYC-associated protein X binding with the variant rs72780850 in RNA helicase DEAD box 1 for susceptibility to neuroblastoma. Sci China Life Sci 64, 991-999.
Jox, T., Buxa, M.K., Bohla, D., Ullah, I., Macinkovic, I., Brehm, A., Bartkuhn, M., Renkawitz, R., 2017. Drosophila CP190- and dCTCF-mediated enhancer blocking is augmented by SUMOylation. Epigenetics Chromatin 10, 32.
Kim, C.A., Bowie, J.U., 2003. SAM domains: uniform structure, diversity of function. Trends Biochem Sci 28, 625-628.
Kim, S.Y., Kim, J.Y., Malik, S., Son, W., Kwon, K.S., Kim, C., 2012. Negative regulation of EGFR/MAPK pathway by Pumilio in Drosophila melanogaster. PLoS One 7, e34016.
Kogan, G.L., Akulenko, N.V., Abramov, Y.A., Sokolova, O.A., Fefelova, E.A., Gvozdev, V.A., 2017. [Nascent Polypeptide-Associated Complex as Tissue-Specific Cofactor during Germinal Cell Differentiation in Drosophila Testes]. Mol Biol (Mosk) 51, 677-682.
Koh, C.M., Bezzi, M., Low, D.H., Ang, W.X., Teo, S.X., Gay, F.P., Al-Haddawi, M., Tan, S.Y., Osato, M., Sabo, A., Amati, B., Wee, K.B., Guccione, E., 2015. MYC regulates the core pre-mRNA splicing machinery as an essential step in lymphomagenesis. Nature 523, 96-100.
Konig, A., Shcherbata, H.R., 2015. Soma influences GSC progeny differentiation via the cell adhesion-mediated steroid-let-7-Wingless signaling cascade that regulates chromatin dynamics. Biol Open 4, 285-300.
Lacoste, J., Codani-Simonart, S., Best-Belpomme, M., Peronnet, F., 1995. Characterization and cloning of p11, a transrepressor of Drosophila melanogaster retrotransposon 1731. Nucleic Acids Res 23, 5073-5079.
Lei, J.X., Liu, Q.Y., Sodja, C., LeBlanc, J., Ribecco-Lutkiewicz, M., Smith, B., Charlebois, C., Walker, P.R., Sikorska, M., 2005. S/MAR-binding properties of Sox2 and its involvement in apoptosis of human NT2 neural precursors. Cell Death Differ 12, 1368-1377.
Lewellyn, L., Cetera, M., Horne-Badovinac, S., 2013. Misshapen decreases integrin levels to promote epithelial motility and planar polarity in Drosophila. J Cell Biol 200, 721-729.
Li, L., Williams, P., Gao, Z., Wang, Y., 2020. VEZF1-guanine quadruplex DNA interaction regulates alternative polyadenylation and detyrosinase activity of VASH1. Nucleic Acids Res 48, 11994-12003.
Liang, H.L., Nien, C.Y., Liu, H.Y., Metzstein, M.M., Kirov, N., Rushlow, C., 2008. The zinc-finger protein Zelda is a key activator of the early zygotic genome in Drosophila. Nature 456, 400-403.
Lindhorst, D., Halfon, M.S., 2023. Reporter gene assays and chromatin-level assays define substantially non-overlapping sets of enhancer sequences. BMC Genomics 24, 17.
Lindner, J.R., Hillman, P.R., Barrett, A.L., Jackson, M.C., Perry, T.L., Park, Y., Datta, S., 2007. The Drosophila Perlecan gene trol regulates multiple signaling pathways in different developmental contexts. BMC Dev Biol 7, 121.
Liu, S., Baeg, G.H., Yang, Y., Goh, F.G., Bao, H., Wagner, E.J., Yang, X., Cai, Y., 2023. The Integrator complex desensitizes cellular response to TGF-beta/BMP signaling. Cell Rep 42, 112007.
Lobbardi, R., Lambert, G., Zhao, J., Geisler, R., Kim, H.R., Rosa, F.M., 2011. Fine-tuning of Hh signaling by the RNA-binding protein Quaking to control muscle development. Development 138, 1783-1794.
Luo, S., Wehr, N.B., Levine, R.L., 2006. Quantitation of protein on gels and blots by infrared fluorescence of Coomassie blue and Fast Green. Anal Biochem 350, 233-238.
Lv, X., Pan, C., Zhang, Z., Xia, Y., Chen, H., Zhang, S., Guo, T., Han, H., Song, H., Zhang, L., Zhao, Y., 2016. SUMO regulates somatic cyst stem cell maintenance and directly targets the Hedgehog pathway in adult Drosophila testis. Development 143, 1655-1662.
Manohar, C.F., Salwen, H.R., Brodeur, G.M., Cohn, S.L., 1995. Co-amplification and concomitant high levels of expression of a DEAD box gene with MYCN in human neuroblastoma. Genes Chromosomes Cancer 14, 196-203.
Martin-Bermudo, M.D., 2000. Integrins modulate the Egfr signaling pathway to regulate tendon cell differentiation in the Drosophila embryo. Development 127, 2607-2615.
Matharu, N.K., Hussain, T., Sankaranarayanan, R., Mishra, R.K., 2010. Vertebrate homologue of Drosophila GAGA factor. J Mol Biol 400, 434-447.
Mathews, K.W., Cavegn, M., Zwicky, M., 2017. Sexual Dimorphism of Body Size Is Controlled by Dosage of the X-Chromosomal Gene Myc and by the Sex-Determining Gene tra in Drosophila. Genetics 205, 1215-1228.
Matsui, M., Sharma, K.C., Cooke, C., Wakimoto, B.T., Rasool, M., Hayworth, M., Hylton, C.A., Tomkiel, J.E., 2011. Nuclear structure and chromosome segregation in Drosophila male meiosis depend on the ubiquitin ligase dTopors. Genetics 189, 779-793.
Means, J.C., Venkatesan, A., Gerdes, B., Fan, J.Y., Bjes, E.S., Price, J.L., 2015. Drosophila spaghetti and doubletime link the circadian clock and light to caspases, apoptosis and tauopathy. PLoS Genet 11, e1005171.

Mellacheruvu, D., Wright, Z., Couzens, A.L., Lambert, J.P., St-Denis, N.A., Li, T., Miteva, Y.V., Hauri, S., Sardiu, M.E., Low, T.Y., Halim, V.A., Bagshaw, R.D., Hubner, N.C., Al-Hakim, A., Bouchard, A., Faubert, D., Fermin, D., Dunham, W.H., Goudreault, M., Lin, Z.Y., Badillo, B.G., Pawson, T., Durocher, D., Coulombe, B., Aebersold, R., Superti-Furga, G., Colinge, J., Heck, A.J., Choi, H., Gstaiger, M., Mohammed, S., Cristea, I.M., Bennett, K.L., Washburn, M.P., Raught, B., Ewing, R.M., Gingras, A.C., Nesvizhskii, A.I., 2013. The CRAPome: a contaminant repository for affinity purification-mass spectrometry data. Nat Methods 10, 730-736.
Michalak, W., Tsiamis, V., Schwammle, V., Rogowska-Wrzesinska, A., 2019. ComplexBrowser: A Tool for Identification and Quantification of Protein Complexes in Large-scale Proteomics Datasets. Mol Cell Proteomics 18, 2324-2334.
Miles, W.O., Jaffray, E., Campbell, S.G., Takeda, S., Bayston, L.J., Basu, S.P., Li, M., Raftery, L.A., Ashe, M.P., Hay, R.T., Ashe, H.L., 2008. Medea SUMOylation restricts the signaling range of the Dpp morphogen in the Drosophila embryo. Genes Dev 22, 2578-2590.
Mitchelson, A., Simonelig, M., Williams, C., O'Hare, K., 1993. Homology with Saccharomyces cerevisiae RNA14 suggests that phenotypic suppression in Drosophila melanogaster by suppressor of forked occurs at the level of RNA stability. Genes Dev 7, 241-249.
Mohan, M., Bartkuhn, M., Herold, M., Philippen, A., Heinl, N., Bardenhagen, I., Leers, J., White, R.A., Renkawitz-Pohl, R., Saumweber, H., Renkawitz, R., 2007. The Drosophila insulator proteins CTCF and CP190 link enhancer blocking to body patterning. EMBO J 26, 4203-4214.
Mohrmann, L., Langenberg, K., Krijgsveld, J., Kal, A.J., Heck, A.J., Verrijzer, C.P., 2004. Differential targeting of two distinct SWI/SNF-related Drosophila chromatin-remodeling complexes. Mol Cell Biol 24, 3077-3088.
Mount, S.M., Salz, H.K., 2000. Pre-messenger RNA processing factors in the Drosophila genome. J Cell Biol 150, F37-44.
Nibu, Y., Senger, K., Levine, M., 2003. CtBP-independent repression in the Drosophila embryo. Mol Cell Biol 23, 3990-3999.
Nilkanta, C., Bagchi, A., 2018. Comparative analysis of prokaryotic and eukaryotic transcription factors using machine-learning techniques. Bioinformation 14, 315-326.
Nir, R., Grossman, R., Paroush, Z., Volk, T., 2012. Phosphorylation of the Drosophila melanogaster RNA-binding protein HOW by MAPK/ERK enhances its dimerization and activity. PLoS Genet 8, e1002632.
Ogirenko, A.A., Karagodin, D.A., Pavlova, N.V., Fedorova, S.A., Voloshina, M.A., Baricheva, E.M., 2008. [Molecular and genetic description of a new hypomorphic mutation of Trithorax-like gene and analysis of its effect on Drosophila melanogaster oogenesis]. Ontogenez 39, 134-142.
Papoulas, O., Daubresse, G., Armstrong, J.A., Jin, J., Scott, M.P., Tamkun, J.W., 2001. The HMG-domain protein BAP111 is important for the function of the BRM chromatin-remodeling complex in vivo. Proc Natl Acad Sci U S A 98, 5728-5733.
Park, J.W., Parisky, K., Celotto, A.M., Reenan, R.A., Graveley, B.R., 2004. Identification of alternative splicing regulators by RNA interference in Drosophila. Proc Natl Acad Sci U S A 101, 15974-15979.
Parker, D.S., Ni, Y.Y., Chang, J.L., Li, J., Cadigan, K.M., 2008. Wingless signaling induces widespread chromatin remodeling of target loci. Mol Cell Biol 28, 1815-1828.
Parkhurst, S.M., Corces, V.G., 1986. Interactions among the gypsy transposable element and the yellow and the suppressor of hairy-wing loci in Drosophila melanogaster. Mol Cell Biol 6, 47-53.
Perez-Lluch, S., Blanco, E., Carbonell, A., Raha, D., Snyder, M., Serras, F., Corominas, M., 2011. Genome-wide chromatin occupancy analysis reveals a role for ASH2 in transcriptional pausing. Nucleic Acids Res 39, 4628-4639.
Phillips, J.W., Pan, Y., Tsai, B.L., Xie, Z., Demirdjian, L., Xiao, W., Yang, H.T., Zhang, Y., Lin, C.H., Cheng, D., Hu, Q., Liu, S., Black, D.L., Witte, O.N., Xing, Y., 2020. Pathway-guided analysis identifies Myc-dependent alternative pre-mRNA splicing in aggressive prostate cancers. Proc Natl Acad Sci U S A 117, 5269-5279.
Phizicky, E.M., Fields, S., 1995. Protein-protein interactions: methods for detection and analysis. Microbiol Rev 59, 94-123.
Pyrowolakis, G., Hartmann, B., Muller, B., Basler, K., Affolter, M., 2004. A simple molecular complex mediates widespread BMP-induced repression during Drosophila development. Dev Cell 7, 229-240.
Riese, J., Yu, X., Munnerlyn, A., Eresh, S., Hsu, S.C., Grosschedl, R., Bienz, M., 1997. LEF-1, a nuclear factor coordinating signaling inputs from wingless and decapentaplegic. Cell 88, 777-787.
Rospert, S., Dubaquie, Y., Gautschi, M., 2002. Nascent-polypeptide-associated complex. Cell Mol Life Sci 59, 1632-1639.
Schneider, M., Khalil, A.A., Poulton, J., Castillejo-Lopez, C., Egger-Adam, D., Wodarz, A., Deng, W.M., Baumgartner, S., 2006. Perlecan and Dystroglycan act at the basal side of the Drosophila follicular epithelium to maintain epithelial organization. Development 133, 3805-3815.
Schweizer, L., Nellen, D., Basler, K., 2003. Requirement for Pangolin/dTCF in Drosophila Wingless signaling. Proc Natl Acad Sci U S A 100, 5846-5851.

Sekelsky, J.J., Newfeld, S.J., Raftery, L.A., Chartoff, E.H., Gelbart, W.M., 1995. Genetic characterization and cloning of mothers against dpp, a gene required for decapentaplegic function in Drosophila melanogaster. Genetics 139, 1347-1358.

Shchepinov, M.S., Udalova, I.A., Bridgman, A.J., Southern, E.M., 1997. Oligonucleotide dendrimers: synthesis and use as polylabelled DNA probes. Nucleic Acids Res 25, 4447-4454.

Sierra, J., Yoshida, T., Joazeiro, C.A., Jones, K.A., 2006. The APC tumor suppressor counteracts beta-catenin activation and H3K4 methylation at Wnt target genes. Genes Dev 20, 586-600.

Sinenko, S.A., Hung, T., Moroz, T., Tran, Q.M., Sidhu, S., Cheney, M.D., Speck, N.A., Banerjee, U., 2010. Genetic manipulation of AML1-ETO-induced expansion of hematopoietic precursors in a Drosophila model. Blood 116, 4612-4620.

Smibert, C.A., Lie, Y.S., Shillinglaw, W., Henzel, W.J., Macdonald, P.M., 1999. Smaug, a novel and conserved protein, contributes to repression of nanos mRNA translation in vitro. RNA 5, 1535-1547.

Somma, M.P., Fasulo, B., Cenci, G., Cundari, E., Gatti, M., 2002. Molecular dissection of cytokinesis by RNA interference in Drosophila cultured cells. Mol Biol Cell 13, 2448-2460.

Song, H., Hasson, P., Paroush, Z., Courey, A.J., 2004. Groucho oligomerization is required for repression in vivo. Mol Cell Biol 24, 4341-4350.

Spitz, F., Furlong, E.E., 2012. Transcription factors: from enhancer binding to developmental control. Nat Rev Genet 13, 613-626.

Symes, A.J., Eilertsen, M., Millar, M., Nariculam, J., Freeman, A., Notara, M., Feneley, M.R., Patel, H.R., Masters, J.R., Ahmed, A., 2013. Quantitative analysis of BTF3, HINT1, NDRG1 and ODC1 protein over-expression in human prostate cancer tissue. PLoS One 8, e84295.


Symes, A.J., Eilertsen, M., Millar, M., Nariculam, J., Freeman, A., Notara, M., Feneley, M.R., Patel, H.R., Masters, J.R., Ahmed, A., 2013. Quantitative analysis of BTF3, HINT1, NDRG1 and ODC1 protein over-expression in human prostate cancer tissue. PLoS One 8, e84295.


Takahashi, S., Takada, I., 2023. Recent advances in prostate cancer: WNT signaling, chromatin regulation, and transcriptional coregulators. Asian J Androl 25, 158-165.

Terriente-Felix, A., de Celis, J.F., 2009. Osa, a subunit of the BAP chromatin-remodelling complex, participates in the regulation of gene expression in response to EGFR signalling in the Drosophila wing. Dev Biol 329, 350-361.

Toledano-Katchalski, H., Nir, R., Volohonsky, G., Volk, T., 2007. Post-transcriptional repression of the Drosophila midkine and pleiotrophin homolog miple by HOW is essential for correct mesoderm spreading. Development 134, 3473-3481.

Tyler, J.K., Bulger, M., Kamakaka, R.T., Kobayashi, R., Kadonaga, J.T., 1996. The p55 subunit of Drosophila chromatin assembly factor 1 is homologous to a histone deacetylase-associated protein. Mol Cell Biol 16, 6149-6159.

Verni, F., Gandhi, R., Goldberg, M.L., Gatti, M., 2000. Genetic and molecular analysis of wings apart-like (wapl), a gene controlling heterochromatin organization in Drosophila melanogaster. Genetics 154, 1693-1710.

Volohonsky, G., Edenfeld, G., Klambt, C., Volk, T., 2007. Muscle-dependent maturation of tendon cells is induced by post-transcriptional regulation of stripeA. Development 134, 347-356.

Wagner, C.R., Hamana, K., Elgin, S.C., 1992. A high-mobility-group protein and its cDNAs from Drosophila melanogaster. Mol Cell Biol 12, 1915-1923.

Walsh, E.P., Brown, N.H., 1998. A screen to identify Drosophila genes required for integrin-mediated adhesion. Genetics 150, 791-805.

Wamsley, J.J., Issaeva, N., An, H., Lu, X., Donehower, L.A., Yarbrough, W.G., 2017. LZAP is a novel Wip1 binding partner and positive regulator of its phosphatase activity in vitro. Cell Cycle 16, 213-223.

Wickens, M., Bernstein, D.S., Kimble, J., Parker, R., 2002. A PUF family portrait: 3'UTR regulation as a way of life. Trends Genet 18, 150-157.

Yan, D., Perrimon, N., 2015. spenito is required for sex determination in Drosophila melanogaster. Proc Natl Acad Sci U S A 112, 11606-11611.

Yao, J., Ardehali, M.B., Fecko, C.J., Webb, W.W., Lis, J.T., 2007. Intranuclear distribution and local dynamics of RNA polymerase II during transcription activation. Mol Cell 28, 978-990.

Yeom, E., Hong, S.T., Choi, K.W., 2015. Crumbs interacts with Xpd for nuclear division control in Drosophila. Oncogene 34, 2777-2789.

Yu, A., Rual, J.F., Tamai, K., Harada, Y., Vidal, M., He, X., Kirchhausen, T., 2007. Association of Dishevelled with the clathrin AP-2 adaptor is required for Frizzled endocytosis and planar cell polarity signaling. Dev Cell 12, 129-141.

Zaessinger, S., Busseau, I., Simonelig, M., 2006. Oskar allows nanos mRNA translation in Drosophila embryos by preventing its deadenylation by Smaug/CCR4. Development 133, 4573-4583.

Zehring, W.A., Lee, J.M., Weeks, J.R., Jokerst, R.S., Greenleaf, A.L., 1988. The C-terminal repeat domain of RNA polymerase II largest subunit is essential in vivo but is not required for accurate transcription initiation in vitro. Proc Natl Acad Sci U S A 85, 3698-3702.
Zeng, Y.A., Rahnama, M., Wang, S., Lee, W., Verheyen, E.M., 2008. Inhibition of Drosophila Wg signaling involves competition between Mad and Armadillo/beta-catenin for dTcf binding. PLoS One 3, e3893.
Zhang, P., Wu, Y., Belenkaya, T.Y., Lin, X., 2011. SNX3 controls Wingless/Wnt secretion through regulating retromer-dependent recycling of Wntless. Cell Res 21, 1677-1690.
