## Supplementary_Figures_Legends for "Functional dissection of *Drosophila Myc cis*-regulatory modules (Myc-CRMs) reveals developmentally active DNA-protein interactions"

### Supplementary Figures and Legends

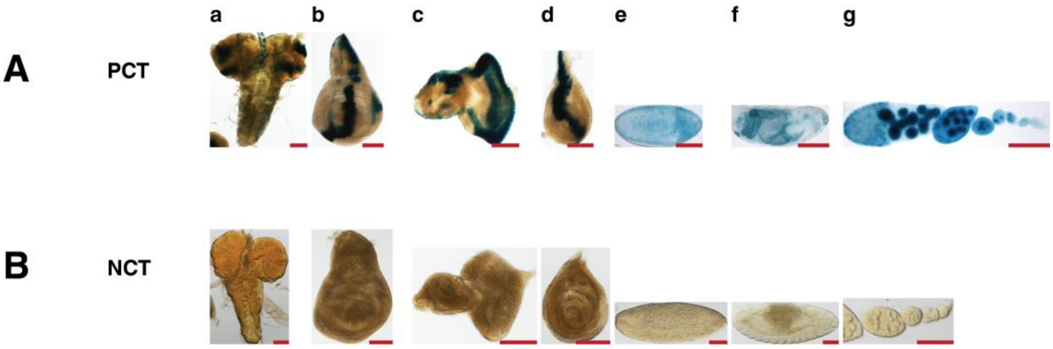


**Fig. S1. β-Galactosidase staining of positive and negative controls.**
Positive control (**A, PCT**) used tissues from 3rd instar larvae and adult females of the *dpp-lacZ* strain, and negative control (**B, NCT**) used *y[1] w[1118]* flies. **Abbreviations**: **a**, brain; **b**, wing disc; **c**, eye-antennal disc, **d**, leg disc; **e,f**, embryos, **g**; ovary. Scale bar: 100 µm.

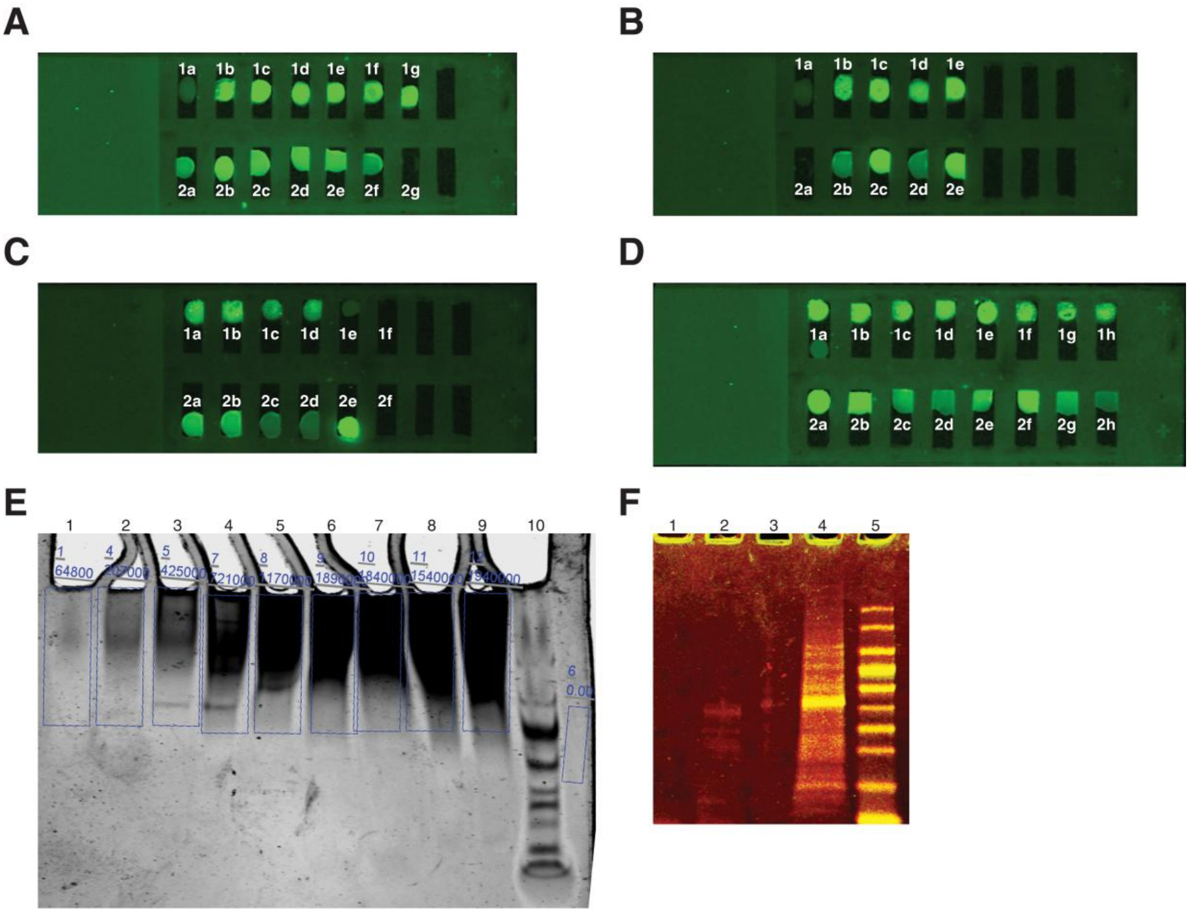


**Fig. S2. Assessment of Magnetic Beads, Bead-PEG23-DNA constructs, and Soluble Nuclear Fraction (SNF) prior to protein identification.**
**(A-D)** Visualization of Bead-PEG23-DNA (IR-labeled) constructs on glass slides and corresponding supernatants before protein binging:
**(A) 1a**: Raw beads; Pos1/Pos2 (**1b-1c**: construct; **2a-2b**: supernatant); P1/P2 (**1d-1g**: construct; **c-f**: supernatant); **2g**: binding buffer (input 3 µL of 0.3 pmol/300 µL; supernatants/buffer 30 µL; raw beads 30 µL).
**(B) 1a**: Raw beads; Scm1/Scm2 (**1b-1e**: construct; **2b-2e**: supernatant); **2a**: buffer; (same input/supernatant beads volumes).
**(C)** P29/P30 (**1a-1d**: construct; **2a-2d**: supernatant); **1e**: raw beads, and **1f**: buffer; (same input/supernatant beads volumes); **2e**:IR5-P29/P30-IR3 only (0.3 pmol); **2f**: buffer.

**(D) 1a bottom**: raw beads; P31/P32 (**1a top & 1b-1d**: construct; **2a-2d**: supernatant); and P35/P36 (**1e-1h**: constructs; **2e-2h**: supernatants) (same input/supernatant beads volumes).
**(E)** Lanes **1-9**: Quality and quantity check of SNF (10-100 µg) under non-denaturing conditions (0.5% TBE, 5% precast PAGE), lane **10**: protein ladder.
**(F)** Magnetic bead contamination test: lanes **1-2** (100 µg beads), Laemmli ± 8M urea; lane **4**, 50 µg SNF; lane **5**, protein ladder. Artifactual bands (~15-40 kDa) were weak; the only unique contaminant from blank beads was collagen alpha I(III) chain (P04258, 1466 residues, 138,4 kDa).
All electrophoresis was run on 5% SDS-PAGE at 90 V for 50 min.

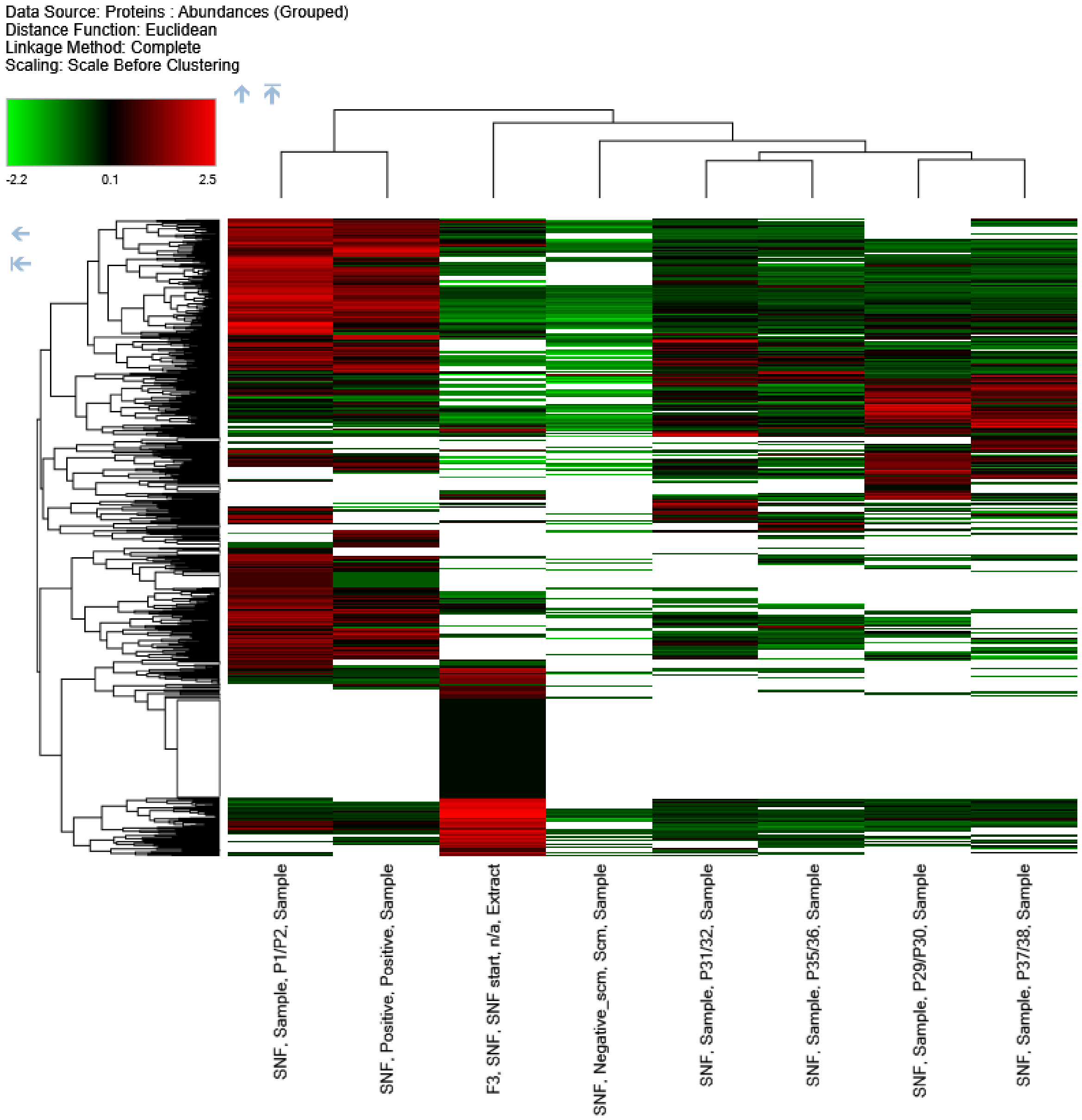


**Fig. S3. Heat map of identified proteins using hierarchical clustering.**
Log2 label-free ratios of 1001 quality-filtered proteins from *Drosophila* embryonic nuclear extracts (0-72 h AEL) were analyzed. Proteins were filtered for "SNF ≥10x Scm" to minimize false detection (FDR) and improve confidence. Each sample included two replicates with either the "top" or "bottom" strand of double-stranded oligo attached to the Bead-PEG23 surface, ensuring retention of proteins associated with both strands.
Oligo pairs are grouped, and functionally similar and related groups cluster together, reference**:** Key, M. (2012) (e.g., Bead-PEG23-IR-P1/P2 with P2/P1). Overall patterns show P1/P2 & Pos1/Pos2, P31/P32 & P35/p36, and P29/P30 & P37/P38 as relative groups, reflecting similar expression patterns and biological functions: P1/P2 & Pos1/Pos2 predominantly as tissue-specific enhancers, P31/P32 & P35/p36 as promoters, and P29/P30 & P37/P38 as strong developmental enhancers.

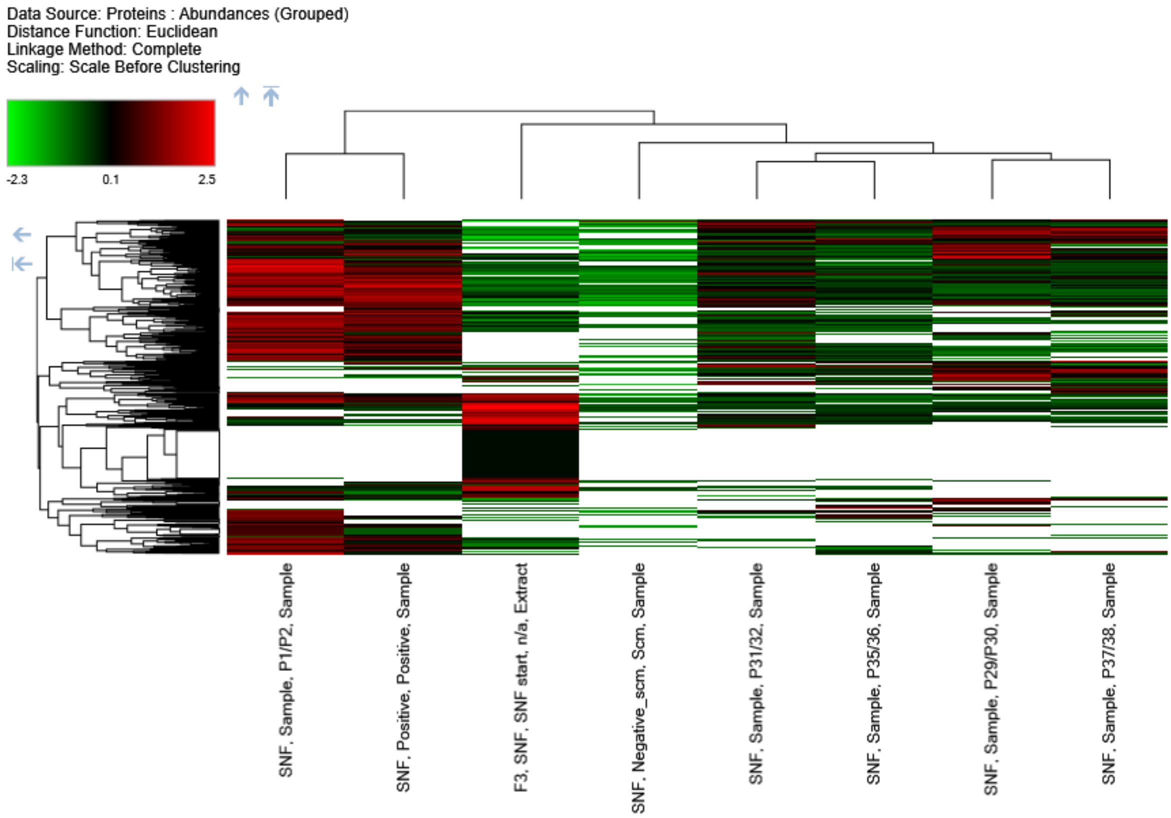


**Fig. S4. Heat map of proteomics data after applying the "SNF ≥2x Scm" quality filter.**
Proteins were filtered so that the mass-to-charge ratio in each sample was at least 2x higher than in the Scrambled negative control (Scm). Functionally similar and related groups cluster together, and sample pairs are grouped similarly to Figure S3. A tutorial in displaying mass spectrometry-based proteomics data using heat maps (Binz et al., 2012).

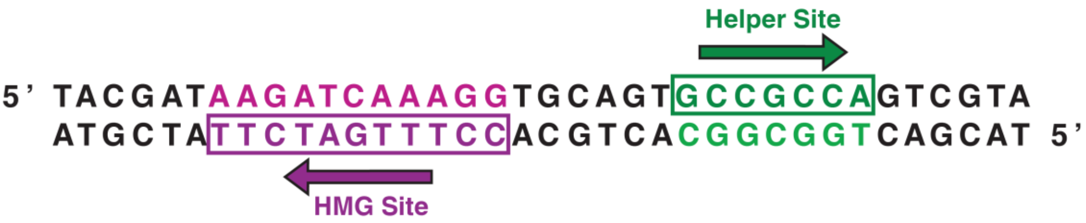


**Fig. S5. W-CRM variant with HMG and Helper sites as a positive control for protein identification.**
The HMG/Helper site pair is arranged in the Akimbo (AK) orientation with six-base spacing (AK6). Arrows indicate the 5′→3′ direction on each strand. HMG (magenta) and Helper (green) sequences correspond to sites used in DNA-binding and synthetic Wnt reporter assays. Figure adapted from Archbold et al. (2014).

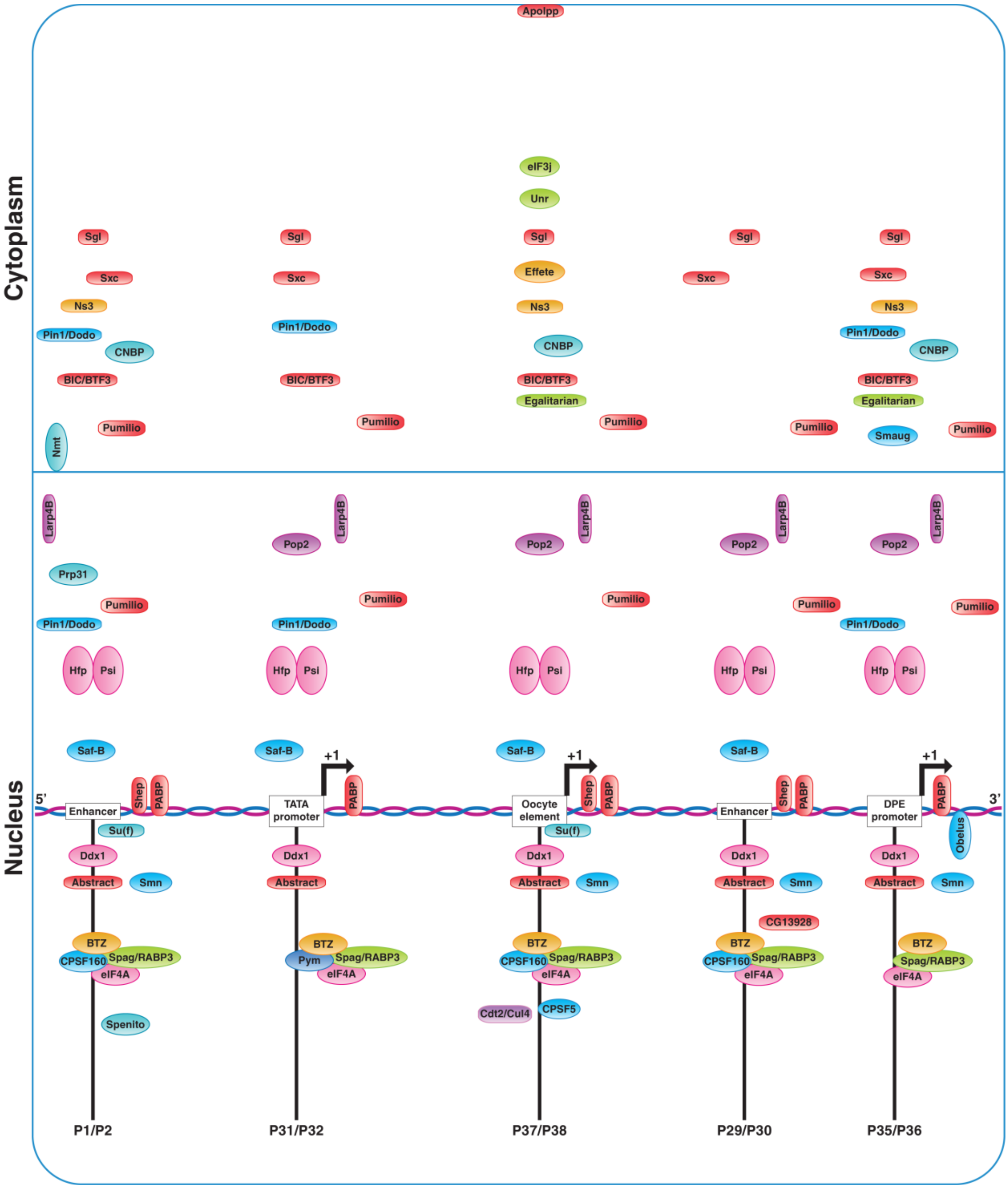


**Fig. S6. Examples of CRM-associated post-transcriptional regulatory network (RNA processing / stability / translation).**
Unr, an RNA chaperone promoting *Myc* mRNA translation, was exclusively associated with the Oocyte Element (P37/P38) with low abundance (and absent from nuclear extract input; Table S6). CNBP, a conserved CCHC-type zinc finger protein, promotes *Myc* translation and associates with the P1/P2, P35/P36, and P37/P38. Core Exon Junction Complex components—Barentsz (Btz), eIF4A, Ddx1, Pym, and Spaghetti (Spag)—coordinate splicing, export, and mRNA stability. Cleavage and polyadenylation factors CPSF160 and CPSF5 were linked to P1/P2, P29/P30, and P37/P38 (CPSF5 only at the P37/P38), with Poly(A) binding protein (PABP) present at all Myc-CRMs. Additional RNA regulatory proteins at all Myc-CRMs included Half pint (Hfp), Psi, and Larp4B.
See Tables S3 and S6 for associations and abundance ratios of all the Myc-CRMs-associated RNA regulatory factors.
