## Supplementary material for "Functional dissection of *Drosophila Myc cis*-regulatory modules (Myc-CRMs) reveals developmentally active DNA-protein interactions": README_20260905

**Organization of Supplementary Data accompanying the manuscript**

This archive contains main manuscript, supplementary figures, tables, datasets, and analysis files referenced in the main text. All files are organized according to their corresponding figure and table numbers.

**Main Manuscript**

**Manuscript_20260904.docx** contains the main manuscript, including the title page, abstract, main text, figure and table callouts, figure legends, references, and citations to the accompanying supplementary materials.

**Supplementary Tables**

**Supplementary Table S1** is provided as a DOCX file and contains resource and methodological information, including plasmid descriptions, PCR and sequencing primers, *Drosophila* stocks, EMSA oligonucleotides (Myc-CRMs; cis-regulatory modules) with annealing schemes (oligo pairs), and proteomics sample names (Tables S1a–S1g).

**Supplementary Data files S2–S7 and S9–**S11 are provided as Excel files and contain Supplementary Tables S2–S7 and S9–S11, respectively. Supplementary Data files S2–S7 additionally include the corresponding Supplementary Figures S7–S12 illustrating the filtering and quality control (QC) workflow.

**Supplementary Tables S8, S12, and S25** are provided as separate standalone files. Table S8 lists the protein complexes associated with the identified factors. Table S12 contains DNA-binding proteins together with their gene IDs, UniProt accession numbers, and protein abundance ratios. Table S25 is derived from Table S7, from which ribosomal, mitochondrial, and most cytosolic proteins have been removed to focus on prioritizing candidates for direct chromatin and transcriptional regulation.

**Supplementary_Data_S13_S18** is provided as a folder containing six Excel files corresponding to Supplementary Tables S13–S18. These files contain the proteomics-derived protein lists for six volcano plot analyses, including UniProt accession numbers and q-values for the tested Myc-CRMs and the positive control W-CRMs.

**Supplementary Methods, Results, and References**

**Supplementary_Methods_Results_Refs.pdf** contains Supplementary Methods (S1–S9), Supplementary Results (S3.1–S3.7 and S3.10–S3.14), and Supplementary References. It includes detailed descriptions of materials and reagents, experimental protocols, molecular cloning, EMSA, Proteomics, data analyses, and detailed description of proteins associated with Myc-CRMs.

**Supplementary Figures and Legends**

**Supplementary_Figures_Legends.pdf** contains Supplementary Figures S1–S6 together with their corresponding figure legends.
